# An internal scaffold supports microtubule architectural diversity in *Xenopus* spermatids

**DOI:** 10.64898/2026.08.28.747525

**Authors:** Céline Callens, Matthieu P.M.H. Benoit, Florian Berger, Quentin Rouger, Roselyne Viel, Claire Heichette, Charlotte Guyomar, Laurence Duchesne, Blandine Guével, Régis Lavigne, Emmanuelle Com, Charles Pineau, Kévin Macé, Jérôme Jullien, Denis Chrétien, Romain Gibeaux

**Author notes:** Equal contributions (Céline Callens, Matthieu P.M.H. Benoit, Florian Berger, Quentin Rouger, Denis Chrétien, Romain Gibeaux).

## Abstract

During spermiogenesis, early round spermatids differentiate into specialized spermatozoa through an extensive reshaping of the nucleus driven by coordinated cytoskeletal and chromatin-based mechanisms. In mammals, this process critically relies on the transient manchette, a microtubule-based structure that remodels the spermatid nucleus and serves a track to transport material required for flagellum assembly. However, the existence, organization, and molecular composition of such a structure in other vertebrates have remained poorly investigated. Here, we establish that an organized microtubule network is present in *Xenopus* spermatids and shares key architectural and molecular hallmarks of the mammalian manchette. We further uncover a large structural heterogeneity of spermatid microtubules, with variable protofilament numbers, skew angles, and lattice organizations. We reveal the presence of a Spaca9–Saxo2 internal scaffold in spermatid microtubules, suggesting an internal reinforcement mechanism necessary for extensive nuclear reshaping and cytoplasm remodeling.

## Main

During vertebrate spermatogenesis, immature diploid spermatogonia undergo successive mitotic and meiotic divisions leading to haploid spermatids. Although spermatids are the end product of meiosis, these cells are round at their early stage, transcriptionally active, and incapable of fertilization. Through spermiogenesis, they undergo a complex differentiation during which the cytoplasm is remodeled, the flagellum is assembled and the nucleus is reshaped to form the sperm head. Nuclear remodeling relies on diverse cytoskeletal and chromatin-based mechanisms, whose relative contributions vary across species. These include endogenous forces generated by acrosome-acroplaxome-dependent processes and by the manchette, chromatin condensation or coiling, as well as exogenous forces exerted by Sertoli cells ^1–5^.

In mammals, the spermatid manchette is a transient structure surrounding the nucleus and composed of actin and packed, largely parallel microtubules, which can number up to ∼1,000 in rodents ^6^. Beyond its role in nuclear shaping ^7^, it also serves as a transport platform that delivers components required for flagellum assembly ^8^. The manchette has been particularly well characterized in murine models, giving rise to our current view of its organization and function. It forms around the distal half of the nucleus during the later stages of spermiogenesis and disassembles by the end of it. During assembly, microtubules polymerize around the nucleus, their plus-ends are captured by the perinuclear ring composed of δ-tubulin and keratin, and they are finally organized into a linear array of cross-linked parallel microtubules ^9–11^. While δ-tubulin is a component of the perinuclear ring, both δ-and ε-tubulin localize along manchette microtubules ^10,11^, where, by analogy to their stabilizing role at centrioles, they have been proposed to contribute to manchette microtubule stability ^12^. In addition, microtubule polarity within the manchette has been inferred from the presence of EB3 at the perinuclear ring and γ-tubulin at the opposite side of the manchette ^13,14^. Once assembled, the rigid manchette exerts forces that progressively sculpt the spermatid nucleus ^7^ by moving caudally while engaging with the nuclear envelope ^15^. This movement is regulated by a wide range of proteins such as dynein ^16^, the katanin catalytic subunit A1-like 2 (KATNAL2) and regulatory subunit B1 (KATNB1) ^11,17^, as well as the cargo adaptor HOOK1 ^18^.

Despite this detailed understanding in mammals, the occurrence, organization and function of such a structure in other vertebrates remain unclear. This is the case of *Xenopus* species, whose sperm exhibit one of the most extreme nuclear shape transformations among vertebrates, producing a characteristically thin corkscrew-shaped head ^19^. This makes *Xenopus* an appealing system in which to explore cytoskeletal elements of nuclear reshaping. Here, we investigated *Xenopus* spermatid microtubules. We found that a dense microtubule network surrounds the spermatid nucleus and reorganizes into a skirt-like structure at its caudal side, which we characterized as having manchette-like characteristics, a structure thought to be absent in *Xenopus* ^20,21^. We uncovered a substantial diversity of microtubule lattice organizations within this spermatid structure, not reported in other cell types or *in vitro*. We moreover identified an internal scaffold within *Xenopus* spermatid microtubules, comprising the Spaca9 and Saxo2 proteins. We also reveal a complex internal membrane organization within which several of these microtubules are embedded. Altogether, this study provides a detailed molecular analysis of spermatid microtubules of non-mammalian species and their internal components within a manchette-like structure, and further suggests a role for this scaffold in reinforcing microtubules to sustain forces required for nuclear reshaping and cytoplasm reorganization.

## Results

### *Xenopus laevis* spermatids harbor a microtubule network with manchette-like characteristics

We first analyzed *Xenopus laevis* germ cells *in situ*. In *Xenopus* seminiferous tubules, germ cells are organized into cysts of clonal populations embedded within Sertoli cells (Fig. 1a). This cyst organization allowed us to distinguish germ cell types, as previously described ^22^. Using histochemistry and DAPI labeling on testis sections, we measured their nuclear sizes (Fig. 1a, left and middle panels). Based on these measurements, *Xenopus laevis* primary spermatocytes, secondary spermatocytes and spermatids were characterized as cells with a mean nuclear diameter of 11.2 ± 1.2 μm (average ± standard deviation), 8.2 ± 1.1 μm, and 5.7 ± 0.7 μm, respectively (Fig. 1a, right panel). Analysis of whole-testis cell suspensions showed a large fraction of sperm cells, representing 88.7 ± 7.5% of the total cell count (Fig. 1b, left and middle panels). To enrich for spermatids, we fractionated the cell populations using a density gradient as previously described ^23^, which reduced the sperm fraction to 25.9 ± 13.7% of the resulting suspension. Among the remaining 74.1 ± 13.7% of germ cells, the mixed-population distribution was modeled as a weighted mixture of the three germ cell type reference distributions, which allowed us to estimate that the suspension contained 1.4 ± 1.1% primary spermatocytes, 42.5 ± 3.6% secondary spermatocytes, and 56.1 ± 3.4% spermatids (Methods; Fig. 1b, rightmost panel). This spermatid-enriched cell suspension was then analyzed by immunofluorescence, and spermatids were categorized from "early" to "late" based on nuclear size and shape (Fig. 1c). Using this characterization, we determined that the spermatid population was composed of 91 ± 3.4% early, 7.8 ± 4.5% mid, and 1.2 ± 1.3% late spermatids. At the early stage, microtubules form a peripheral network around the nucleus (Fig. 1c, left panel) and progressively reorganize into an asymmetric microtubule array (Fig. 1c, middle panel). As the nucleus changes shape, decreases in size, and adopts a pear-like morphology from mid to late stages, a polarized skirt-like microtubule structure becomes apparent at the caudal side of the nucleus (Fig. 1c, right panel), reminiscent of the mammalian manchette. This structure is then disassembled and absent in mature sperm cells.

**Fig. 1:**
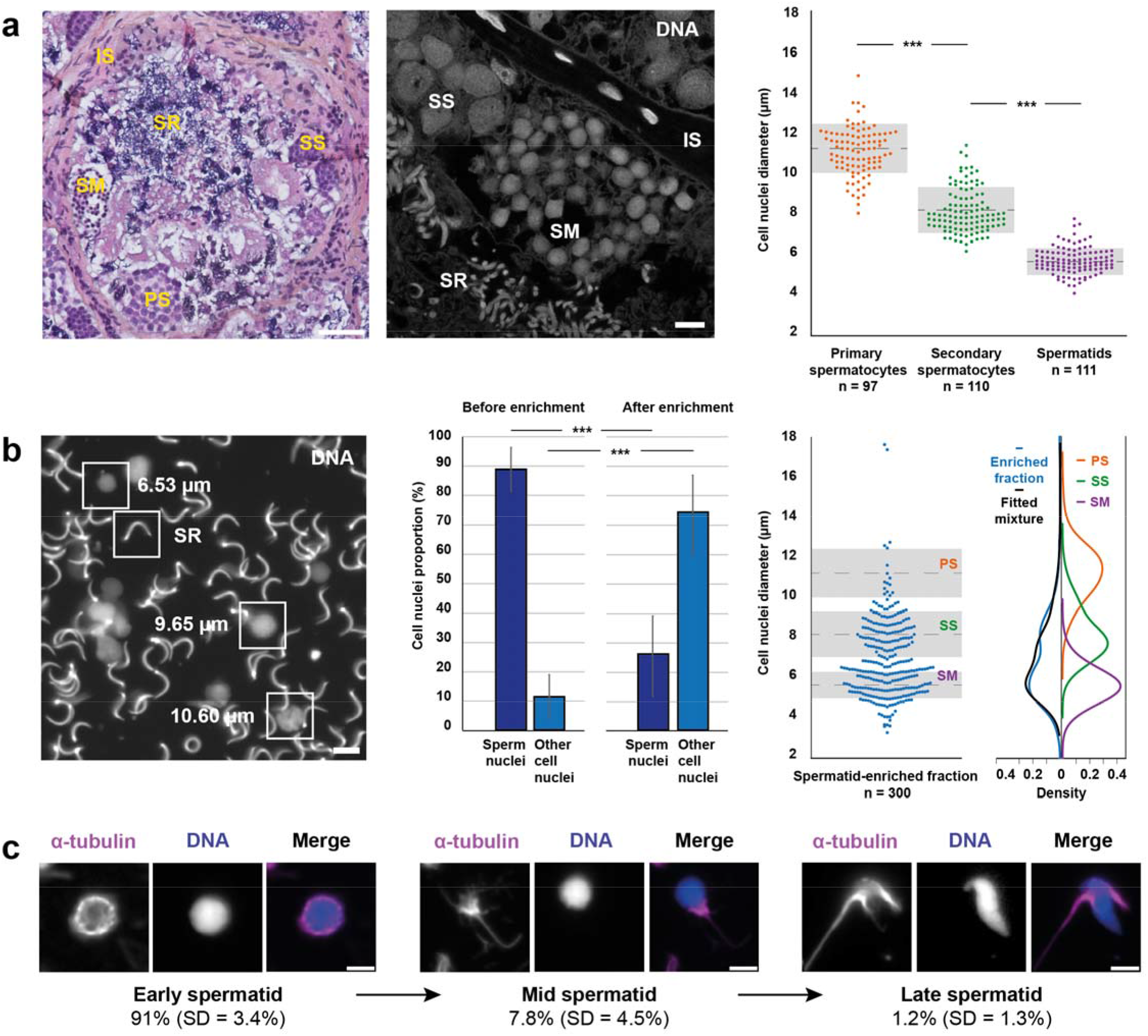
*Xenopus laevis* spermatid characterization. **a,** Representative testis section stained with hematoxylin and eosin (left panel). Intertubular space (IS), primary spermatocytes (PS), secondary spermatocytes (SS), spermatids (SM), and sperm cells (SR), are labeled. Scale bar: 50 µm. Cysts of germ cells visualized *in situ* by nuclear staining using Hoechst (middle panel). Scale bar: 10 µm. Nuclear diameters of primary spermatocytes, secondary spermatocytes and spermatids, with a mean of 11.2 ± 1.2 µm, 8.2 ± 1.1 µm and 5.7 ± 0.7 µm, respectively (right panel). Dashed lines indicate the mean and the gray boxes the standard deviation. **b,** Image of testicular whole cell suspension stained with DAPI (left panel), nuclei with representative sizes of the different types of germ cells are boxed. Scale bar: 10 µm. Quantification of sperm nuclei percentage before enrichment in other germ cell types (88.7 ± 7.5%), and after enrichment (25.9 ± 13.7%) (middle panel; mean of three independent experiments; total of 3,357 nuclei). Measurement of nuclear diameters in the spermatid-enriched fraction (right panel). Dashed lines indicate the mean and the gray boxes the standard deviation from panel (a) of primary spermatocytes, secondary spermatocytes, and spermatids, labeled as PS, SS, and SM, respectively. The nuclear size distribution of the spermatid enriched fraction was fitted as a weighted mixture of the kernel density distributions of the three reference cell types (PS, SS and SM). The summed fitted distribution closely matched the observed population density. **c,** Immunofluorescence images of isolated spermatids. Microtubules (magenta) and the spermatid nucleus (blue) are shown. Microtubule network and nuclear size and shape allowed classification into early, mid and late spermatid stages. Scale bar: 5 µm. Statistical significance analysis in a.-b. was performed using the Kruskal-Wallis test.

We further characterized this microtubule network by localizing known mammalian manchette markers from early to late spermatid stages (Fig. 2a-c, Supplementary Fig. 1). Actin localizes at the anterior side of the spermatid nucleus during the early stages of spermiogenesis, and progressively shifts toward the caudal region (Supplementary Fig. 1a), consistent with the formation of the acrosome-acroplaxome complex, of which F-actin is a known component in rodents ^4^. At mid-spermatid stage, actin colocalizes with the polarized microtubule network, and this colocalization persists as the structure moves caudally, indicating the presence of an actin cytoskeleton integrated within the network (Fig. 2a, top panel), as described in mammalian manchettes. We next examined δ-tubulin, a known marker of the manchette ^24^. As spermiogenesis progresses, we observed δ-tubulin colocalizing with microtubules (Supplementary Fig. 1b), particularly with the skirt-like microtubule structure (Fig. 2a, bottom panel). Furthermore, we investigated the polarity of microtubules in this structure by assessing the localization of EB1 and γ-tubulin (Fig. 2b, Supplementary Fig. 1c,d). For antibody-compatibility reasons, microtubules were detected using an anti-polyglutamylation antibody. EB1 appeared as puncta along microtubules surrounding the nucleus, whereas γ-tubulin appeared more as a nucleation ring, with no clear dichotomy, suggesting a microtubule organization that may not be as polarized as described in mammalian manchettes ^13^. Next, we assessed microtubule post-translational modifications, including acetylation, polyglutamylation, and tyrosination (Fig. 2c). Microtubules within the polarized network were minimally acetylated, slightly less glutamylated and more tyrosinated than axonemal microtubules, in line with the fact that manchette and axoneme microtubules display distinct post-translational modification profiles 25_._

**Fig. 2:**
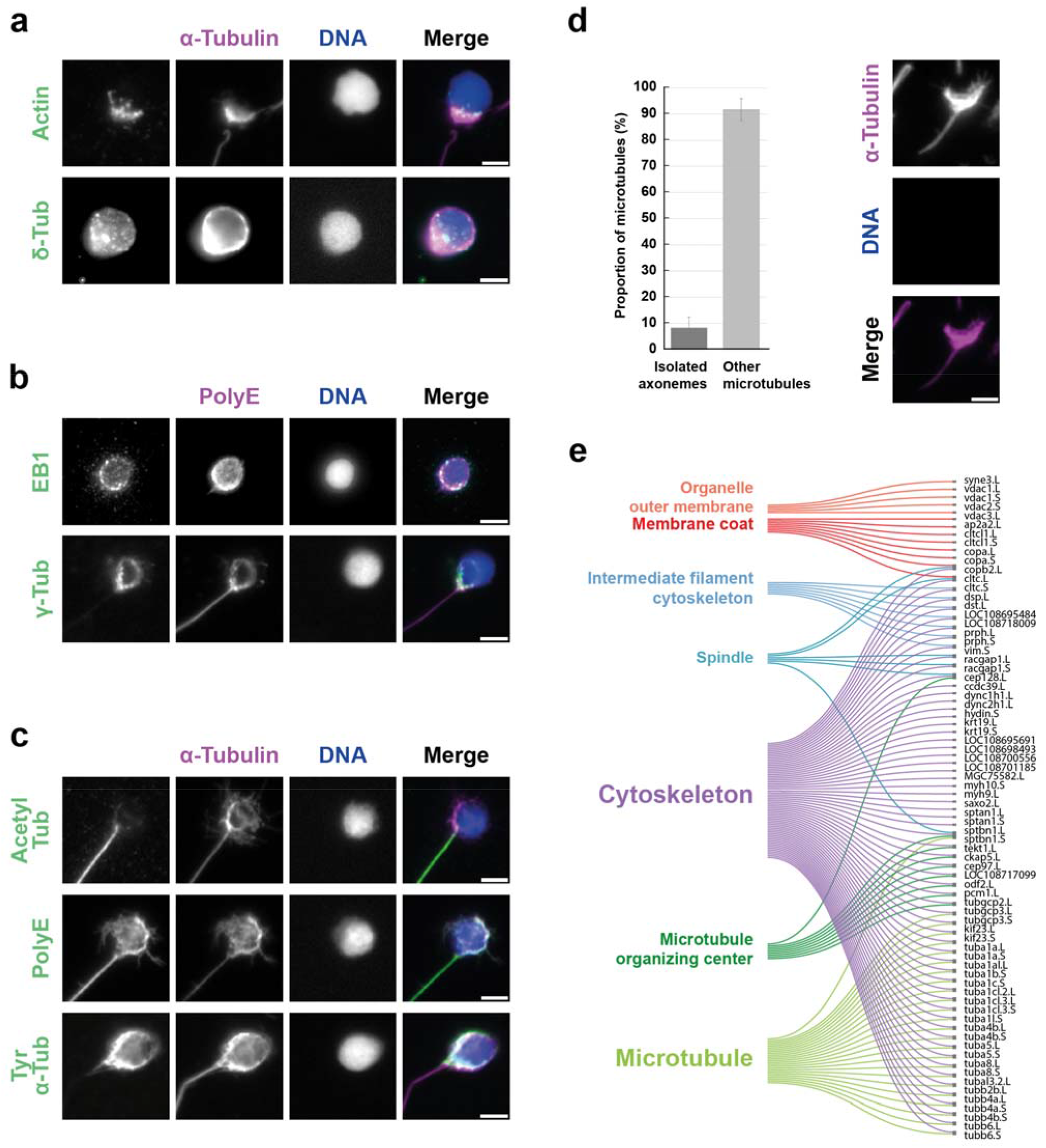
Molecular analysis of *Xenopus laevis* spermatid microtubule network. **a,** Immunofluorescence analysis of actin and δ-tubulin. **b,** Immunofluorescence analysis of EB1 and γ-tubulin. **c,** Immunofluorescence analysis of acetyl-tubulin, polyglutamylation, tyrosinated tubulin. **a-c,** Microtubules (magenta) and nuclei (blue) are shown with proteins of interest (green), scale bar: 5 µm. **d.** Percentage of microtubules from contaminating isolated axonemes and from other microtubule structures present in the spermatid microtubules-enriched fraction (left). Image of purified spermatid microtubules examined by immunofluorescence. Tubulin (magenta), DNA (blue; no signal was visible, demonstrating the absence of nucleus) are shown (right). Scale bar: 5 µm. **e,** Sankey diagram of all significantly enriched child GO terms following PANTHER overrepresentation test using GO-Slim Cellular Component terms (“microtubule”, “intermediate filament”, “membrane coat”, “organelle outer membrane”, “spindle” and “microtubule organizing center”). The terms “endocytic vesicle” and “clathrin-coated vesicle” fully overlapping with “membrane coat” are not displayed. The significantly enriched “cytoskeleton” parent term is shown for comparison.

We finally purified spermatid microtubules, taking advantage of a sucrose-gradient method originally developed for rat manchette isolation ^26^. Using immunofluorescence, we confirmed that the retrieved structures showed minimal contamination by axonemes (8.3 ± 4.1%) and were highly enriched in other microtubule structures (91.7 ± 4.1%) (Fig. 2d, left), detached from nuclei (Fig. 2d, right; note the absence of DNA signal). These preparations were therefore subjected to mass spectrometry proteomics. The 200 proteins with the highest scores were analyzed using a GO term overrepresentation test (Fig. 2e), which revealed significant enrichment in cytoskeletal component terms, including “microtubule”, “intermediate filament”, “microtubule organizing center”, consistent with manchette characteristics ^27^. The term “spindle” was also enriched, likely reflecting structural similarities between the manchette and the spindle. Notably, all genes from these four child terms also belonged to the overrepresented parent term “cytoskeleton”. In addition, “membrane coat” and “organelle outer membrane” child terms were also enriched, consistent with the presence of vesicles transported along microtubules as observed within manchettes ^28^. Taken together, our results reveal the existence of a polarized microtubule network in *Xenopus laevis* spermatids that shares several key features with mammalian manchette ^8,12^.

### *Xenopus laevis* spermatid microtubules contain an internal scaffold

To gain structural insights into the microtubules of this structure, intact isolated spermatids were first analyzed by cryo-electron microscopy (Fig. 3a, left panels). Although intact cells were too thick to be examined at high resolution, a dense microtubule network was visible at their periphery (Fig. 3a, middle panel). Strikingly, these microtubules displayed a striation repeating with an 8 nm periodicity (Fig. 3a, right panels).

**Fig. 3:**
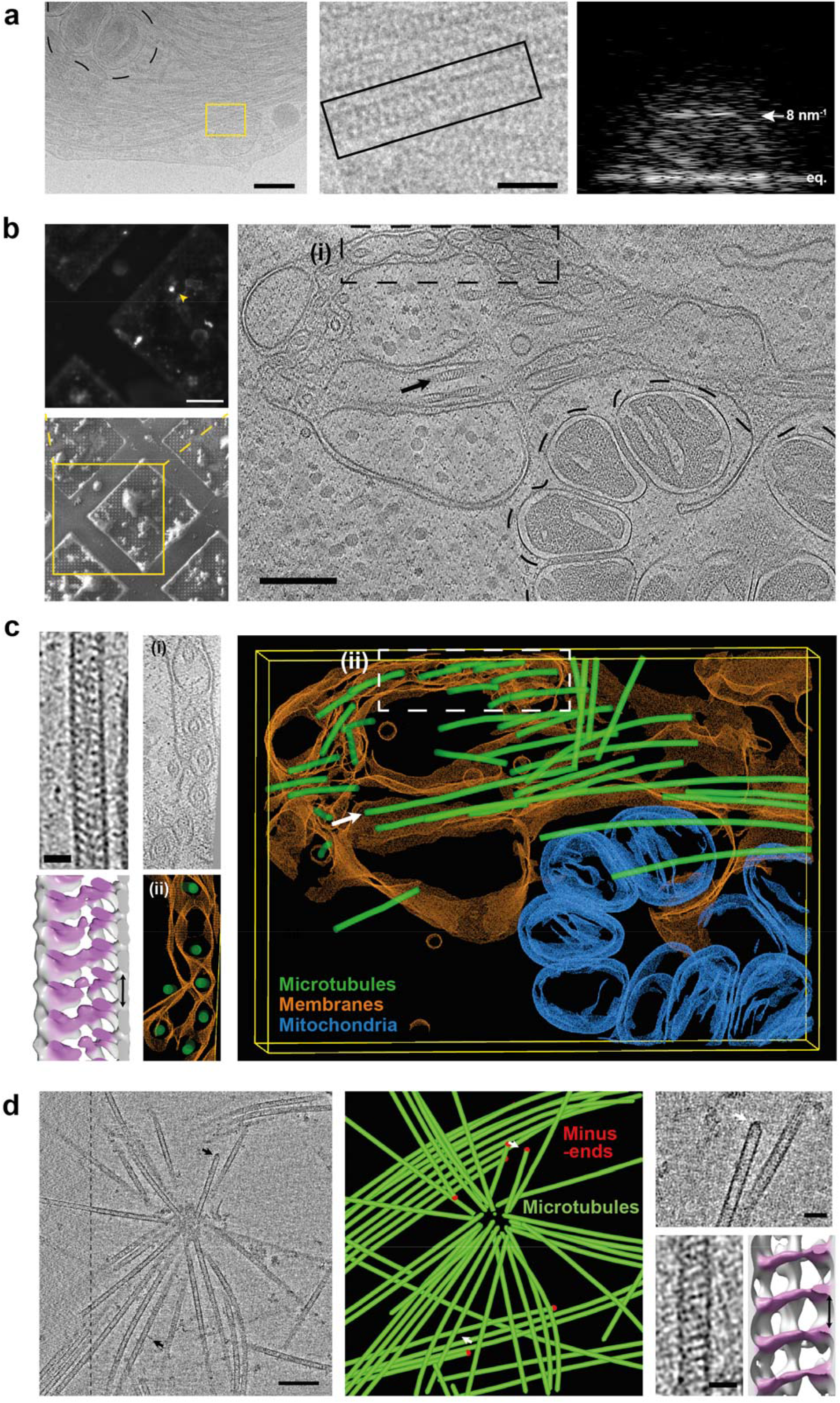
Structural characterization of *Xenopus laevis* spermatid microtubules.**a,** Cryo-electron microscopy images of an isolated spermatid showing a dense microtubule network within the cell, together with stacked mitochondria (left, black dashed contour). Microtubule internal striations are shown at higher magnification (middle), together with the Fourier transform of the boxed microtubule in the middle panel (right), showing a layer line at 8 nm^-1^ corresponding to the periodicity observed (eq. = equator line). Scale bars: 200 nm (left), 50 nm (middle). **b,** Spermatid nuclei stained with Hoechst (top, left) to target the isolated cell of interest on the EM grid (yellow square) for SEM imaging (bottom, left) prior to FIB milling. Cryo-electron tomography slice (right) with striated microtubules visible within the cells (black arrow), complex membrane organization and stacked mitochondria (black dash contour). Scale bars: 25 µm, 50 µm and 200 nm, respectively. **c,** Cryo-electron tomography slice of an individual striated microtubule corresponding to the microtubule pointed in (b) (top left) and its subtomogram average (bottom left) showing an internal helical organization. Scale bar: 25 nm. Areas boxed as **i** in tomography slice in (b) (middle top) and as **ii** in model in (c) (middle bottom) are shown and oriented to show microtubules surrounded by membranes. Whole model of the tomogram shown in (b). Intricate membranes (orange), mitochondria (blue) and microtubules (green) are shown. Striated microtubule in (c), left, is pointed (white arrow). **d,** Cryo-electron tomography image (sum of 20 slices) of a microtubule aster detached from spermatids after Triton X-100 treatment (left). Scale bar: 200 nm. Dashed line indicates the separation between single-(left) and dual-axis (right) areas of the tomogram. Top black arrow points the minus-end shown and bottom black arrow the microtubule used for subtomogram averaging. Model of the same aster (middle), with microtubules in green. Red dots mark identified microtubule minus ends. Top white arrow points the minus-end shown and bottom white arrow the microtubule used for subtomogram averaging. Example of identified minus-end (top right, top arrow) with a conical shape reminiscent of the γTuRC complex. Scale bar: 50 nm. Cryo-electron tomography slice of an individual striated microtubule observed in detached asters and its subtomogram averaging (bottom right) showing the same internal organization as in intact spermatids. Scale bar: 25 nm.

Using a correlative approach to identify individual spermatids, based on the size of their Hoechst-labeled nuclei (Fig. 3b, top left), we prepared ice lamellae from these cells using cryo-focused ion beam scanning electron microscopy (cryo-FIB SEM) (Fig. 3b, bottom left). Cryo-electron tomograms acquired from these lamellae confirmed the presence of spermatid microtubules (Fig. 3b, right) containing a periodic internal structure (Fig. 3c, top left). Subtomogram averaging of individual microtubules further showed that this internal structure adopts a pseudo-helical organization (Fig. 3c, bottom left). The tomograms also revealed stacked, small, and rather spherical mitochondria (Fig. 3c, blue; dashed-contour in Fig. 3b), as seen in intact cells (Fig. 3a, left panel), and characteristic of elongating spermatids in mammals ^29^. In addition, a complex membrane organization was observed within these cells (Fig. 3c, orange, middle and right panels). Notably, these complex membrane stacks surround individual microtubules (Fig. 3c, middle panels), suggesting a role in cytoplasmic remodeling.

To better characterize these microtubules while preserving their native state as much as possible, spermatid suspensions were treated with 0.1% Triton X-100 immediately prior to cryo-fixation, thereby releasing microtubules from the cells with minimal perturbation. Some released microtubules appeared clustered as asters and were further analyzed using cryo-electron tomography (Fig. 3d, left; Supplementary Fig. 2a). By identifying a subset of microtubule minus ends by their conical shape reminiscent of the γTuRc complex, marked as red dots in the model (Fig. 3d, middle and right, top), we found that microtubules were focused at their plus ends, similar to manchette microtubule plus ends captured at the perinuclear ring in mammalian species, such as rat ^30^. Subtomogram averaging analysis of individual microtubules within these structures (Fig. 3a, right, bottom) confirmed that released microtubules contain the same internal scaffold as in intact cells.

### *Xenopus laevis* spermatid microtubules display extensive structural heterogeneity

Numerous ring-like dense microtubule bundles, reminiscent of early spermatid peripheral networks, could be seen lacking nuclei and plasma membrane (Fig. 4a, left), with a significant proportion of individualized microtubules (Fig. 4a, right). We thus collected cryo-electron microscopy images of released microtubules for high-resolution analysis (Supplementary Fig. 3). Images of microtubules, filtered in Fourier space, displayed distinct fringe patterns that were used to determine protofilament number as previously described ^31^. Within microtubule bundles, we observed a heterogeneous distribution of protofilament number, ranging from 13 to 16 protofilaments (Fig. 4b,c). Surprisingly, we observed a high variability of skew angles for a given protofilament number, including values not previously reported and diverging from the lattice accommodation model, which accurately predicts protofilament skew angles for microtubules assembled *in vitro* ^32^ (Fig. 4d). The most striking diversity occurred in 13 protofilament microtubules, with varied skew angles, indicating a range of lattice conformations (Fig. 4d,e; Supplementary Fig. 4). To estimate the relative abundance of the different microtubule configurations present in spermatid microtubules, we used a reference-based 2D classification approach using microtubule references with varying protofilament numbers and skew angles (Methods; Supplementary Table 1,2; Supplementary Fig. 5,6). This analysis revealed that the most prevalent configuration was made of 13 parallel protofilaments (66%, Fig. 4f), likely adopting a 3-start monomer helical organization such as observed *in vitro* with purified tubulin with a skew angle close to 0°. Yet, 20% of the 13 protofilament microtubules representing 17% of the total had protofilament skew angles, ranging between -0.3° and -4°. The remaining configurations (17%) ranged from 14 to 16 protofilaments. Altogether, this suggests a strong diversity in microtubule configurations present in *Xenopus laevis* spermatid microtubules. Nonetheless, given the limited amount of *in situ* cryo-electron tomography data, we could not determine whether the relative abundance of the different microtubule lattice configurations observed after extraction fully reflects their distribution in intact spermatids. Yet, the internal scaffold produced a layer line at 8 nm^-1^ in each configuration (Supplementary Fig. 4) consistent with the periodicity previously revealed by sub-tomogram averaging (Fig. 3). From this analysis, we chose to focus on the predominant 13 parallel protofilament population (13_3a) for further high-resolution analysis of the proteins present in spermatid microtubules.

**Fig. 4:**
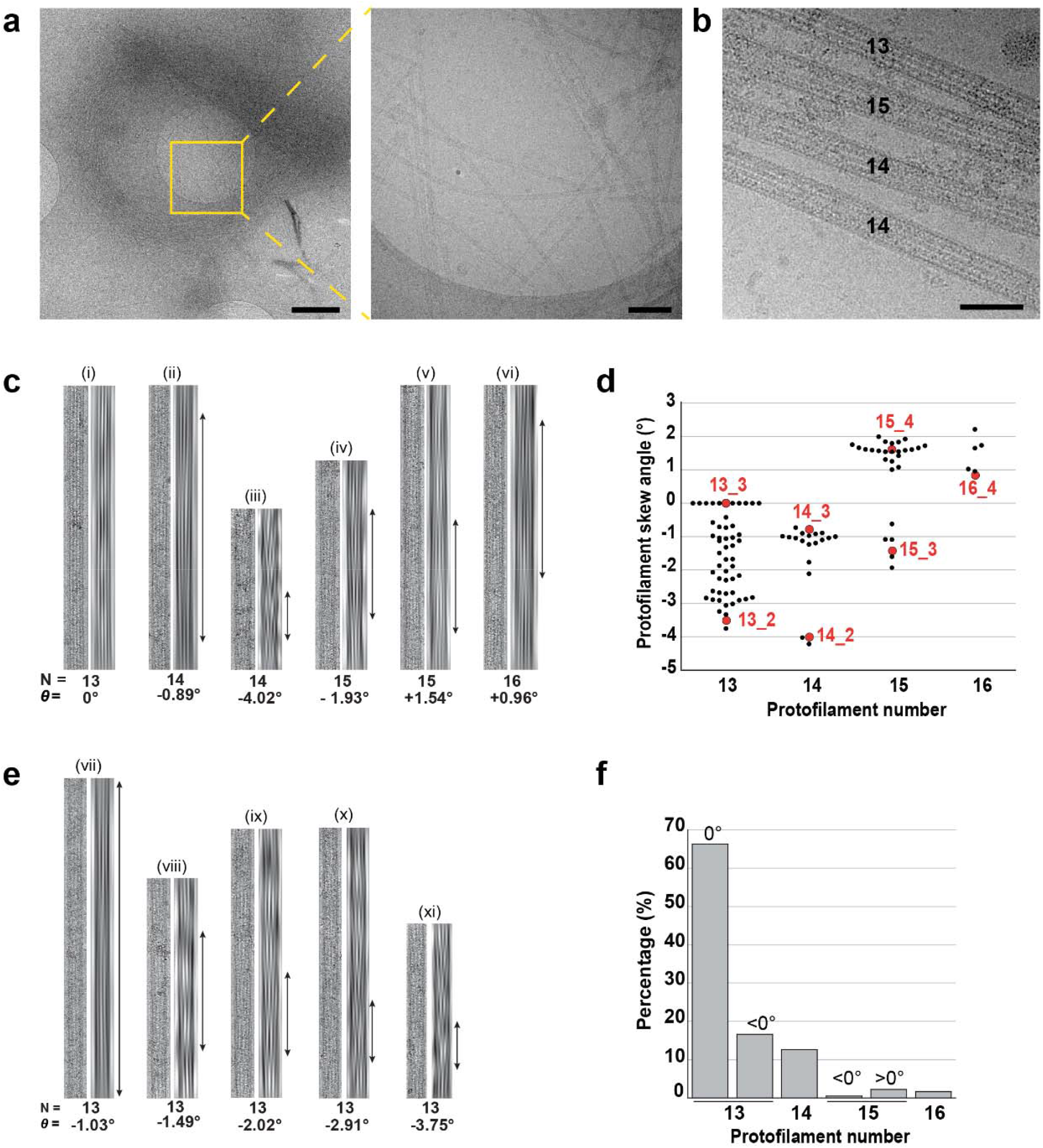
Structural analysis of isolated spermatid microtubules. **a,** Cryo-electron microscopy images of microtubules released from spermatids, circularly organized (left) similarly to peripheral network observed in early spermatids (Fig. 1), with visible individualized microtubules (right). Scale bars: 1 µm and 200 nm, respectively. **b,** High-resolution cryo-electron microscopy images of microtubules with variable protofilament numbers released from spermatids after Triton X-100 treatment. Scale bar: 50 nm. **c,** Analysis of microtubule protofilament number. For each microtubule type, a raw digitally unbent image (left) and the corresponding filtered image with the enhanced fringe pattern (right) are shown. The moiré pattern allows determination of the protofilament number (N) and the absolute protofilament skew angle (*θ*). Arrows indicate distances between the periodicity of the moiré pattern. The sign of *θ* is determined from the relative positions of layer lines at ∼1/4 nm⁻¹ in the corresponding Fourier transforms, as previously described ^79^ (Supplementary Fig. 2). **d,** Distribution of protofilament skew angles measured for different microtubule populations (black circles). Red circles indicate values predicted by the lattice accommodation model for microtubules assembled *in vitro* from purified tubulin with GTP ^32^. **e,** Analysis of 13 protofilament microtubule skew. For each microtubule type, a raw digitally unbent image (left) and the corresponding filtered image with the enhanced fringe pattern (right) are shown. The periodicity of the moiré pattern allows the determination of the skew angle. **f,** Estimation of the percentage of the different microtubule configurations present in spermatid microtubule dataset, based on a reference-based approach (see Methods; Supplementary Fig. 5 for detailed class assignment).

### Spaca9 and Saxo2 are components of the *Xenopus laevis* spermatid microtubule internal scaffold

To characterize the internal proteins forming the microtubule internal scaffold at high resolution, we performed single-particle cryo-EM analysis. Initial helical refinement of the dominant microtubule population, classified as 13_3a, revealed substantial structural heterogeneity at the level of the microtubule lattice, motivating 3D classification to isolate a homogeneous particle subset (Supplementary Fig. 7a,b). This subset was refined to 3.5 Å resolution after CTF refinement. Up to this stage, the imposed helical symmetry and axial sampling resulted in overlapping α-and β-tubulin subunits. To resolve the α/β-tubulin register and examine the luminal proteins in their different lattice environments, we used symmetry expansion followed by focused refinement of either a single short protofilament segment comprising four tubulin subunits or two adjacent short protofilament segments (Supplementary Fig. 7c). The one-protofilament analysis yielded a decorated reconstruction at 3.1 Å resolution over the focused protofilament region (Fig. 5a; Supplementary Fig. 8a,b; Supplementary Table 3). Processing of adjacent protofilament pairs further separated the B-and A-lattice configurations, yielding final decorated maps at 3.2 Å and 3.9 Å resolution, respectively (Fig. 5b, Supplementary Fig. 8a-b; Supplementary Table 3). The decorated one-protofilament and two lattice configuration maps have clearly separated α-and β-tubulin subunits (Supplementary Fig. 8c) and were used for identification and structural analysis of the luminal proteins. The two lattice configurations differed both in protofilament orientation and in the organization of the luminal decoration (Fig. 5c).

**Fig. 5:**
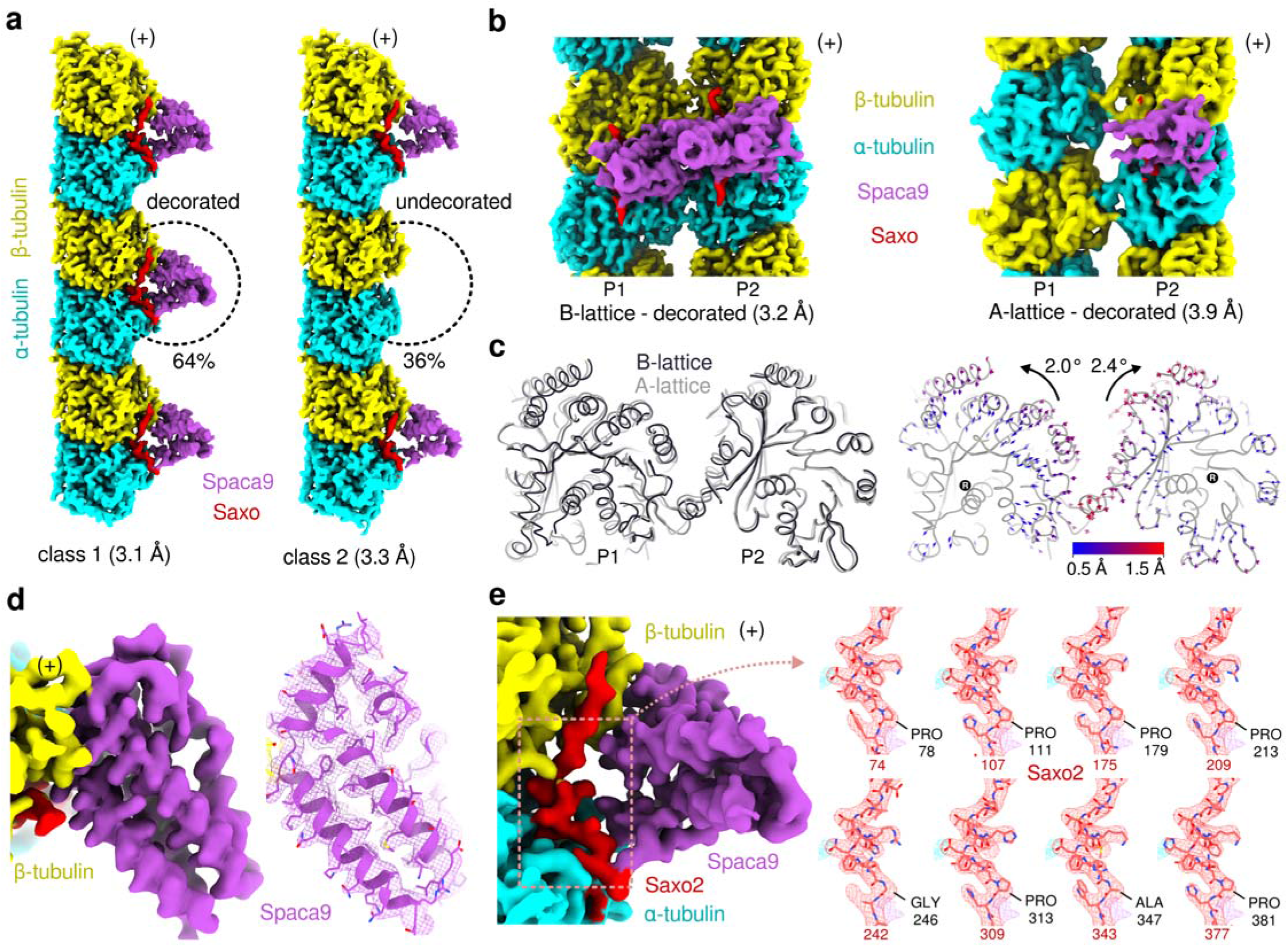
Cryo-EM identification of Spaca9 and Saxo2 as components of the luminal scaffold in *Xenopus laevis* spermatid microtubules. Microtubule polarity is indicated in each panel by a (+) symbol. When the (+) end is oriented toward the top of the figure, the (+) symbol is placed beside the map/model. When the (+) end points toward the viewer, the (+) symbol is placed on the map/model. P1 and P2 denote protofilaments 1 and 2, respectively, in the two-protofilament reconstructions. **a**, High-resolution cryo-EM reconstructions of the one-protofilament classes classified according to the presence or absence of luminal decoration at the central tubulin dimer. **b**, Decorated B-and A-lattice reconstructions obtained by focused refinement of adjacent protofilament pairs, revealing similar internal protein complexes but distinct average patterns of luminal decoration. **c**, Comparison of protofilament orientations in the B-and A-lattice configurations. The B-lattice model was fitted into the A-lattice model, and displacement vectors from the B-to the A-lattice positions were drawn between equivalent Cα atoms separated by at least 0.5 Å, highlighting differences in protofilament rotation. Rotation angles were quantified in ChimeraX around the axis indicated by R. **d**, Cryo-EM map (left) and fitted Spaca9 atomic model (right). A slab view of the model highlights resolved side chains, including hydrophobic residues buried within the protein core, supporting the fit of the distinctive predominantly α-helical fold. **e**, Inset of the decorated one-protofilament cryo-EM map showing the SAXO-like density (left), with the eight candidate Saxo2.L Mn-motif fragments fitted into the density (right). The central residue of each candidate motif is indicated below the corresponding fit. Saxo2 corresponds to UniProt accession Q6DCB9.

The high-resolution maps revealed two distinct components of the luminal density: a folded, predominantly α-helical domain (Fig. 5d) and an approximately 20-residue peptide extending along the tubulin dimers and forming a short helix at the α/β-tubulin intradimer interface (Fig. 5e). This architecture closely resembled the previously described SPACA9–SAXO human complex ^33,34^. ModelAngelo ^35^ readily assigned the larger component to Spaca9, an identification further supported by its distinctive fold (Fig. 5d), but did not identify the smaller peptide. The position of this peptide and the presence of a short helix at the intradimer interface were characteristic of the Mn motifs found in SAXO proteins ^36–38^, suggesting that the peptide originated from an Mn-motif-containing protein. Because the peptide could not be identified directly from the cryo-EM density, we combined sequence constraints derived from the map with mass spectrometry and AlphaFold3 analysis (Methods). This reduced the 1,634 proteins detected by mass spectrometry to a small set of candidates, among which only Saxo2.L showed a predicted interaction on the luminal face of tubulin coincident with the experimental density (Supplementary Fig. 9a,b, 10a,b). Analysis of individual Mn-motif-containing regions further identified eight Saxo2.L candidate fragments that engaged the same tubulin interface (Supplementary Fig. 9c,d, 10c,d) and were collectively compatible with the cryo-EM density (Fig. 5e), consistent with several repeated Saxo2.L Mn motifs being able to occupy the same binding site. In contrast, equivalent AlphaFold3 analysis of Saxo4.L, the only other SAXO protein detected by mass spectrometry, did not identify a compatible tubulin interaction. Saxo2.L was also consistently more abundant than Saxo4.L across both mass-spectrometry replicates (Supplementary Table 4), despite their relatively similar lengths (469 and 384 residues, respectively), with emPAI values of 11.82 versus 3.44 in both replicates and specific spectral counts of 226 versus 7 and 138 versus 12, respectively (Supplementary Table 5). Together with the structural analyses, these results support Saxo2.L as the major source of the peptide density and identify the luminal complex as comprising Spaca9.L and Saxo2.L. Accordingly, the peptide density was modeled using one of the Saxo2.L candidate Mn motifs (Saxo2.L_209, residues 199–218; Fig. 5e). Saxo2.L and Spaca9.L are hereafter referred to as Saxo2 and Spaca9, respectively.

### A discontinuous Spaca9–Saxo2 luminal scaffold connects adjacent protofilaments in *Xenopus laevis* spermatid microtubules

A distinctive feature of *Xenopus laevis* spermatid microtubules is a direct interaction between Spaca9 molecules bound to neighboring protofilaments. Like human SPACA9, *Xenopus laevis* Spaca9 contains a long C-terminal extension beyond the conserved folded domain, although this region is poorly conserved in sequence. When an adjacent Spaca9 molecule is present, the proximal segment of this extension becomes ordered and docks along the lumen-facing tip of the neighboring Spaca9 (Fig. 6a). The PGPQ segment (residues 162– 165), resolved at the backbone level, lies in proximity to Thr85, Val105 and Pro110 of the adjacent molecule, thereby directly connecting Spaca9 molecules across neighboring protofilaments (Fig. 6b). In classes lacking the neighboring Spaca9, this tail segment is not resolved (Fig. 6c), indicating that its ordering depends on the intermolecular contact. This specific interaction is not apparent in the previously reported high-resolution human and bovine sperm SPACA9 structures ^33,34^. These Spaca9–Spaca9 interactions may contribute to stabilizing the lumen-facing tip of Spaca9, which extends toward smaller microtubule radii and appears more positionally or conformationally heterogeneous based on its lower local resolution (Supplementary Fig. 8b).

**Fig. 6:**
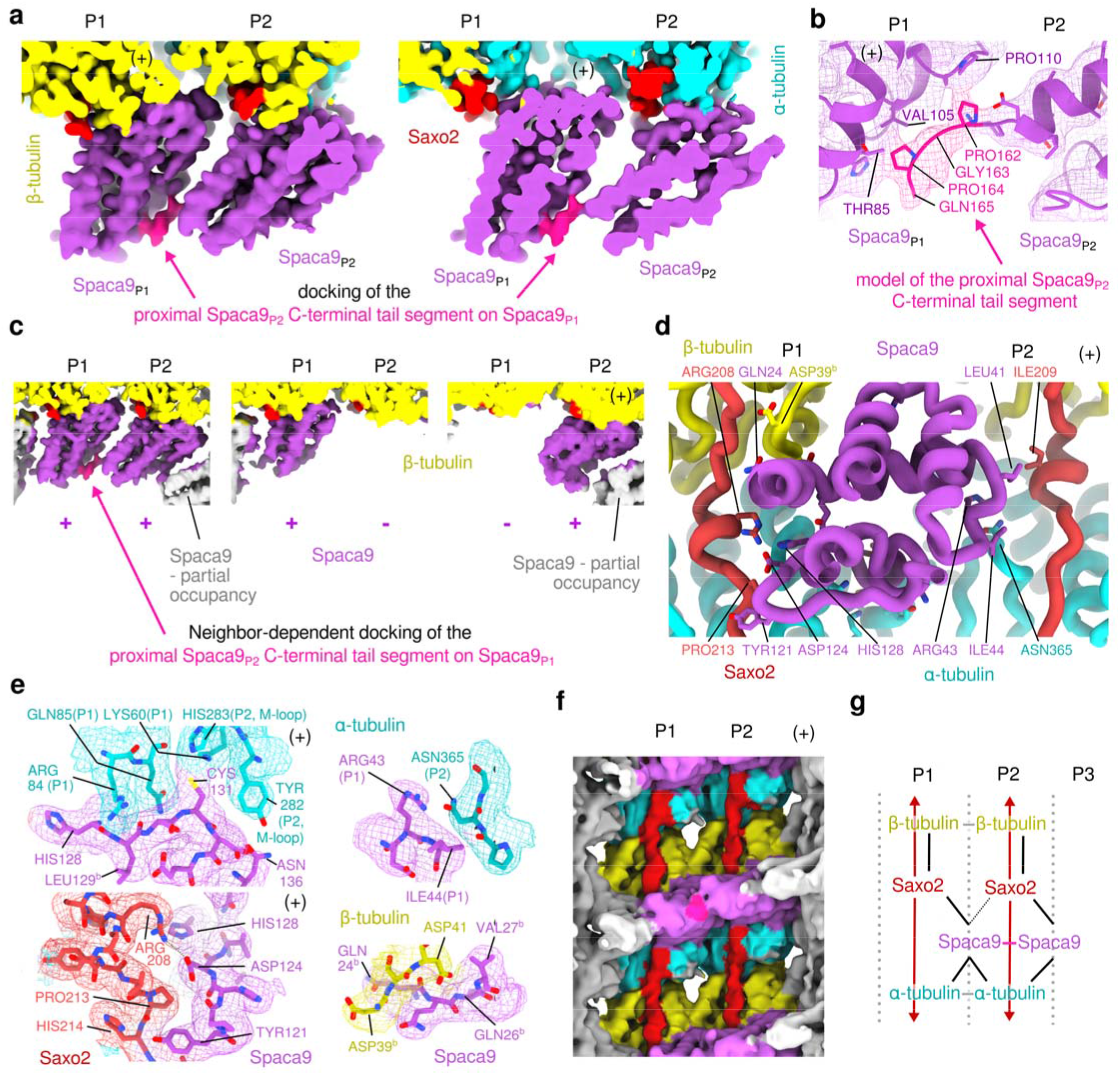
Molecular organization of the Spaca9-Saxo2 luminal scaffold on *Xenopus laevis* **spermatid microtubules.** Microtubule polarity and protofilaments 1 and 2 are indicated as in Fig. 5. **a**, Slab views of the Spaca9 map in the decorated B-lattice reconstruction at two positions along the protofilament pair. An additional density connects neighboring Spaca9 molecules and extends from the C-terminal region of the Spaca9 molecule on P2. **b**, Atomic modelling showing that the connecting density in panel f is consistent with the proximal region of the Spaca9 C-terminal tail, defined here as the segment following the last residue of the C-terminal helix. The exact side-chain positions remain uncertain owing to the lower local resolution in this region (Supplementary Fig. 8b). Spaca9 tail residues and candidate contact residues from the neighboring Spaca9 are shown as sticks. Spaca9 corresponds to UniProt accession A0A8J0PW71. **c**, Class averages from 3D classification of the B-lattice internal protein densities, showing the three observed decoration patterns. The density connecting neighboring Spaca9 molecules shown in (a) and (b) is resolved only when both the P1 and P2 positions are decorated, indicating that the proximal C-terminal tail of Spaca9 at P2 becomes ordered upon interaction with the neighboring Spaca9 at P1. **d**, Overview of the main contacts between Spaca9 and the five neighboring proteins in the decorated B-lattice complex model (on P1: α-tubulin, β-tubulin and Saxo2; on P2: α-tubulin and Saxo2). The model is viewed from the lumen, with the portion of Spaca9 extending toward the lumen cropped to better visualize the cross-protofilament contacts near the tubulin interface. Spaca9 residues and candidate interacting residues from neighboring proteins are shown as sticks. Residues interacting only through their backbone are indicated by b. The α-and β-tubulin isotypes used in the models correspond to Tuba5.S (UniProt accession code: Q7ZTP0) and Tubb4b.L (UniProt accession code: P30883), respectively. **e**, Insets of the decorated B-lattice map showing different types of Spaca9 interactions with surrounding proteins. Candidate interacting residues from neighboring proteins are shown as sticks. Residues interacting only through their backbone are indicated by b. **f**, B-lattice map low-pass filtered to 7 Å and displayed at a low threshold, showing apparently continuous Saxo-associated densities along the protofilaments. **g**, Schematic summarizing the protein-interaction network derived from the decorated B-lattice reconstructions presented in this figure. Solid lines indicate resolved interactions and the thin dashed line indicates a putative interaction.

The overall discontinuous organization of the Spaca9 scaffold is reminiscent of the interrupted luminal helix originally observed in human sperm singlet microtubules ^39^, which was subsequently shown to comprise noncontinuous SPACA9 spirals interrupted at the seam ^33^. In the *Xenopus laevis* reconstructions, Spaca9 density was likewise absent from the same protofilament bordering the seam (the left-hand protofilament when viewed toward the plus end, Fig. 5b), but occupancy was additionally partial: 64% of tubulin-dimer particles were classified as decorated and 36% as undecorated at the central dimer. In the latter class, the neighboring dimers toward both the plus and minus ends remained partially decorated (Fig. 5a), indicating that the Spaca9 array can also be locally interrupted along individual protofilaments. Whether this partial occupancy is specific to *Xenopus laevis* spermatid microtubules or represents a more general feature of SPACA9–SAXO arrays revealed by the focused classification used here remains to be determined.

Beyond this intermolecular interaction, Spaca9 engages the microtubule lattice through an extensive cross-protofilament interface similar to that described for human ciliary SPACA9 ^33^. Spaca9 simultaneously contacts α-and β-tubulin within one protofilament and α-tubulin from the adjacent protofilament, with several interactions involving the α-tubulin M-loop (Fig. 6d,e). These contacts provide a direct structural basis for reinforcement of the lateral protofilament interface, consistent with the stabilizing role previously proposed for the SPACA9–SAXO system ^33^. As in the previously described human SPACA9–SAXO complex ^33^, Spaca9 may also contact a second SAXO segment, here identified as Saxo2 and associated with the neighboring protofilament, although this putative interaction is limited to a single-residue contact (Fig. 6d). In the decorated B-lattice map low-pass filtered to 7 Å, Saxo-associated densities appear connected along the same protofilaments (Fig. 6f). The connected density suggests that at least a fraction of Saxo2 extends longitudinally along individual protofilaments. Such an arrangement has structural precedents among Mn-motif proteins. In sperm microtubules, SAXO proteins containing repeated Mn motifs can extend longitudinally along individual protofilaments, as shown for SAXO5 ^34^.

Together, these structures suggest that the lumen of spermatid microtubules contains a mechanically connected yet geometrically adaptable scaffold (Fig. 6g). Spaca9 provides lateral reinforcement by spanning neighboring protofilaments and engaging the lateral interface, including the α-tubulin M-loop, while neighboring Spaca9 molecules can be further coupled through a conditional C-terminal tail interaction. In the orthogonal direction, repeated Saxo2 interactions may provide longitudinal connectivity along individual protofilaments, with Spaca9–Saxo2 contacts linking these two components of the network.

### Spaca9 and Saxo2 are specifically expressed in spermatids and part of their microtubule network

Earlier studies of *Xenopus* spermatogenesis reported the absence of a manchette structure and suggested that microtubules from Sertoli cells were involved in shaping the spermatid nucleus. To further validate whether the microtubules that we analyzed derived from spermatids and not from Sertoli cells, we examined the expression of *spaca9* and *saxo2 in situ* (Fig. 7a). Importantly, previous work showed that both genes are expressed from the L, but not the S, subgenome of the *Xenopus laevis* allotetraploid genome ^40^, as also confirmed by our proteomic data (Supplementary Table 4). We therefore focused our analysis on *spaca9.L* and *saxo2.L*. Testis lobules were dissected and processed by RNA-FISH. Both mRNAs were prominently detected in cysts of putative spermatids, characterized by low expression of the spermatocyte marker *mybl1.L*, adjacent to spermatocyte cysts showing high *mybl1.L* expression. Importantly, *spaca9.L* and *saxo2.L* were not detected around the cysts, supporting that Sertoli cells do not express either gene. Similarly, no *spaca9.L* or *saxo2.L* signal was detected in mature sperm cells. Furthermore, analysis of previously published single-cell RNA-seq datasets ^41^ consistently showed that *spaca9.L* and *saxo2.L* are more strongly expressed in spermatids, whereas *mybl1.L* is, as expected, mostly expressed in spermatocytes (Fig. 7b).

**Fig. 7:**
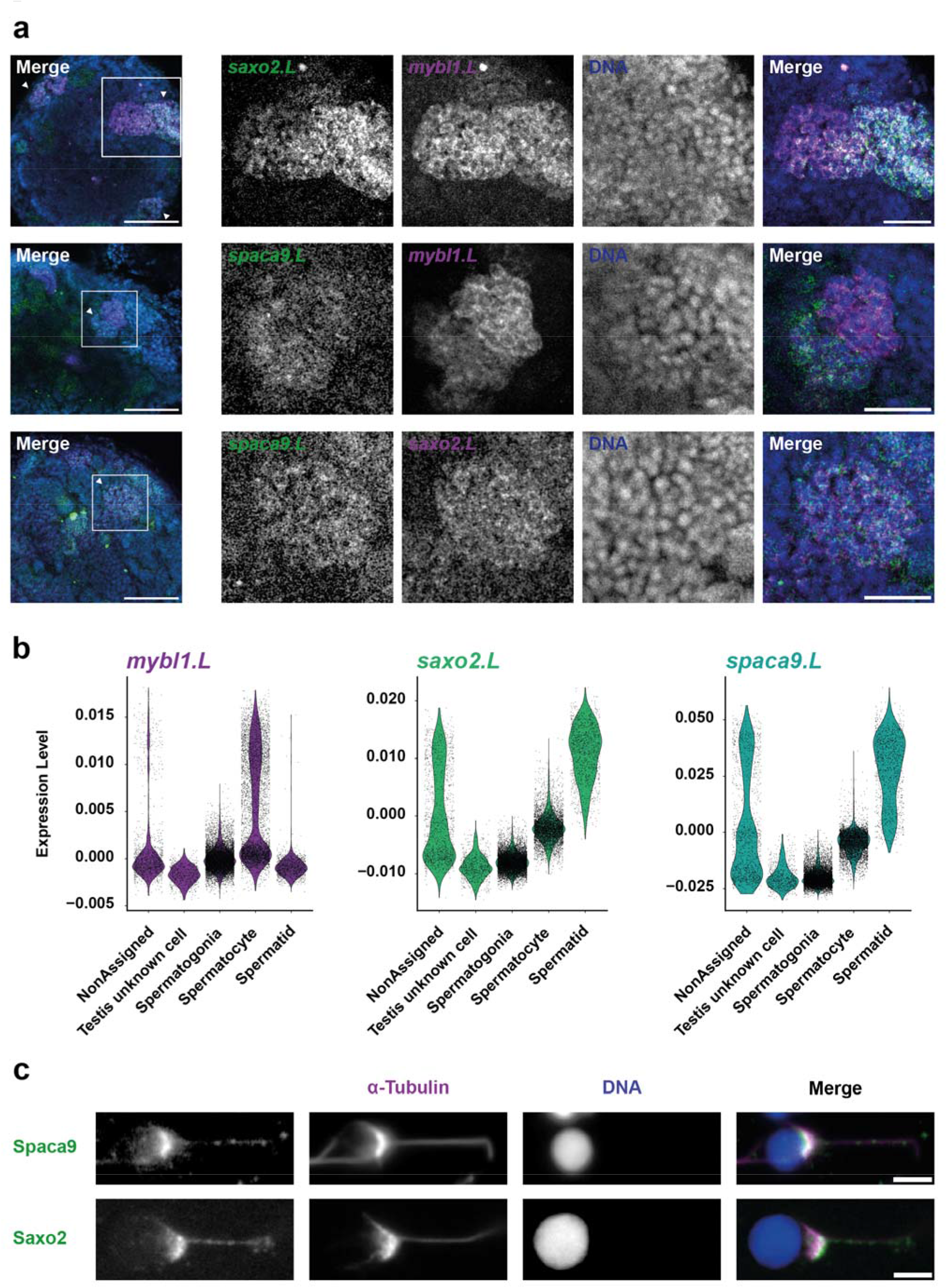
Analysis of *spaca9.L* and *saxo2.L* expression in *Xenopus laevis* spermatids. **a,** RNA-FISH analysis of testis lobules using probes for *mybl1.L*, *saxo2.L,* and *spaca9.L*. *saxo2.L* (green) and *mybl1.L* (magenta) (upper panel; 3 lobules analyzed), *spaca9.L* (green) and *mybl1.L* (magenta) (middle panel; 2 lobules analyzed), *spaca9.L* (green) and *saxo2.L* (magenta) (lower panel; 2 lobules analyzed). Cysts of interest are shown. Scale bars: 100 µm, in low magnification merge views, and 40 µm in magnified cysts. **b,** Expression levels of *mybl1.L*, *saxo2.L,* and *spaca9.L* in germ cells. Single-cell RNA-seq dataset from Liao *et al.*, 2022 ^41^ was used. **c,** Immunofluorescence images of isolated spermatids. Microtubules (magenta), DNA (blue), and Spaca9 (top) or Saxo2 (bottom) (green) are shown. Scale bars: 5 µm.

Finally, immunofluorescence against Spaca9 and Saxo2 confirmed the labeling of spermatid microtubules, particularly the polarized network (Fig. 7c, Supplementary Fig. 1e,f). Together, these results demonstrate that *spaca9.L* and *saxo2.L* are specifically expressed in spermatids and confirm that their protein products are part of the spermatid microtubule network.

### The spermatid microtubule internal scaffold is conserved between *Xenopus laevis* and ***Xenopus tropicalis***

To assess whether the microtubule internal structure uncovered in *Xenopus laevis* is conserved across *Xenopus* species, we examined the distant species *Xenopus tropicalis*, who diverged from a common ancestor ∼48 million years ago ^40^. We first analyzed *Xenopus tropicalis* germ cells *in situ*. Using fluorescent histochemistry (Fig. 8a, left), we identified spermatids as cells with nuclei measuring 4.4 ± 0.8 µm in diameter (Fig. 8a, right). As expected, this size is smaller than in *Xenopus laevis*, consistent with the biological scaling at play in *Xenopus* species of different sizes ^19^.

**Fig. 8:**
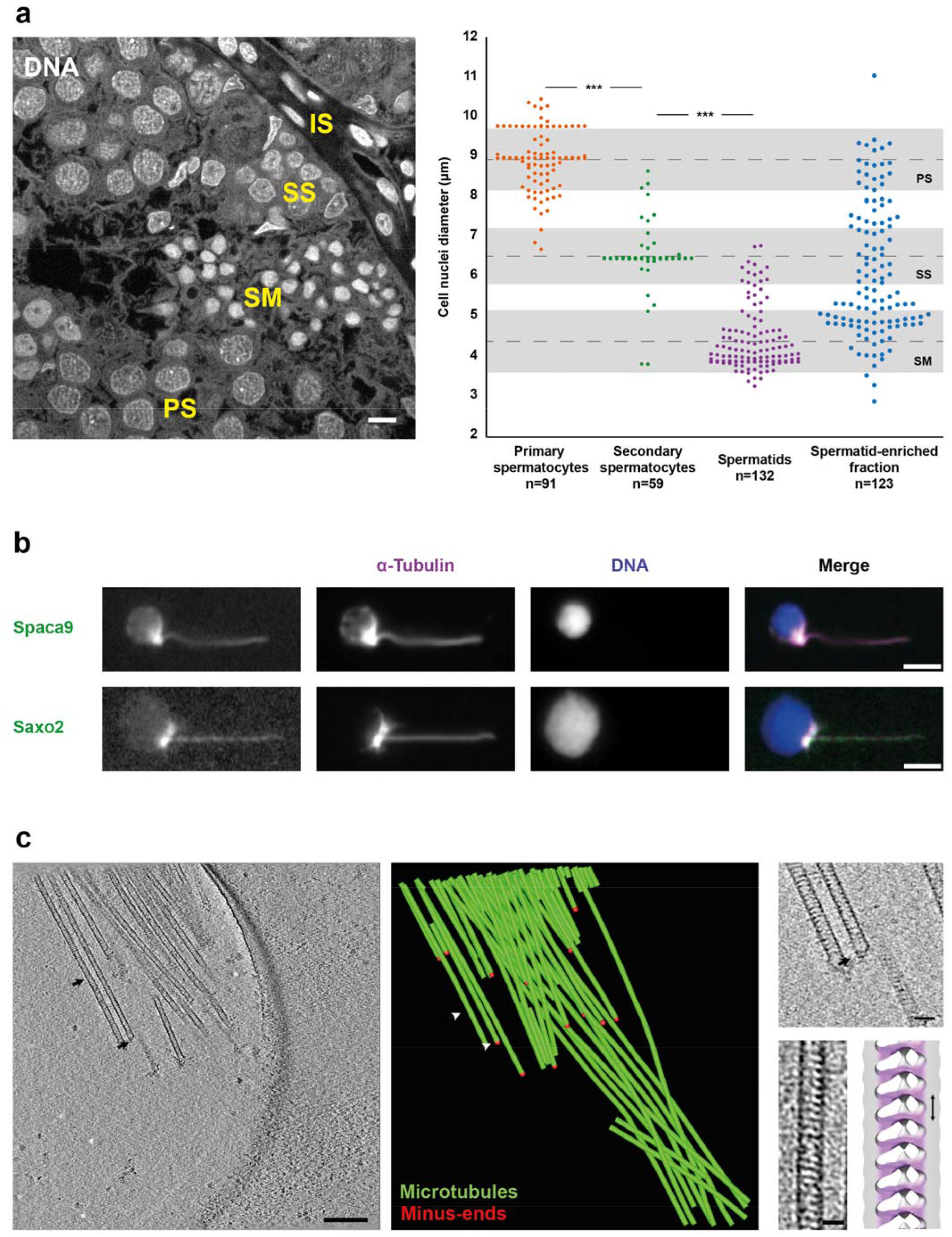
*Xenopus tropicalis* spermatid analysis. **a,** Image of cysts of germ cells visualized *in situ* by nuclear staining using Hoechst (left panel). Intertubular space (IS), primary spermatocytes (PS), secondary spermatocytes (SS), spermatids (SM), and sperm cells (SR) are labeled. Scale bar: 10 µm. Diameters of nuclei in primary spermatocytes, secondary spermatocytes, and spermatids, with a mean of 8.9 ± 0.8 µm, 6.5 ± 0.7 µm and 4.4 ± 0.8 µm, respectively, and of nuclei in the spermatid-enriched fraction (right panel). Dashed lines indicate the mean and the gray boxes the standard deviation, plotted for primary spermatocytes, secondary spermatocytes, and spermatids and labeled as PS, SS, and SM, respectively. **b,** Immunofluorescence images of isolated spermatids. Microtubules (magenta), DNA (blue), and Spaca9 (top) or Saxo2 (bottom) (green) are shown. Scale bars: 5 µm. **c,** Cryo-electron tomography image (sum of 20 slices) of an aster of microtubules detached from spermatids after Triton X-100 treatment (left). Scale bar: 200 nm. Model of the same aster (middle), with microtubules in green. Bottom white arrow points the minus-end shown and top white arrow the microtubule used for subtomogram averaging. Image of γ-TuRC of pointed microtubule minus ends (right, top). Scale bar: 50 nm. Cryo-electron tomography slice of an individual striated microtubule observed in detached asters and its subtomogram averaging (right, bottom) showing similar internal densities as in *Xenopus laevis* spermatids. Scale bar: 25 nm.

We next prepared a spermatid-enriched cell suspension and analyzed the cells using immunofluorescence. As observed in *Xenopus laevis*, spermatids showed a polarized microtubule network at the caudal side of the nucleus, with both Spaca9 and Saxo2 colocalizing with this structure (Fig. 8b).

Finally, spermatid-enriched cell suspensions were treated with 0.1% Triton X-100 prior to cryo-fixation to release their microtubules. Cryo-electron tomography revealed detached microtubules containing an internal repeat, and subtomogram averaging confirmed the pseudo-helical nature of the internal structure (Fig. 8c; Supplementary Fig. 2b).

Together, these results provide compelling evidence that the presence of a spermatid microtubule internal scaffold is conserved across divergent *Xenopus* species.

## Discussion

In this study, we reveal the existence of a complex spermatid microtubule network in *Xenopus*. Moreover, these microtubules adopt unexpectedly diverse lattices and contain an internal scaffold containing Spaca9 and Saxo2 in *Xenopus laevis*, similar to the SPACA9 and SAXO interrupted spiral previously characterized in human sperm singlet microtubules ^33^.

Previous studies reported the absence of a manchette in *Xenopus* ^20,21^. Based on conventional electron microscopy of chemically fixed, dehydrated and resin-embedded testis tissue, the authors concluded that microtubules were nearly absent within *Xenopus* spermatids, instead proposing that microtubules participating in shaping the nucleus originated from Sertoli cells ^20,21^. Conversely, in our work, (i) immunofluorescence analysis of isolated spermatids, where Sertoli cells are eliminated during isolation, revealed a polarized skirt-like microtubule network at the caudal side of the nucleus that bears molecular markers similar to those of mammalian manchettes; (ii) proteomics analysis of purified spermatid microtubules revealed an enrichment in cytoskeleton components reminiscent of the mammalian manchette; (iii) cryo-electron microscopy of intact isolated spermatids confirmed a dense microtubule network in these cells; and (iv) microtubules released from spermatids were occasionally observed to be focused at their plus ends, as described for manchette microtubules. The identification of Spaca9 and Saxo2 as specific components of spermatid microtubules, together with RNA-FISH and single-cell RNA-seq showing that their expression is specific to spermatids, provided decisive evidence for the presence of a spermatid-specific microtubule network with manchette-like characteristics.

From an organizational perspective, *Xenopus* spermatid microtubules display notable similarities to the cortical microtubules of *Toxoplasma gondii* ^38^. This parasite possesses 22 cortical microtubules that originate from an apical polar ring and extend to cover two-thirds of the cell body, forming a basket-like structure, reminiscent of the polarized manchette organization, and conferring the rigidity required to maintain cell shape and resist external forces ^42^. Strikingly, both systems rely on an internal helical structure, TrxL1/2-SPM1 in *Toxoplasma gondii* and Spaca9–Saxo2 in *Xenopus* spermatids, or similarly SPACA9–SAXO in human sperm singlet microtubules ^33^, despite lacking sequence and structural homology. As highlighted by Gui and colleagues ^33^, in both helical structures, one protein binds longitudinally along protofilaments (SPM1 or SAXO), while the partner protein binds across protofilaments (TrxL1/2 or SPACA9), suggesting a striking convergence toward an architectural solution for reinforcing microtubules against mechanical stress. Gui and colleagues therefore proposed that such an internal scaffold, which stabilizes microtubules to counter external stress in *Toxoplasma gondii*, would have a similar role in sperm. Our data support such a model by revealing a similar luminal scaffold within a system subjected to extreme mechanical constraints necessary for spermatid differentiation.

The role of SAXO proteins (SAXO, short for “stabilizer of AXOnemal microtubules”) in microtubule stabilization has been previously documented. SAXOs constitute a MAP6-related protein family and share repeated Mn-like microtubule-binding motifs with MAP6 (STOP, short for “Stable Tubule Only Polypeptide”) proteins. In MAP6 proteins, Mn modules contribute to microtubule stabilization and resistance to cold-and nocodazole-induced depolymerization ^36,43^. SAXO1 was similarly shown to rely on its Mn motifs for microtubule binding and stabilization upon cold treatment, while specifically localizing to ciliary and flagellar microtubules ^44^. Comparative analyses of microtubule inner proteins subsequently identified Mn repeats as a conserved hallmark of several luminal microtubule-associated proteins, including MAP6, SAXO1 and SAXO2 ^45^. High-resolution structures of sperm axonemal doublet microtubules further revealed that SAXO proteins interact with the luminal surface through multiple Mn motifs, binding to both individual and neighboring protofilaments, consistent with a role in stabilizing the microtubule lattice ^34^.

Meanwhile, the SPACA9 protein recently emerged as a structurally versatile microtubule inner protein that differentially associates with several distinct microtubule architectures in motile cilia, sperm flagella and developing spermatids ^33,34,46,47^. In human respiratory cilia, SPACA9 forms three-copy intraluminal striations that repeat every 8 nm along protofilaments B02–B05 of the B-tubule of the axoneme. Each molecule binds at the luminal α/β-tubulin intradimer interface, contacting adjacent α-tubulin molecules, including their M-loops, while simultaneously interacting with SAXO-family proteins ^33^. This binding mode appears to be conserved in the sperm axoneme ^34^. In the distal singlet microtubules of human sperm, SPACA9 is distributed much more extensively, with 12 copies per 8-nm repeat forming discontinuous spirals interrupted at the microtubule seam ^33^. SPACA9 is also present within the B-tubules of mammalian sperm outer doublets, where SPACA9 molecules are arranged in 3 to 5 adjacent ones that are repeated longitudinally and contribute, together with a putative SAXO1 filament, to a luminal protein network ^34,48^. In mouse sperm, related but distinct SPACA9 distributions have been observed in both axoneme central-pair microtubules, C1 and C2, with 3 to 7 adjacent SPACA9 repeats and one protofilament decorated with SPACA9 without adjacent neighbors, further illustrating the variability of binding geometries to different microtubule environments ^46^. Despite these variable stoichiometries and distributions, SPACA9 preserves a similar structure on the luminal side of protofilament pairs, where it bridges longitudinal and lateral tubulin contacts and frequently associates with additional luminal proteins, including SAXO proteins ^33,34^. Our data revealed an extensive Spaca9–Saxo2 decoration with additional connections between neighboring Spaca9 molecules, which together strongly support a role in microtubule stabilization. Consistent with a stabilization role of Spaca9, recent *in vitro* reconstitutions showed that it inhibits dynamic instability and stabilizes protofilaments at growing microtubule ends ^49^. A markedly different organization of SPACA9 was recently identified within spermatid manchette microtubules in rat, which undergoes a substantially different degree and geometry of nuclear reshaping compared with *Xenopus.* There, only a single SPACA9 molecule repeats every 8 nm specifically along protofilament 1 at the seam and associates with the SH3-domain protein MNMIP1 (Sh3d21 in *Xenopus*) ^47^. In this complex, SPACA9 contacts both α-and β-tubulin of protofilament 1, whereas MNMIP1 interacts with SPACA9 and extends across the seam to bind β-tubulin in protofilament 13, together bridging the seam ^47^. These differences from the microtubules analyzed in this work suggest that distinct evolutionary solutions have emerged to stabilize spermatid microtubules during nuclear remodeling.

In addition, we found that spermatid microtubules in *Xenopus* vary in protofilament number, with extended variation of skew angles and configurations not reported *in vivo* or *in vitro*. Despite reports documenting some lattice variability *in vivo* ^50–53^, previous work in *Xenopus* showed that microtubules assembled in egg extracts predominantly adopt a 13-protofilament configuration with no skew, with only a very low degree of variability, comprising about 1% of each 12-and 14-protofilament microtubules ^54^. By comparison, *in vitro* GTP-assembled microtubules revealed that ∼33% and ∼65% of the total microtubule length correspond to 13-and 14-protofilament lattices, respectively, with only ∼1% each of 12-and 15-protofilament lattices ^54^. Here, we show that microtubules in *Xenopus* spermatids populate a very broad set of lattice configurations, with substantial proportions (17%) of 14-, 15-, and 16-protofilament configurations, all with varying skew angles, that deviate from the canonical architecture, particularly in 13-protofilament microtubules with skew angles ranging between 0° and -4°. The persistence of highly skewed lattices and otherwise unfavorable microtubule geometries supports a stabilization role for the Spaca9–Saxo2 scaffold. *In vitro*, measured protofilament skew angles indeed rarely exceed ±2° ^32^. Therefore, the scaffold may allow both otherwise unstable skewed configurations and mechanically induced lattice distortions to persist.

Beyond stabilizing microtubules ^33,34,49^, a Spaca9–Saxo2 scaffold could impose geometrical constraints on the microtubule lattice that influence the organization of tubulin heterodimers and protofilament number. In microtubules with fewer protofilaments, steric clashes between adjacent Spaca9 molecules may limit their assembly or require alternative protofilament skews to accommodate the luminal decoration, whereas increasing the protofilament number would reduce steric hindrance and permit lattice geometries closer to those predicted by the lattice accommodation model ^32^. Such a mechanism could explain the reduced skew-angle diversity observed in wider microtubules compared with that in 13-protofilament microtubules. In addition, because increasing microtubule diameter can theoretically increase their stiffness, favoring microtubules with a higher number of protofilaments could provide an additional means of modulating the mechanical properties of spermatid microtubules. Although this hypothesis remains speculative and will require deeper structural analysis strategies, it raises the possibility that luminal proteins not only reinforce unfavorable microtubule architectures, but actively shape the geometry of the microtubule lattice during assembly. Another appealing, non-exclusive hypothesis is that spermatid microtubule lattices remain mechanically responsive through their internal scaffold, exhibiting variable skew as a consequence of the forces they experience, with an internal scaffold limiting their depolymerization. Cryo-fixation may therefore capture transient intermediates of these dynamic lattice rearrangements, explaining the observed wide range of skew angles. By permitting a broader range of lattice architectures and distortions than previously reported, together with increased stiffness of microtubules with a higher number of protofilaments, the internal scaffold may enable these microtubules to explore and stabilize various lattice conformations. This could confer enhanced mechanical resilience and adaptability required to withstand the substantial forces associated with nuclear reshaping and cytoplasm remodeling during spermiogenesis.

Another striking observation is that isolated *Xenopus* spermatids contain individual microtubules surrounded by membranes, suggesting a possible role for the spermatid microtubule cytoskeleton in cytoplasm remodeling. Together with nuclear reshaping, cytoplasm remodeling is a key process for successful spermiogenesis. In mammals, this process relies on tubulobulbar complexes, spermatid plasma membrane projections extending into Sertoli cells and surrounded by Sertoli-cell actin machinery ^55,56^. These complexes contribute to the removal of excess cytoplasm that forms residual bodies subsequently phagocytosed by Sertoli cells prior to sperm release ^57^. While *Xenopus* spermiogenesis has been described morphologically and ultrastructurally ^20,58,59^, no studies have addressed the mechanisms underlying cytoplasm elimination. These observations open the possibility that, in *Xenopus*, cytoplasm removal may involve spermatid-intrinsic mechanisms, rather than relying exclusively on Sertoli-cell–driven processes.

Altogether, our findings offer new perspectives on *Xenopus* spermiogenesis. They highlight a dedicated regulation of microtubules that likely contributes to extreme morphogenetic processes characteristic of *Xenopus* species and may be essential for sperm integrity and fertility. The precise contribution of the discovered spermatid microtubule network to nuclear reshaping remains an open question, and additional mechanisms previously proposed to influence nuclear morphology, including those involving nurturing Sertoli cells, may also be at play ^1–5^. Moreover, whether a similar extensive internal scaffold is present in the manchette microtubules of other species and adapted through evolution remains unknown and represents an exciting direction for future research into conserved and divergent strategies of spermiogenesis.

## Methods

### Animals

Mature *Xenopus* males were used for animal experimentation according to our animal use protocol APAFiS #45521-2023101308502762 approved by the Animal Use Ethics Committee (#7, Rennes, France) and the French Ministry of Higher Education, Research and Innovation.

### Chemicals

Unless otherwise stated, all chemicals were purchased from Sigma-Aldrich, Merck.

### Tissue collection and histological analysis

Samples were fixed in 4% formalin (pH 7) 24 h before paraffin inclusion.

For Hematoxylin-Eosin staining, 4 µm tissue sections were deparaffinized with a mix of xylene/alcohol, then washed and stained with Gill II hematoxylin (Thermo Fisher Scientific). After washes, sections were incubated with bluing reagent (Thermo Fisher Scientific) and Eosin Y (Thermo Fisher Scientific). After washes, tissues were incubated with Safran (Microm Microtech France). Finally, tissue sections were dehydrated with a mix of toluene/alcohol before mounting. Mounted sections were finally imaged using a NanoZoomer S60 scanner (Hamamatsu).

For fluorescent staining, following deparaffination with Discovery wash solution (Ventana) at 75 °C for 8 min, antigen retrieval was performed using Ventana proprietary, Tris-based buffer solution pH 8, at 95 °C for 40 min. Endogenous peroxidase was blocked with 3% H_2_O_2_ for 12 min. After rinsing, slides were incubated at 37 °C for 60 min with primary antibody mouse anti-tubulin (dilution at 1/100). Signal enhancement was performed using a goat anti-mouse HRP at 37 °C for 16 min and DISCOVERY FAM Kit (490-520 nm) (Roche Diagnostic) for 8 min. To visualize the nucleus, DAPI staining was added and coverslipped. Single plan images were acquired on a Zeiss LSM 900 Axio Observer 7 confocal microscope using the Zen software and a Plan-Apochromat 63x/1.40 Oil DIC objective. Images are mean averages of two unidirectional scans with a depth of 16 bits. Pinhole size was chosen to correspond to 1 Airy unit. Images were processed using the Zeiss LSM plus module and then analyzed using Fiji ^60^.

### *Xenopus* spermatids enrichment

This isolation protocol was adapted from Teperek et al. ^23^. One testis from adult *Xenopus laevis* was isolated and manually cleaned from blood vessels and fat bodies in MMR (Marc’s Modified Ringer’s: 5 mM HEPES; 0.1 mM EDTA; 100 mM NaCl; 2 mM KCl; 1 mM MgCl_2_; 2 mM CaCl_2_; pH 7.8). Testis was cut in pieces with scissors, briefly spun, torn again, and homogenized manually with a pestle in 1 mL MMR. After 50-µm filtration (CellTrics cat. 04-0042-2317), the suspension was spun down at 800 g in AM2.18 rotor (Jouan MR23i centrifuge), at 16 °C, for 20 min, and the cell pellet was resuspended in 2 mL of MMR. Step gradients of iodixanol (Optiprep; Sigma, D1556; 60% iodixanol in water) in MMR were manually prepared in 14 mL glass tube (Kimble® HS No. 45500-15) in the following order from the bottom to the top of the tube: 4 mL of 30% iodixanol, 1 mL of 20% iodixanol, 5 mL of 12% iodixanol (all diluted in MMR), and 2 mL of cell suspension in MMR on top. Gradients were spun down at 10,000 g using JS13.1 rotor (Beckman Coulter; Avanti JXN-26 centrifuge), at 16 °C, for 15 min, deceleration without brake. The top interface fraction (between MMR and 12% iodixanol cushion), enriched in spermatids, was collected and diluted six times with MMR and then pelleted by spinning at 3,220 g in AM50C.13 rotor (Jouan MR23i centrifuge), at 16 °C, for 20 min. Pelleted cells were resuspended in about 500 µL MMR. The proportions of cell types within the isolated fraction were estimated as follows. The size distributions of the three reference cell types (PS, SS and SM) were estimated by kernel density estimation using a common bandwidth. The mixed-population distribution in the isolated fraction was modeled as a weighted mixture of these three reference distributions, with non-negative mixture proportions constrained to sum to one and estimated by maximum likelihood. Uncertainty in the estimated proportions was assessed by bootstrap resampling of the mixed population with replacement (1,000 iterations), while keeping the reference distributions fixed, and is reported as ± standard deviation.

### Immunofluorescence of spermatid-enriched cell suspension

Immunofluorescence was performed as previously described ^61^. A volume of 10 to 50 µL of spermatids was gently mixed in 1 mL of 30% v/v glycerol in BRB80 supplemented with 0.5% Triton X-100 and 2.5% formaldehyde, layered on a 5 mL 40% v/v glycerol in BRB80 cushion, and centrifuged for 16 min at 17,000 g, using JS13.1 rotor (Beckman Coulter; Avanti JXN-26 centrifuge), at 16 °C. Cells spun on a coverslip were fixed in cold methanol for 5 min, rehydrated 3 times with PBS-NP40 0.1% and blocked in PBS-BSA (w/v) 3% for 1 h at room temperature. Slides were incubated with primary antibody diluted in PBS-BSA 3% for 1 h at room temperature. After 3 washes with PBS-NP40, slides were incubated with secondary antibody diluted 1:1,000 in PBS-NP40 for 30 min at room temperature and finally washed 3 times with PBS-NP40 before mounting on a slide with antifade mounting medium containing DAPI (Vectashield, H-1200). Images were acquired on an Olympus BX51 microscope, equipped with ORCA-ER (Hamamatsu) or Prime BSI (Photometrics) cameras, using the micromanager software v1.4 ^62^ and analyzed using Fiji. Importantly, finding spermatids at all stages for imaging was limited by their uneven representation within the enriched cell suspension, in which early, mid, and late spermatids accounted for 37.8%, 3.2%, and 0.5% of all cells, respectively. The following primary antibodies were used: anti-α-tubulin (Sigma, T6199; 1:500), anti-β-actin (Abcam, ab8227; 1:500), anti-acetyl-α-tubulin (Lys40) (Cell Signaling, #5335; 1:500), anti-polyglutamylation (AdipoGen Life Sciences, AG-25B-0030; 1:500), anti-tyrosinated tubulin (YL1/2) (Abcam, ab6160; 1:500), anti-tubulin D1 (Abcam, ab214216; 1:500), anti-SPACA9 (Atlas Antibodies, HPA022243; 1:500), anti-SAXO2 (Atlas Antibodies, HPA040487; 1:500). The following secondary antibodies were used: anti-mouse AlexaFluor488 (Invitrogen, A11029; 1:1,000), anti-rabbit AlexaFluor 488 (Invitrogen, A11034; 1:1,000), anti-mouse AlexaFluor 555 (Invitrogen, A31570; 1:1,000), anti-rabbit AlexaFluor 568 (Invitrogen, A11036; 1:1,000).

### Mass spectrometry proteomics of isolated spermatid microtubules

This *Xenopus* spermatid microtubules isolation protocol was inspired by the published rat manchette isolation method ^26^. Spermatids from 12 *Xenopus laevis* testes were purified, as described above, using MMR supplemented with 2.5 mM EGTA, 5 mM DTT, 10 μg/mL (final) of each leupeptin, pepstatin, and chymostatin (Millipore) protease inhibitors, and 0.5% DMSO, instead of regular MMR, starting from the testis homogenization step. Pelleted cells were resuspended in 500 µL MMR^+^ supplemented with 1.8% Triton X-100. A volume of 480 µL of this cell suspension was mixed with 3.52 mL of 2.5 M sucrose (final mix reaching 2.2 M sucrose), dropped in a centrifugation tube (Beckman, ultraclear, thin wall, #344057). Two sucrose layers were then added on top of the cell suspension: first, 640 µl of 2.05 M sucrose, and then, 320 µL 1 M sucrose on top. After centrifugation at 85,000 g, 16 °C, 110 min in a SW 55 Ti rotor, the spermatid microtubule-containing fraction located at the first interface was collected. A 20 µL-sample was taken aside to control dissociation of manchette-like structures from cell nuclei, using immunofluorescence. The remaining 1-2 mL volume was diluted 10 times in PBS, centrifuged at 10,000 g for 10 min at 16 °C. Supernatant was removed and the pellet was snap frozen in liquid nitrogen.

Pellets were processed using the PreOmics iST kit (PreOmics GmbH, Planegg, Germany), following the manufacturer’s instructions. Briefly, the samples were thawed and lysed (denatured, reduced and alkylated) for 10 min at 95 °C, and then digested with Trypsin/LysC for 3 h at 37 °C. Peptide purification was then carried out at room temperature using a spin cartridge, after which the peptides were eluted in 10 μL of LC-load buffer. Peptide concentrations were determined by measuring the absorbance at 205 nm using a NanoDrop Eight Spectrophotometer. Approximately 350 ng of tryptic peptide samples were separated onto a 75 μm × 250 mm IonOpticks Aurora 3 column (Ion Opticks Pty Ltd., Australia) packed with a 120 A pore, 1.7 μm particle size C18 beads. A reversed-phase gradient of basic buffers (buffer A: 0.1% formic acid, 98% H2O Milli-Q, 2% acetonitrile; buffer B: 0.1% formic acid, 100% acetonitrile) was run on a NanoElute high-performance liquid chromatography (HPLC) system (Bruker Daltonik) at a flow rate of 250 nL/min at 50 °C. The LC run lasted for 80 min with a starting concentration of 2% buffer B increasing to 13% over the first 42 min was first performed, and buffer B concentrations were increased up to 20% at 65 min; 30% at 70 min; 85% at 75 min, and finally 85% for 5 min to wash the column. The NanoElute HPLC system was coupled online to a Tims time-of-flight (TOF) Pro mass spectrometer (timsTOF Pro; Bruker Daltonik) with a CaptiveSpray ion source (Bruker Daltonik). The CaptiveSpray nanoflow electrospray (ESI) source was directly attached to a vacuum inlet capillary via a short capillary extension heated using the instrument’s drying gas. High voltage for the ESI process was applied to the vacuum capillary inlet, whereas the sprayer was kept at ground. The temperature of the ion transfer capillary was set at 180 °C. The spray type was automatically mechanically aligned on the axis with the capillary inlet without the need for any adjustment. Ions were accumulated for 100 ms, and mobility separation was achieved by ramping the entrance potential from −160 to −20 V within 114 ms. The acquisition of the MS mass spectra with the trapped ion mobility spectrometry (TIMS) TOF Pro was done with an average resolution of 60,000 fwhm (mass range 100− 1700 m/z). To enable the PASEF method, precursor m/z and mobility information was first derived from full scan TIMS-MS experiments, MS and MS/MS data were collected over the m/z range 254.1 - 1200 and over the mobility range from 1/K0 = 0.75 to 1/K0 = 1.33 Vs cm^-2^. Resulting quadrupole mass, collision energy and switching times were automatically transferred to the instrument controller as a function of the total cycle time. The quadrupole isolation width was set to 2 and 3 Th and, for fragmentation, the collision energies varied between 20 and 59 eV depending on precursor mass and charge. TIMS, MS operation and PASEF were controlled and synchronized using the control instrument software timsControl 6.0 (Bruker Daltonik). LC-MS/MS data were acquired using the PASEF method with a total cycle time of 1.17 s, including 1 TIMS MS scan and 10 PASEF MS/MS scans. The 10 PASEF scans (100 ms each) contain on average 20 MS/MS scans per PASEF scan. In addition, the most abundant precursors which could have been sequenced in previous scan cycles are dynamically excluded from resequencing. The acquisition of the MS/MS mass spectra with the TIMS TOF Pro is also done with an average resolution of 50,000 FWHM (mass range 100-1700 m/z). Ion mobility resolved mass spectra, nested ion mobility vs m/z distributions, as well as summed fragment ion intensities were extracted from the raw data file with DataAnalysis 6.0 (Bruker Daltonik). Signal-to-noise (S/N) ratios were increased by summations of individual TIMS scans. Mobility peak positions and peak half-widths were determined based on extracted ion mobilograms (± 0.05 Da) using the peak detection algorithm implemented in the DataAnalysis software. Feature detection was also performed using DataAnalysis 6.0 software and exported in .mgf format. Peptide and protein identification were performed using the Mascot database search engine (Mascot server v2.6.2; http://www.matrixscience.com) and its automatic decoy database search to calculate a false discovery rate (FDR). MS/MS spectra were queried against *Xenopus laevis* 10.1 reference proteome UniProt ID: UP000186698 (https://download.xenbase.org/xenbase/Proteomes/; 2023-01-23 release) and a common proteomic contaminant database from the Max Planck Institute of Biochemistry, Martinsried. Mass tolerance for MS and MS/MS was set at 15 ppm and 0.05 Da. The enzyme selectivity was set to trypsin with one miscleavage allowed. Protein modifications were fixed carbamidomethylation of cysteines, variable oxidation of methionine, variable acetylation of the N-terminal of proteins and variable deamidation of asparagine and glutamine. Identification results from Mascot (.dat files) were imported into the Proline Studio software (v2.2.0) ^63^, which was then used for validation and spectral counting, as previously described ^64^. Identified peptides were validated with a peptide rank of 1 and filtered based on Mascot score values to obtain a false discovery rate (FDR) of 1% at the PSM level.

The 200 proteins with the best score average between the 2 biological replicates were subjected to an Overrepresentation Test (Released 20240807) using PANTHER version 19.0 Released 2024-06-20 ^65,66^ against the *Xenopus laevis* (all genes) reference list and the following parameters: GO-Slim Cellular Component annotations, Fisher’s Exact test and Bonferroni correction for multiple testing. From significantly enriched GO-terms, relevant non-redundant cellular components with highest fold-enrichment were plotted with corresponding proteins as a Sankey diagram. The diagram was generated using R and the NetworkD3 package (doi: 10.32614/CRAN.package.networkD3).

### Cryo-electron microscopy of intact spermatids

For cryo-fixation, 3 µl of 1X MMR was applied at the surface of a glow-discharged holey carbon grid (Quantifoil R2/2, Cu200) in the temperature (23 °C) and humidity-controlled atmosphere (∼95%) of an automatic plunge-freezer (EM GP, Leica). Then, 3 µl of spermatid-enriched cell suspension (see above) supplemented with Hoechst (1:1,000) was deposited on the opposite side of the grid. The grid was finally blotted from the opposite side of the sample with the EM GP for 2-3 s using Whatman grade 4 filter paper and plunged into liquid ethane. Specimen grids were transferred to a dual-grid cryo-transfer holder model 205 (Simple Origin) and were observed using a 200 kV electron microscope equipped with a LaB6 filament (Tecnai G2 T20 Sphera, FEI) and a 4K × 4K CMOS camera (XF416, TVIPS).

### Preparation and cryo-electron tomography of lamellae

Cryo-EM grids of intact spermatids were prepared as above. Lamellae of isolated spermatids were produced and acquired at Central European Institute of Technology (CEITEC) at Masaryk University (Brno, Czech Republic). Cellular lamellae were prepared in an Arctis Dual-Beam cryo-plasma focus ion beam (PFIB) (Thermo Fisher Scientific) equipped with an integrated fluorescence microscope module (iFLM). A conductive platinum layer was sputtered onto the grid for 120 seconds at 70 nA and 12 kV Xenon plasma to minimize charging effects during lamella preparation. Additionally, a platinum layer with a thickness of approximately 700 nm was deposited over a period of 2 min utilizing a gas injection system (GIS). Lamellae were prepared using automated lamella workflow with WebUI 1.3 software (Thermo Fisher Scientific). Scanning electron microscopy (SEM) tile set of usable area of TEM grid was collected at 0° tilt in 25 pA and 2 kV and potential lamella sites were identified for further inspection in iFLM. After automated collection of SEM images and optical Z-stack in 388 nm and 470 nm fluorescent channels (Z-stack depth was 10 µm, step 1 µm), SEM images in potential lamella sites were compared with the fluorescence images and placed FIB milling patterns onto the region of suitable fluorescent signal. Each lamella was thinned stepwise at a milling angle of 15° (-23° stage tilt angle) with gradually decreasing beam currents (from 1 nA to 30 pA) until the desired final thickness of 200 nm was reached. The process ended with a polishing step of the lamellae with 10-30 pA at 30 kV.

Tomographic tilt series of lamellae were collected using a Thermo Fisher Scientific Titan Krios transmission electron microscope operating at 300 kV. Data were collected using a Gatan K3 BioContinuum direct electron detector, post-GIF, operating in zero-loss imaging mode, with the energy-selecting slit width set to 10 eV. First, the grid was screened; then, maps of individual lamellae were created, and positions of interest were localized on the lamellae. Cryo-ET data collection was then performed at these positions. The tilt series were collected using SerialEM software ^67^ with a dose-symmetric tilt scheme ^68^ and an angular range of ± 50° (2.5° increment). The total electron exposure was 80 e^-^/Å². Individual images were saved as 4-frame movies in counting mode. Data were collected at a magnification of 33,000 ×, corresponding to a physical pixel size of 2.68 Å, with a target defocus of -5 μm.

### Cryo-electron tomography of microtubules released from spermatids

For cryo-fixation, 4 µl of spermatid-enriched cell suspension (see above) was extemporaneously mixed with 4 µl of nanoparticle buffer (1X MMR, 200 nM mix-matrix capped gold nanoparticles ^69,70^, and 0.2% Triton X-100, added at the last minute), and quickly mixed by pipetting up and down using a cut tip. A volume of 4 µl of this solution was deposited at the surface of a glow-discharged holey carbon grid (Quantifoil R2/2, Cu200) in the temperature (23 °C) and humidity-controlled atmosphere (∼95 %) of an automatic plunge-freezer (EM GP, Leica). After a wait of 10 s, the grid was blotted from the sample side with the EM GP for 1-2 s using Whatman grade 4 filter paper and plunged into liquid ethane. Specimen grids were transferred to a dual-grid cryo-holder model 205 (Simple Origin) or to a rotating cryo-holder CT3500TR (Gatan) to acquire single-and dual-axis tilt series, respectively ^71^, and were observed using a 200 kV electron microscope equipped with a LaB_6_ filament (Tecnai G^2^ T20 Sphera, FEI). Data were acquired using the SerialEM software ^67^ on a 4K × 4K CMOS camera (XF416, TVIPS) at a magnification of 29,000 × (pixel size 3.67 Å) in binning mode 1 or 2. For single-axis tilt series, 40 images were typically taken with 3 degrees increment in an angular range of ∼ ± 60° starting from 0°. For dual-axis tilt series, 31 images per tilt series were typically taken with 3.5 degrees increment in an angular range of ∼ ± 54° starting from 0°. Defocus values were set between -3 and -9 µm. The electron dose was kept constant to ∼0.3 e^-^/Å^2^ per image.

### Tomogram reconstruction and modeling

Tomograms were reconstructed using the Etomo graphical user interface of the IMOD program ^72^. Tilt series were typically filtered after alignment using a low-pass filter. Tomograms were reconstructed in 3D using the SIRT-like filter of Etomo with 15 equivalent iterations. Cryo-electron tomograms were converted to bytes before further processing. Microtubules were then modeled from cryo-tomograms using the 3dmod software.

Tomograms of lamellae were segmented independently for membranes using TARDIS ^73^ and microtubules using 3dmod. The instance.csv files generated from TARDIS were converted into text format and then into IMOD models using the “point2model” command. The membrane and microtubule models were merged using the ’imodjoin’ command, and then visualized in 3dmod superposed on the original tomograms.

### Subtomogram averaging

Subtomogram averages were calculated as previously described ^74^. Briefly, a first model was created by following individual protofilaments in cross section using the slicer tool in IMOD. Usually, ∼50 electronic slices were averaged to reinforce the contrast. A second model was next extrapolated from the first one to mark the microtubule center at the same point positions. Then, a third model was calculated from the previous ones with points spaced every ∼8 nm, and a motive list containing Euler angles of each sub-volume with respect to the chosen reference was created. Sub-volumes of ∼40 pixels^3^ were extracted at each point position using the graphical user interface of the PEET program ^75^. Inner and outer cylindrical masks were used to isolate the microtubule wall densities. Registration of the microtubule sub-volumes was performed by cross-correlation, limiting rotational angular searches around the microtubule axis to about half the angular separation between protofilaments. Other angles were set to take into account variations of microtubule curvature in the X, Y, and Z directions.

### Cryo-electron microscopy data collection for high-resolution analysis of microtubules released from spermatids

For cryo-fixation, 4 µl of spermatid-enriched cell suspension (see above) was extemporaneously mixed with an equal volume of 1X MMR supplemented with 0.2% Triton X-100 and quickly mixed by pipetting up and down using a cut tip. A volume of 4 µl of this solution was deposited at the surface of a glow-discharged holey carbon grid (Quantifoil R2/2, Cu200) in the temperature (23 °C) and humidity-controlled atmosphere (∼95 %) of an automatic plunge-freezer (EM GP, Leica). After a wait of 10 s, the grid was blotted from the sample side with the EM GP for 1-2 s using Whatman grade 4 filter paper and plunged into liquid ethane. Grids were next imaged at the Netherlands Centre for Electron Nanoscopy (NeCEN), Leiden University. Data were collected at 300 kV on a Titan Krios transmission electron microscope (NeCEN Krios 1) equipped with a Gatan K3 direct electron detector and a BioQuantum energy filter. Acquisition was controlled using EPU, with the energy-selecting slit width set to 20 eV. A total dose of 50 e⁻/Å² per movie was fractionated over 50 frames during a 1.53 s exposure (Supplementary Table 3). Targeting was done manually in EPU, selecting areas to be collected around basket-like structures which appeared to contain microtubules (Supplementary Fig.3a). At each of three time points - the beginning, midpoint, and end of the session - 100 images of a cross-grating grid were acquired to estimate magnification anisotropy using mag_distortion_estimate v1.0.1 ^76^. The estimated major-axis scale, minor-axis scale, and distortion angle were 1.0102, 1.0000, and 54.3°, respectively. These parameters were used for frame alignment in MotionCor2 v1.6.4 ^77^, yielding stretch-corrected images with a pixel size of 0.832 Å instead of 0.836 Å. Dose-weighted and non-dose-weighted aligned averages were generated for subsequent particle processing and CTF estimation, respectively. Due to the sample heterogeneity in terms of ice thickness and microtubule density, micrographs were reviewed manually post collection. Of the 37,945 images collected, 25,840 were discarded as they had no microtubule or the sample was too dense/thick to see molecular structures. The 12,105 remaining images were then manually picked to trace filaments, leading to a set of 9,595 usable images containing 20,832 picked filaments. Examples of micrographs illustrating the sample diversity are provided in Supplementary Fig.3b-h.

### Analysis of microtubule lattice parameters

Microtubules are defined by their protofilament (*N*) and monomer helix-start (*S*) numbers and skew angle (*θ*). To determine lattice parameters, images obtained by cryo-electron microscopy were analyzed using TubuleJ – a plugin of ImageJ software developed previously ^31^. Briefly, the microtubule was digitally straightened and centered from the raw image. The Fourier transform was filtered on the *J_0_* and *J_N_* near equatorial terms, resulting in a filtered image showing its fringe pattern. Fuzzy regions correspond to areas of the microtubule wall where protofilaments are intercalated in projection. Dark fringes correspond to regions where protofilaments from the front and back sides overlap in projection. The resulting moiré pattern allows determination of the protofilament number *N*, as previously described ^78^. The period of the moiré pattern (*L*) reflects the protofilament skew angle (*θ*), calculated as *θ = sin*⁻*¹(*δ*x/L),* where δ*x* is the protofilament separation (51.4 Å). The Fourier transform of each raw image converts spatial information into frequency components. Layer-lines observed in the power spectra reflect regular periodicities of the tubulin helix (at ∼1/4 nm^⁻¹^) and the internal proteins (at ∼1/8 nm^⁻¹^). The shift between the 1/4 nm^⁻¹^ lines reflects the sign of the skew angle (*θ*), as previously described ^79^ (Supplementary Fig. 4). Stack of particle-images extracted from straight fibers were created, allowing basic 3D reconstructions with TubuleJ and were visualized with UCSF ChimeraX ^80–82^.

### Generation of 2D microtubule references with the Lattice Accommodation Model

Microtubule references were built from the α-tubulin monomer of the PDB 6WWS model, and the lattice accommodation model parameters ^32^ for different protofilament numbers (*N*), helical-start numbers (*S*), and protofilament skew angles (θ), by varying the helical rise (*r*) of the subunits with respect to the protofilament axis. The subunit spacing along protofilaments (*a*) and the separation between protofilaments (δ*x*) were fixed to 40.1 Å and 51.4 Å, respectively. These references were used in the 2D classification approach described in the next paragraph, to determine the percentage of microtubule configurations belonging to each class (Supplementary Table 1).

The rise needed to build references with different values of θ is given by:

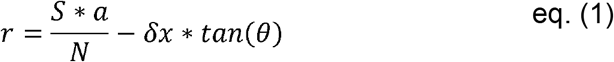

Helical parameters required to generate the references (Supplementary Table 2) were calculated as follow:

**Theta_deg**: angle (°) of the protofilaments with respect to the longitudinal Z-axis of the microtubule.

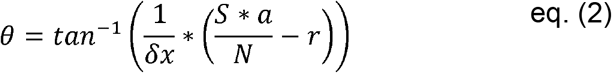

**r_su_A**: rise (Å) of the subunits with respect to the Z-axis.

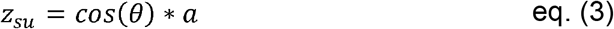

**Phi_su_deg**: rotation angle (°) of the subunits with respect to the Z-axis.

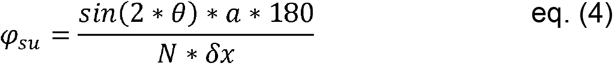

**Delta_C_A**: Radial shift (Å) of the subunits with respect to the initial position of the α-subunit.

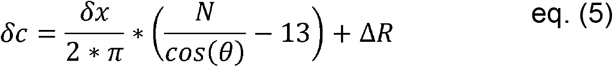

Initial positioning was done using a 13 protofilament microtubule with straight protofilaments (EMDB 5193). A radial shift of 5 Å (Δ*R*) was necessary to avoid steric clashes between tubulin lateral interactions in the references (MT type: 13_3a, Delta_C_A=5.0).

**r_pf_A**: rise (Å) of the protofilaments with respect to the Z-axis.

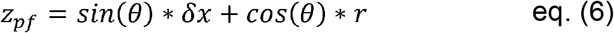

**Phi_pf_deg**: rotation angle (°) of the protofilaments with respect to the Z-axis.

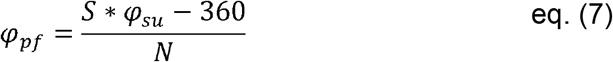

Microtubule models were then generated using UCSF ChimeraX. EMDB 5193 was fetched, the origin index was changed to ’99.5 99.5 59.5’ and the map rotated by -7.7°around the Z-axis to center one protofilament on the Y-axis. PDB 6WWS was fetched and chain K was deleted. The dimer model was placed in the right orientation at the base of the microtubule map. 6WWS was fitted in EMDB 5193, and the chain B was deleted. The remaining α subunit was then tilted by θ (turn 0,1,0 Theta_deg models #2 coordinateSystem #1), a single protofilament (sym #2 h,r_su_A,Phi_su_deg,n,-n/2 coord #1 center 0,Delta_C_A,0 copies true; where ’n’ is the number of subunits/protofilament) and then the microtubule were generated (sym #3 h,r_pf_A,Phi_pf_deg,N coord #1 center 0,Delta_C_A,0 copies true). Finally, the microtubule was centered (move 0,1,0 -Delta_C_A coord #4 models #4). The PDB models were ultimately transformed into densities in ’mrc’ format at the desired resolution using the molmap command in ChimeraX (Supplementary Fig. 5).

TubulePy software can be used to automate these tasks (see Code availability section).

### Cryo-EM data processing for high-resolution analysis of spermatid microtubules

Per micrograph contrast transfer function (CTF) parameters were estimated with Gctf v1.06 ^83^ on non dose-weighed aligned stack averages generated with MotionCor2. 269768 microtubule segments were extracted in Relion ^84^ using 100-pixel boxes at 8 Å per pixel sampled every 160 Å along the microtubule axis. The segments were subjected to reference-based 2D classification in Eman2 ^85^. The reference microtubules were generated as detailed in the previous paragraph. The maps of the references were projected using relion_project with a sampling every 18° for the azimuthal angle and 10° for the out of plane tilt angle (in the range 70-110°). The results of this classification were unusually heterogeneous with filaments assigned to multiple microtubule references along their length. After smoothing of the results to replace isolated outliers by their dominant neighbors, selecting microtubule subsets having at least 4 consecutive segments assigned to the same type led to a set of 92,972 microtubule segments. The distribution of microtubule types assigned to these segments is provided in Supplementary Fig. 6. For 3D reconstruction of microtubule segments, images were resampled to 1.4 Å per pixel in 400-pixel boxes. For the major class of microtubules, which contained 13 protofilaments and displayed the least protofilament skew (13_3a class), 208,920 segments were extracted every 40.1 Å along the microtubule axis, using the in-plane angular assignments refined during the Eman2 2D classification step. Local CTF parameters around the segments were estimated with Gctf v1.06. The segments were subjected to a 3D helical refinement in Relion against a naked microtubule reference, using the 13_3a helical parameters (Supplementary Table 3) and allowing for helical symmetry search. The refinement converged to a map at 4.5 A with a twist of -27.69 degrees and a rise of 9.3 Angstroms and showed substantial heterogeneity in the tubulin subunits. Data were resampled to 4 Å per pixel for 3D classification with alignment (8 classes, tau2_fudge = 4, initial resolution of 15 Å, 100 iterations), imposing the refined helical symmetry (Supplementary Fig. 7a). All resulting 3D classes displayed internal helical-like densities upon symmetry imposition. The most abundant class (Class1, 20% of the particles) was the most homogeneous class with well-resolved tubulin subunits matching what is expected at that pixel sampling (Supplementary Fig.7a). Attempts to align the tubulin subunits within the other classes by 3D refinement were difficult and will require dedicated methodological developments. The analysis was therefore continued with the 42,581 particles from class 1.

3D refinement of the class 1 particles at 1.4 Å per pixel led to a segment resolution of 3.6 Å, which improved to 3.5 Å after CTF refinement (Supplementary Fig. 7b). Up to this stage, the α-and β-tubulin subunits were purposely overlapped because of the 40.1 Å sampling and imposed helical parameters. Processing was then divided into two branches (Supplementary Fig. 7c), using masks that selected either a single short protofilament segment comprising four tubulin subunits or two adjacent short protofilament segments, the latter allowing A-and B-lattice configurations to be reconstructed. The masks extended broadly into the microtubule lumen to retain any potential associated internal protein density. In both branches, particles were symmetry-expanded 13-fold to separate the different protofilament registers, followed by signal subtraction and recentering on masks encompassing either one or two short protofilament segments. The resulting particle images were cropped into 240-pixel boxes and subjected to local 3D refinement in Relion with C1 symmetry, using a mask similar to that used for signal subtraction and the symmetry-expanded poses as priors for the local refinement (healpix_order 5, auto_local_healpix_order 5, sigma_ang 2).

For the one-protofilament branch, particles were subjected to 3D classification without alignment to assess potential heterogeneity in the internal proteins (6 classes, tau2_fudge = 4, initial resolution of 15 Å, 30 iterations). The classification rapidly converged to two approximately equally populated classes, while classes 3–6 each contained less than 0.001% of the particles (Supplementary Fig. 7c). The two major classes displayed the same SPACA9-like internal density at two registers separated by ∼40 Å, with well-separated α-and β-tubulin subunits. As these two registers represented redundant views of the same structure, class 1, in which the internal protein density was centered in the box, was selected for further processing (Supplementary Fig. 7c). Because the internal protein density was weaker than the tubulin signal, focused 3D classification was then performed on the central luminal protein density at 3.5 Å per pixel to classify its occupancy (2 classes, tau2_fudge = 4, initial resolution of 15 Å, 150 iterations). This yielded one class in which the central tubulin was decorated (64% of particles) and one in which it was undecorated (36%). Both classes were reconstructed at 1.4 Å per pixel from unsubtracted particles, yielding maps at 3.3 Å and 3.5 Å resolution, respectively, over three consecutive tubulin dimers. For the decorated class, the resolution reached 3.1 Å over the short protofilament segment decorated with the internal protein, for which local resolution was also calculated (Supplementary Fig. 8a,b). This map was used as the final map for 1 protofilament. It showed well-separated α-and β-tubulin densities (Supplementary Fig. 8c).

For the two-protofilament branch, particles were subjected to 3D classification separately for each protofilament to determine its tubulin register. Based on the results of the one-protofilament classification, three classes were used: two classes corresponding to the two major registers and a third class to accommodate particles that did not fit either register well (tau2_fudge = 4, initial resolution of 15 Å, 50 iterations). For each protofilament, this classification yielded two major, approximately equally populated classes, whereas the third class contained only ∼2% of the particles (Supplementary Fig. 7c). Of the two possible same-register combinations, the one with tubulin dimers centered in the box was selected to reconstruct the B-lattice. Similarly, of the two possible alternating-register combinations, one was selected to reconstruct the A-lattice (seam), as they represented redundant views. Both lattice configurations were initially reconstructed at 3.5 Å per pixel (Supplementary Fig. 7c). In each case, a broad mask encompassing the luminal region was then used for focused 3D classification of the internal protein density at this sampling into four classes (B-lattice: tau2_fudge = 4, initial resolution = 15 Å, 15 iterations; A-lattice: tau2_fudge = 8, initial resolution = 15 Å, 50 iterations). For the B-lattice, this classification resolved three occupancy patterns across the protofilament pair (Supplementary Fig. 7c), whereas for the A-lattice it separated decorated and undecorated classes. The resulting classes were reconstructed at 1.4 Å per pixel from unsubtracted particles (Supplementary Fig. 7c). The fully decorated B-lattice and decorated A-lattice reconstructions reached resolutions of 3.2 Å and 4.0 Å, respectively, and were used as the final maps for the respective lattice configurations (Supplementary Fig. 8a,b). Both maps showed well-separated α-and β-tubulin densities (Supplementary Fig. 8c).

Due to the heterogeneous local resolution of the maps (Supplementary Fig. 8b), the final maps were filtered with EMReady ^86^ after post-processing in RELION using a B-factor of −20 Å². For display as meshes in Fig. 5-6, the maps were resampled twofold in Coot ^87^.

### Identification of internal proteins from the high-resolution cryo-EM maps

In the three final cryo-EM maps corresponding to the decorated one-protofilament, B-lattice and A-lattice reconstructions, the internal protein density comprised a folded, predominantly α-helical domain and an approximately 20-residue peptide extending along the tubulin dimers, including a short helix positioned at the α/β-tubulin intradimer interface (Fig. 5a,b). This architecture closely resembled the previously described SPACA9–SAXO human complex ^33,34^. Protein identities were first assessed using ModelAngelo ^35^, which readily assigned the larger of the two internal densities to Spaca9 but did not identify the smaller peptide density. The Spaca9 assignment was further supported by its distinctive fold (Fig. 5d). The Saxo model from a previously determined SPACA9–SAXO complex ^33^ closely matched the position and trajectory of the peptide density. SAXO proteins contain repeated microtubule-binding Mn motifs, homologous to the Mn modules originally described in MAP6-family proteins ^36,37^. Structural studies have shown that Mn motifs form short helices at the α/β-tubulin intradimer interface ^38^. This strongly suggested that the peptide originated from a protein containing an Mn motif. However, the limited number of resolved residues and the local resolution of the map precluded direct identification from the cryo-EM density alone. To address this limitation, we integrated information from the cryo-EM map with mass spectrometry and AlphaFold3 analyses. We first used the cryo-EM density to define a likely sequence motif over a 10-residue window: position 3, F/Y/H; position 4, R/K/F/Y/H/L/I/M/Q; position 7, Y/F/H; and position 9, P. This motif was then complemented using the two conserved residue positions characteristic of Mn motifs ^38^: a T/S constraint was added at position 1, whereas the second conserved position was already satisfied by the aromatic residue at position 7.

A strict search for this motif within 10-residue windows across the 1,634 protein sequences identified by mass spectrometry yielded 19 candidates (Supplementary Fig. 9a). For each candidate, protein regions extending up to 500 residues around the region containing the matching motif(s), where sequence length permitted, were selected for AlphaFold3 analysis. The most abundant α-and β-tubulin isotypes identified by mass spectrometry (Supplementary Table 4), Tuba5.S (UniProt accession code: Q7ZTP0) and Tubb4b.L (UniProt accession code: P30883), respectively, were used for this analysis. Two candidates contained multiple compatible sites within their sequences: Saxo2.L (Uniprot accession code: Q6DCB9), with 11 potential sites, and Saxo4.L, with four potential sites. Fifteen models with an inter-chain predicted aligned error (inter-PAE) greater than 10 Å were excluded including Saxo4.L, and the remaining four models were evaluated using the iQ-score metric ^88^ (Supplementary Fig. 9a). The predicted structures of these candidates relative to the tubulin dimer are shown in Supplementary Fig. 9b. Candidate interface residues were defined as residue pairs with a center-of-mass distance below 10 Å and an inter-PAE below 10 Å ^88,89^.

Among the four candidates, only Saxo2.L exhibited predicted interfaces on the luminal side of the microtubule (Supplementary Fig. 9b; Supplementary Fig. 10a,b), consistent with the experimentally observed cryo-EM density, and multiple regions of Saxo2.L satisfied both the Mn-motif and cryo-EM side-chain density constraints. To further refine the localization of the interaction sites, Saxo2.L was segmented into 51-residue sequence fragments centered on the matching motifs, allowing at most one mismatched motif position or satisfying the Mn-motif criterion (defined as T/S at position 1 and F/Y at position 7), and the same AlphaFold3 modelling and evaluation pipeline was applied to each of the 16 resulting fragments, named after their central residue number (Supplementary Fig. 9c,d). Nine fragments were predicted to interact with α-and β-tubulin. However, the Saxo2.L_141 fragment, which does not contain the complete Mn motif, bound at a position inconsistent with the experimentally observed cryo-EM density and was therefore excluded, leaving eight candidate fragments (Supplementary Fig. 9c,d; Supplementary Fig. 10c,d).

Although Saxo4.L was not retained in the previous analysis, it was the other SAXO protein detected by mass spectrometry and the only other SAXO protein present in the *Xenopus laevis* proteome used for this analysis. We therefore applied the same fragment-based analysis to the ten 51-residue Saxo4.L sequences partially matching the motif as done for Saxo2.L. None showed a predicted tubulin interaction supported by the coevolutionary signal. Saxo2.L was also consistently more abundant than Saxo4.L across the two mass-spectrometry replicates, as assessed by emPAI and specific spectral counts (Supplementary Table 5). At the residue-interaction level, all eight retained Saxo2.L fragments engaged the same tubulin interface (Supplementary Fig.9c,d; Supplementary Fig. 10c,d) and were collectively compatible with the cryo-EM density (Fig. 5e), suggesting that the observed density likely represents mainly a mixture of several Saxo2.L Mn motifs binding at the same site. Saxo2.L and Spaca9.L are hereafter referred to as Saxo2 and Spaca9, respectively.

### High-resolution cryo-EM model building and resolution estimations

Model building was initiated using the single-protofilament reconstruction and the bovine sperm tubulin-SAXO1-SPACA9 model from PDB 8OU0 ^34^ as an initial template. Individual chains extracted from 8OU0 were fitted into the cryo-EM density in ChimeraX ^82^ and remodeled to the corresponding *Xenopus laevis* sequences using MODELLER ^90^ (UniProt accession codes: Spaca9.L, A0A8J0PW71; Saxo2.L, Q6DCB9; Tuba5.S, Q7ZTP0; Tubb4b.L, P30883). For Saxo2.L, the candidate SAXO-binding motif spanning residues 199– 218 (Saxo2.L_209 in Fig. 5) was used for modelling. The resulting model was locally fitted to the density and iteratively refined in Coot ^91^. This model was subsequently used to populate the decorated B-lattice and A-lattice reconstructions, and the individual chains were then manually adjusted and refined in Coot to account for differences between the maps. Finally, all three atomic models were refined using Phenix real-space refinement ^92^, retaining four tubulin subunits per protofilament in the refinement models. Atomic-model and cryo-EM density figures were prepared using ChimeraX.

Final global resolutions were estimated from Fourier shell correlation (FSC) curves using the FSC = 0.143 criterion (Supplementary Fig. 8a), generated with RELION PostProcess. Local-resolution estimates were also calculated in RELION (Supplementary Fig. 8b). For global resolution estimation, FSC curves were calculated between the two independently refined half-maps (gold-standard refinement) using soft masks generated with relion_mask_create from densities calculated from the refined atomic models (low-pass filter, 15 Å; soft edge, 5 pixels). The cryo-EM density maps and atomic coordinates (Supplementary Table 3) are deposited to the Electron Microscopy Data Bank and Protein Data Bank, respectively, with the following accession numbers: EMDB-59871, PDB 34IK (1 protofilament-decorated), EMDB-59872, PDB 34IL (B-lattice-decorated), and EMDB-59873, PDB 34IM (A-lattice-decorated).

### RNA-FISH

RNA-FISH was performed according to the HCR™ v3.0 system (Molecular Instruments, Inc) protocol with probes against *mybl1.L* (NM_001087710), *spaca9.L* (NM_001091855) and *saxo2.L* (NM_001093710) mRNAs produced by the manufacturer and proprietary to them. Briefly, testes from adult *Xenopus laevis* male were fixed in 4% PFA in PBS for 30 min at room temperature. After 3 washes with PBS, tissues were permeabilized for 1 h with 70% Ethanol at room temperature, then stored at -20 °C in 70% Ethanol until the experiment. The day of the experiment, testes lobules were dissected and isolated using forceps and transferred to 1 mL of Wash Buffer (pre-warmed at 37 °C) and incubated for 10 min. The Wash Buffer was then replaced by 500 µL of Hybridization Buffer (pre-warmed at 37 °C) and incubated for 10 min. Finally, the Hybridization Buffer was replaced by 500 µL of probe solution (500 µL Hybridization Buffer supplemented with 3 µL of 1 µM probe stock solution, pre-warmed at 37 °C) and incubated overnight at 37 °C. The probe solution was discarded and the lobules washed 2 times for 20 min with 1 mL of Wash Buffer (pre-warmed at 37 °C) and then 2 times for 20 min with 5X SSC (prepared from 20X SSC commercial solution, diluted using Milli-Q water) supplemented with 10% Tween 20. Volumes of 4 µL of both H1 and H2 3 µM hairpin stocks were heated at 95 °C for 90 s and left to cool at room temperature in dark. The SSC solution was discarded, replaced by 1 mL of Amplification Buffer and incubated for 10 min on a rotating wheel for 10 min. To prepare the hairpin solution, the two heated 4 µL of hairpins were mixed together in 500 µL of Amplification Buffer and incubated for 5 min at room temperature. The Amplification Buffer was discarded, replaced by the hairpin solution and incubated overnight at room temperature in the dark on a rotating wheel. The hairpin solution was then discarded and lobules were washed 2 times for 20 min with 5X SSC supplemented with 10% Tween 20, the second wash containing 10 μg/mL of Hoechst. Lobules were finally mounted one by one in cavity slides with one drop of Prolong Gold antifade reagent (Invitrogen #P36 934) and covered with a coverslip. Z-stacks of 19 or 20 slices, separated by 41.22 or 43.51 µm, respectively, were acquired on a Zeiss LSM 900 Axio Observer 7 confocal microscope using the Zen software and a LD Plan-Neofluar 20x/0.4 Korr M27 objective. Images are mean averages of eight bidirectional scans with a depth of 16 bits. Pinhole size was chosen to correspond to 1 Airy unit. Images were processed using the Zeiss LSM plus module and then analyzed using Fiji.

### Single-cell RNA-seq data analysis

Liao et al., 2022 raw *Xenopus laevis* scRNA-seq data ^41^ from two adult testis samples was retrieved from GEO repository through accession number GSE195790 and aligned against *Xenopus laevis* 10.1 reference genome (https://download.xenbase.org/xenbase/Genomics/JGI/Xenla10.1/2023-07-06) provided by Xenbase (http://www.xenbase.org/, RRID:SCR_003280) ^93^. Samples were integrated and batch corrected using RunFastMNN function from SeuratWrappers (v0.3.1) ^94^. Cells passing the following quality controls were kept for further analysis: 200 < mRNA counts < 40,000; unique genes detected < 6,000 and fraction of mitochondrial genes < 10%. Finally, violin plots were generated using VlnPlot function from Seurat v4 ^95^.

## Code availability

TubuleJ software is available at https://github.com/dchretien35/TubuleJ

TubulePy software is available at https://github.com/dchretien35/TubulePy.

AlphaFold screening software and scoring are available at https://github.com/Qrouger/PPIFold and https://github.com/Qrouger/HInt.

## Acknowledgments

We thank members of the Gibeaux team (MiToS), past and present, for support and fruitful discussions. We are grateful to Julien Maurais at the IGDR *Xenopus* facility for dedicated frog care. This work was supported by the ANR (JCJC 22-CE11-0010), which provided the postdoctoral salary of M.P.M.H.B., and by a 2024 Scientific Challenges grant from the University of Rennes, Research Division of the Research and Innovation Directorate (DRI). K.M. was supported by France 2030 (ANR-22-PAMR-0005). We thank Biosit core facilities (UAR 3480 US_S 018) H2P2, Protim and TEM2C. H2P2 facility is supported by France- BioImaging (ANR-10-INBS-04). Protim facility is supported by grants from Biogenouest, Infrastructures en Biologie Santé et Agronomie (IBiSA), and Conseil Régional de Bretagne awarded to C.P. The TEM2C facility has received funding from the European Union through the CPER B2S (2021–2027), co-funded by the French Ministry of Higher Education and Research, the Brittany Region, the Ille-et-Vilaine Department, Rennes Métropole, and the CNRS, as well as from additional grants from IBiSA and Biogenouest. This work benefited from access to the NeCEN electron microscopy facility, an Instruct-ERIC center. We thank Willem Noteborn at NeCEN for microscope alignment and set up of the data collection parameters. Financial support was provided by European EC Horizon2020 iNEXT (Project 653706), iNEXT-Discovery (PID: 30546). We acknowledge the CEITEC Cryo-electron Microscopy and Tomography facility of CIISB, Instruct-CZ Centre, supported by MEYS CR (LM2023042) and European Regional Development Fund-Project *“*Innovation of Czech Infrastructure for Integrative Structural Biology” *(No.* CZ.02.01.01/00/23_015/0008175), and are particularly grateful to Jiří Nováček, Zuzana Hlavenková and Jana Moravcová. Financial support was provided by Instruct-ERIC (PID: 39231) and through the EDUC-WIDE program, supported by the European Union under Grant Agreement No. 101136533.

## Author contributions

R.G. and D.C. designed and supervised the project. C.C. performed the molecular and cell biology, as well as the low-resolution electron microscopy experiments with D.C., and prepared the samples for high-resolution cryo-EM. M.P.M.H.B. performed the high-resolution cryo-EM data collection, designed and performed the single-particle analysis, and carried out the modelling and maps analysis. Q.R. performed the AlphaFold3 based analyses to identify the Saxo candidates, under the supervision of K.M. F.B. performed the RNA-FISH experiments and analyzed the single-cell RNA-seq data, supervised by J.J. R.V. processed the testis samples for histology analysis. C.H. supported all wet-lab aspects of the work, C.G. performed critical cryo-fixations, and L.D. prepared the gold nanoparticles essential for cryo12 electron tomography. B.G. prepared and analyzed the protein samples and R.L. optimized and performed the mass spectrometry acquisitions, supervised by E.C. and C.P. C.C., M.P.M.H.B, Q.R., R.G. and D.C. prepared the figures. R.G., C.C. and M.P.M.H.B. wrote the manuscript, with help from all authors.

## References

1. Phillips, D. M. Nuclear shaping in the absence of microtubules in scorpion spermatids. The Journal of Cell Biology 62, 911–917 (1974).

2. Martianov, I., et al. Polar nuclear localization of H1T2, a histone H1 variant, required for spermatid elongation and DNA condensation during spermiogenesis. Proc. Natl. Acad. Sci. U.S.A. 102, 2808–2813 (2005).

3. Asa, C. S. & Phillips, D. M. Nuclear shaping in spermatids of the Thai leaf frog *Megophrys* montana. Anat. Rec. 220, 287–290 (1988).

4. Kierszenbaum, A. L. & Tres, L. L. The acrosome-acroplaxome-manchette complex and the shaping of the spermatid head. Arch. Histol. Cytol. 67, 271–284 (2004).

5. Kierszenbaum, A. L., Rivkin, E. & Tres, L. L. Molecular biology of sperm head shaping. Soc Reprod Fertil Suppl 65, 33–43 (2007).

6. Rattner, J. B. & Brinkley, B. R. Ultrastructure of mammalian spermiogenesis. Journal of Ultrastructure Research 41, 209–218 (1972).

7. Dunleavy, J. E. M., O’Bryan, M. K., Stanton, P. G. & O’Donnell, L. The cytoskeleton in spermatogenesis. Reproduction 157, R53–R72 (2019).

8. Kierszenbaum, A. L., Rivkin, E. & Tres, L. L. Cytoskeletal track selection during cargo transport in spermatids is relevant to male fertility. Spermatogenesis 1, 221–230 (2011).

9. Mochida, K., Rivkin, E., Gil, M. & Kierszenbaum, A. L. Keratin 9 Is a Component of the Perinuclear Ring of the Manchette of Rat Spermatids. Developmental Biology 227, 510– 519 (2000).

10. Kato, A., Nagata, Y. & Todokoro, K. δ-Tubulin is a component of intercellular bridges and both the early and mature perinuclear rings during spermatogenesis. Developmental Biology 269, 196–205 (2004).

11. Dunleavy, J. E. M. et al. Katanin-like 2 (KATNAL2) functions in multiple aspects of haploid male germ cell development in the mouse. PLoS Genet 13, e1007078 (2017).

12. Stathatos, G. G., Dunleavy, J. E. M., Zenker, J. & O’Bryan, M. K. Delta and epsilon tubulin in mammalian development. Trends in Cell Biology 31, 774–787 (2021).

13. Akhmanova, A. et al. The microtubule plus-end-tracking protein CLIP-170 associates with the spermatid manchette and is essential for spermatogenesis. Genes Dev. 19, 2501–2515 (2005).

14. Lehti, M. S. & Sironen, A. Formation and function of the manchette and flagellum during spermatogenesis. REPRODUCTION 151, R43–R54 (2016).

15. Russell, L. D., Russell, J. A., MacGregor, G. R. & Meistrich, M. L. Linkage of manchette microtubules to the nuclear envelope and observations of the role of the manchette in nuclear shaping during spermiogenesis in rodents. Am. J. Anat. 192, 97–120 (1991).

16. Yoshida, T., Ioshii, S. O., Imanaka-Yoshida, K. & Izutsu, K. Association of cytoplasmic dynein with manchette microtubules and spermatid nuclear envelope during spermiogenesis in rats. Journal of Cell Science 107, 625–633 (1994).

17. O’Donnell, L. et al. An Essential Role for Katanin p80 and Microtubule Severing in Male Gamete Production. PLoS Genet 8, e1002698 (2012).

18. Mendoza-Lujambio, I. The Hook1 gene is non-functional in the abnormal spermatozoon head shape (azh) mutant mouse. Human Molecular Genetics 11, 1647–1658 (2002).

19. Kitaoka, M., Heald, R. & Gibeaux, R. Spindle assembly in egg extracts of the Marsabit clawed frog, Xenopus borealis. Cytoskeleton 75, 244–257 (2018).

20. Bernardini, G., Podini, P., Maci, R. & Camatini, M. Spermiogenesis in Xenopus laevis: from late spermatids to spermatozoa. Mol Reprod Dev 26, 347–355 (1990).

21. Reed, S. C. & Stanley, H. P. Fine structure of spermatogenesis in the South African clawed toad Xenopus laevis daudin. Journal of Ultrastructure Research 41, 277–295 (1972).

22. Eimanifar, A., Aufderheide, J., Schneider, S. Z., Krueger, H. & Gallagher, S. Development of an in vitro diagnostic method to determine the genotypic sex of *Xenopus laevis*. PeerJ 7, e6886 (2019).

23. Teperek, M. et al. Sperm is epigenetically programmed to regulate gene transcription in embryos. Genome Res. 26, 1034–1046 (2016).

24. Stathatos, G. G. et al. Delta tubulin stabilizes male meiotic kinetochores and aids microtubule remodeling and fertility. Journal of Cell Biology 224, e202412056 (2025).

25. Gadadhar, S., Hirschmugl, T. & Janke, C. The tubulin code in mammalian sperm development and function. Seminars in Cell & Developmental Biology 137, 26–37 (2023).

26. Mochida, K., Tres, L. L. & Kierszenbaum, A. L. Isolation of the Rat Spermatid Manchette and Its Perinuclear Ring. Developmental Biology 200, 46–56 (1998).

27. Hu, W. et al. CAMSAP1 role in orchestrating structure and dynamics of manchette microtubule minus-ends impacts male fertility during spermiogenesis. Proceedings of the National Academy of Sciences 120, e2313787120 (2023).

28. Judernatz, J. H., Pérez Pañeda, L., Kadavá, T., Heck, A. J. & Zeev-Ben-Mordehai, T. Characterisation of the manchette architecture and its role as transport scaffold using cryo-electron tomography. Life Sci. Alliance 8, e202503415 (2025).

29. Pleuger, C., Lehti, M. S., Dunleavy, J. E., Fietz, D. & O’Bryan, M. K. Haploid male germ cells—the Grand Central Station of protein transport. Human Reproduction Update 26, 474–500 (2020).

30. Tres, L. L. & Kierszenbaum, A. L. *Sak57*, an acidic keratin initially present in the spermatid manchette before becoming a component of paraaxonemal structures of the developing tail. Mol. Reprod. Dev. 44, 395–407 (1996).

31. Ku, S., Messaoudi, C., Guyomar, C., Kervrann, C. & Chrétien, D. Determination of Microtubule Lattice Parameters from Cryo-electron Microscope Images Using TubuleJ. Bio Protoc 10, e3814 (2020).

32. Chrétien, D. & Fuller, S. D. Microtubules switch occasionally into unfavorable configurations during elongation. Journal of Molecular Biology 298, 663–676 (2000).

33. Gui, M., et al. SPACA9 is a lumenal protein of human ciliary singlet and doublet microtubules. Proc. Natl. Acad. Sci. U.S.A. 119, e2207605119 (2022).

34. Leung, M. R. et al. Structural specializations of the sperm tail. Cell 186, 2880–2896.e17 (2023).

35. Jamali, K. et al. Automated model building and protein identification in cryo-EM maps. Nature 628, 450–457 (2024).

36. Bosc, C. et al. Identification of Novel Bifunctional Calmodulin-binding and Microtubule-stabilizing Motifs in STOP Proteins. Journal of Biological Chemistry 276, 30904–30913 (2001).

37. Dacheux, D. et al. A MAP6-Related Protein Is Present in Protozoa and Is Involved in Flagellum Motility. PLoS ONE 7, e31344 (2012).

38. Wang, X. et al. Cryo-EM structure of cortical microtubules from human parasite Toxoplasma gondii identifies their microtubule inner proteins. Nat Commun 12, 3065 (2021).

39. Zabeo, D. et al. A lumenal interrupted helix in human sperm tail microtubules. Sci Rep 8, 2727 (2018).

40. Session, A. M. et al. Genome evolution in the allotetraploid frog Xenopus laevis. Nature 538, 336–343 (2016).

41. Liao, Y. et al. Cell landscape of larval and adult Xenopus laevis at single-cell resolution. Nat Commun 13, 4306 (2022).

42. Harding, C. R. & Frischknecht, F. The Riveting Cellular Structures of Apicomplexan Parasites. Trends in Parasitology 36, 979–991 (2020).

43. Lefèvre, J. et al. Structural Basis for the Association of MAP6 Protein with Microtubules and Its Regulation by Calmodulin. Journal of Biological Chemistry 288, 24910–24922 (2013).

44. Dacheux, D. et al. Human FAM154A (SAXO1) is a microtubule-stabilizing protein specific to cilia and related structures. Journal of Cell Science 128, 1294–1307 (2015).

45. Andersen, J. S. et al. Uncovering structural themes across cilia microtubule inner proteins with implications for human cilia function. Nat Commun 15, 2687 (2024).

46. Zhu, Y. et al. In situ structure of the mouse sperm central apparatus reveals mechanistic insights into asthenozoospermia. Cell Res 35, 551–567 (2025).

47. Judernatz, J. H. et al. SPACA9 and MNMIP1 bridge the seam of spermatid manchette microtubules. EMBO J 10.1038/s44318-026-00833-w(2026) doi:10.1038/s44318-026-00833-w.

48. Tai, L., Yin, G., Huang, X., Sun, F. & Zhu, Y. In-cell structural insight into the stability of sperm microtubule doublet. Cell Discov 9, 116 (2023).

49. Aboraya, M., Ben-Uliel, S. F. & Orbach, R. SPACA9 Acts as a Molecular Staple Modulating Microtubule Dynamic Instability.

50. Foster, H. E., Ventura Santos, C. & Carter, A. P. A cryo-ET survey of microtubules and intracellular compartments in mammalian axons. Journal of Cell Biology 221, e202103154 (2021).

51. Tsuji, C. et al. CryoET reveals actin filaments within platelet microtubules. Nat Commun 15, 5967 (2024).

52. Chalfie, M. & Thomson, J. N. Structural and functional diversity in the neuronal microtubules of Caenorhabditis elegans. The Journal of cell biology 93, 15–23 (1982).

53. Renauld, J., Thelen, N., Bartholomé, O., Malgrange, B. & Thiry, M. Dispensability of Tubulin Acetylation for 15-protofilament Microtubule Formation in the Mammalian Cochlea. Cell Struct. Funct. 46, 11–20 (2021).

54. Guyomar, C. et al. Changes in seam number and location induce holes within microtubules assembled from porcine brain tubulin and in Xenopus egg cytoplasmic extracts. eLife 11, e83021 (2022).

55. Russell, L. D. Spermatid-Sertoli tubulobulbar complexes as devices for elimination of cytoplasm from the head region late spermatids of the rat. Anat Rec 194, 233–246 (1979).

56. Upadhyay, R. D., Kumar, A. V., Ganeshan, M. & Balasinor, N. H. Tubulobulbar complex: cytoskeletal remodeling to release spermatozoa. Reprod Biol Endocrinol 10, 27 (2012).

57. Kerr, J. B. & de Kretser, D. M. Proceedings: The role of the Sertoli cell in phagocytosis of the residual bodies of spermatids. J Reprod Fertil 36, 439–440 (1974).

58. Kalt, M. R. Ultrastructural observations on the germ line of Xenopus laevis. Z Zellforsch Mikrosk Anat 138, 41–62 (1973).

59. Bernardini, G., Stipani, R. & Melone, G. The ultrastructure of Xenopus spermatozoon. Journal of Ultrastructure and Molecular Structure Research 94, 188–194 (1986).

60. Schindelin, J. et al. Fiji: an open-source platform for biological-image analysis. Nat Methods 9, 676–682 (2012).

61. Hannak, E. & Heald, R. Investigating mitotic spindle assembly and function in vitro using Xenopus laevis egg extracts. Nat Protoc 1, 2305–2314 (2006).

62. Edelstein, A. D. et al. Advanced methods of microscope control using μManager software. J Biol Methods 1, e10 (2014).

63. Bouyssié, D. et al. Proline: an efficient and user-friendly software suite for large-scale proteomics. Bioinformatics 36, 3148–3155 (2020).

64. Méar, L. et al. The Eutopic Endometrium Proteome in Endometriosis Reveals Candidate Markers and Molecular Mechanisms of Physiopathology. Diagnostics 12, 419 (2022).

65. Thomas, P. D. et al. PANTHER : Making genome-scale phylogenetics accessible to all. Protein Science 31, 8–22 (2022).

66. Mi, H. et al. Protocol Update for large-scale genome and gene function analysis with the PANTHER classification system (v.14.0). Nat Protoc 14, 703–721 (2019).

67. Mastronarde, D. N. Automated electron microscope tomography using robust prediction of specimen movements. Journal of Structural Biology 152, 36–51 (2005).

68. Hagen, W. J. H., Wan, W. & Briggs, J. A. G. Implementation of a cryo-electron tomography tilt-scheme optimized for high resolution subtomogram averaging. Journal of Structural Biology 197, 191–198 (2017).

69. Duchesne, L., Gentili, D., Comes-Franchini, M. & Fernig, D. G. Robust Ligand Shells for Biological Applications of Gold Nanoparticles. Langmuir 24, 13572–13580 (2008).

70. Guesdon, A. et al. EB1 interacts with outwardly curved and straight regions of the microtubule lattice. Nat Cell Biol 18, 1102–1108 (2016).

71. Guesdon, A., Blestel, S., Kervrann, C. & Chrétien, D. Single versus dual-axis cryo-electron tomography of microtubules assembled in vitro: Limits and perspectives. Journal of Structural Biology 181, 169–178 (2013).

72. Kremer, J. R., Mastronarde, D. N. & McIntosh, J. R. Computer Visualization of Three-Dimensional Image Data Using IMOD. Journal of Structural Biology 116, 71–76 (1996).

73. Kiewisz, R. et al. Accurate and fast segmentation of filaments and membranes in micrographs and tomograms with TARDIS. Preprint at 10.1101/2024.12.19.629196 (2024).

74. Bousquet, C., Heumann, J., Chrétien, D. & Guyomar, C. Characterization of Microtubule Lattice Heterogeneity by Segmented Subtomogram Averaging. BIO-PROTOCOL 13, (2023).

75. Nicastro, D. et al. The Molecular Architecture of Axonemes Revealed by Cryoelectron Tomography. Science 313, 944–948 (2006).

76. Grant, T. & Grigorieff, N. Automatic estimation and correction of anisotropic magnification distortion in electron microscopes. Journal of Structural Biology 192, 204– 208 (2015).

77. Zheng, S. Q. et al. MotionCor2 - anisotropic correction of beam-induced motion for improved cryo-electron microscopy. Nat Methods 14, 331–332 (2017).

78. Chrétien, D. & Wade, R. H. The microtubule surface lattice. AIP Conference Proceedings 226, 153–159 (1991).

79. Chrétien, D., Kenney, J. M., Fuller, S. D. & Wade, R. H. Determination of microtubule polarity by cryo-electron microscopy. Structure 4, 1031–1040 (1996).

80. Meng, E. C. et al. UCSF ChimeraX: Tools for structure building and analysis. Protein Sci 32, e4792 (2023).

81. Goddard, T. D. et al. UCSF ChimeraX: Meeting modern challenges in visualization and analysis. Protein Sci 27, 14–25 (2018).

82. Pettersen, E. F. et al. UCSF ChimeraX: Structure visualization for researchers, educators, and developers. Protein Sci 30, 70–82 (2021).

83. Zhang, K. Gctf: Real-time CTF determination and correction. J Struct Biol 193, 1–12 (2016).

84. Zivanov, J. et al. New tools for automated high-resolution cryo-EM structure determination in RELION-3. eLife 7, e42166 (2018).

85. Tang, G. et al. EMAN2: an extensible image processing suite for electron microscopy. J Struct Biol 157, 38–46 (2007).

86. He, J., Li, T. & Huang, S.-Y. Improvement of cryo-EM maps by simultaneous local and non-local deep learning. Nat Commun 14, 3217 (2023).

87. Emsley, P. & Cowtan, K. Coot: model-building tools for molecular graphics. Acta Crystallogr D Biol Crystallogr 60, 2126–2132 (2004).

88. Rouger, Q., Giudice, E., Meyer, D. F. & Macé, K. PPIFold: a tool for analysis of protein-protein interaction from AlphaPullDown. Bioinform Adv 5, vbaf090 (2025).

89. Rouger, Q. et al. HInt: interaction-based homology discovery through accelerated genome-scale AlphaFold screening. Preprint at 10.64898/2026.07.31.741991 (2026).

90. Webb, B. & Sali, A. Comparative Protein Structure Modeling Using MODELLER. Curr Protoc Bioinformatics 54, 5.6.1-5.6.37 (2016).

91. Emsley, P., Lohkamp, B., Scott, W. G. & Cowtan, K. Features and development of Coot. Acta Crystallogr D Biol Crystallogr 66, 486–501 (2010).

92. Afonine, P. V. et al. Real-space refinement in PHENIX for cryo-EM and crystallography. Acta Crystallogr D Struct Biol 74, 531–544 (2018).

93. Fisher, M. et al. Xenbase: key features and resources of the *Xenopus* model organism knowledgebase. GENETICS 224, iyad018 (2023).

94. Haghverdi, L., Lun, A. T. L., Morgan, M. D. & Marioni, J. C. Batch effects in single-cell RNA sequencing data are corrected by matching mutual nearest neighbours. Nat Biotechnol 36, 421–427 (2018).

95. Hao, Y. et al. Integrated analysis of multimodal single-cell data. Cell 184, 3573–3587.e29 (2021).

