## Supplemental Files for "An internal scaffold supports microtubule architectural diversity in *Xenopus* spermatids"

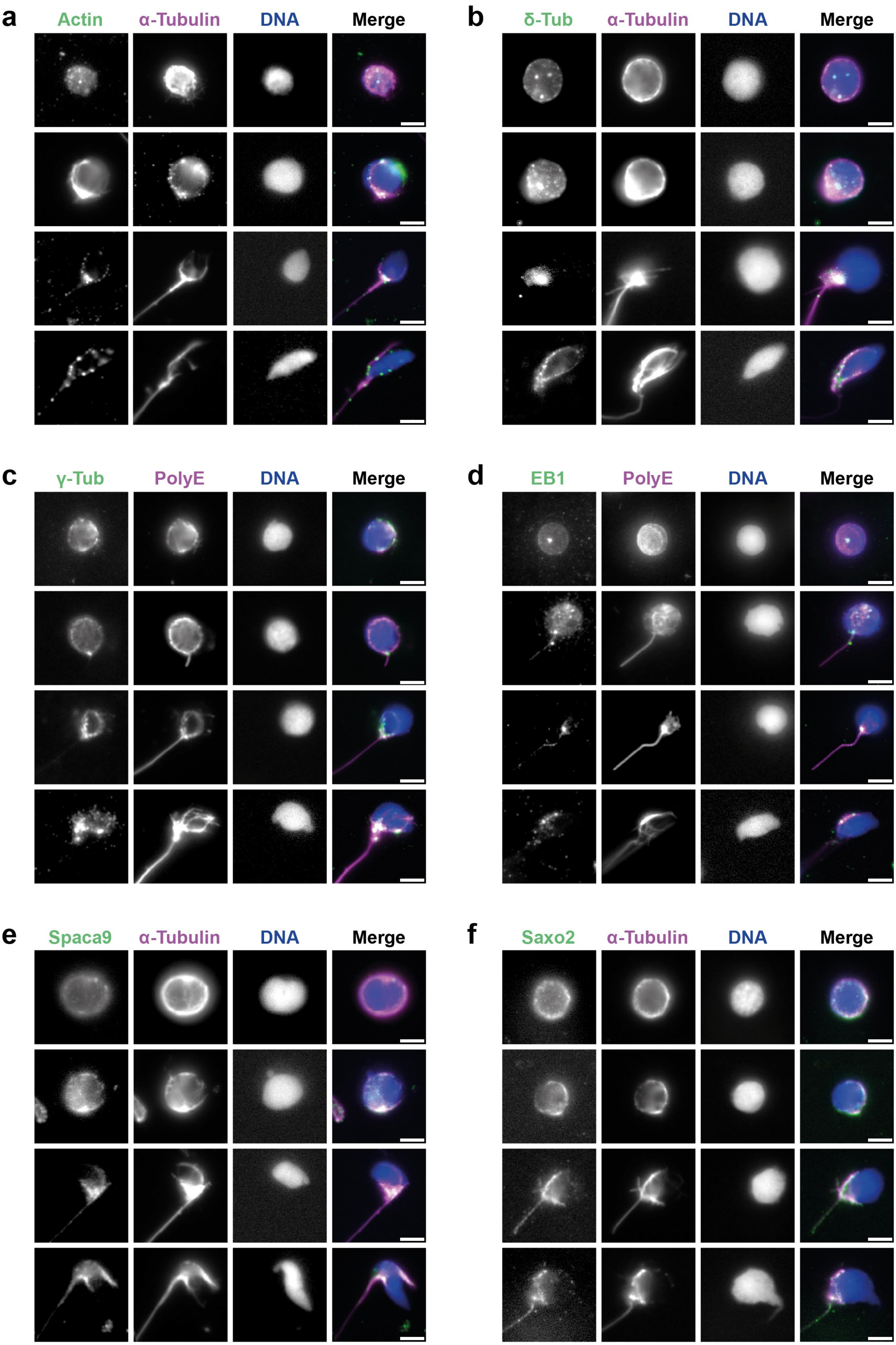
a
Actin
α-Tubulin
DNA
Merge
b
δ-Tub
α-Tubulin
DNA
Merge
c
γ-Tub
PolyE
DNA
Merge
d
EB1
PolyE
DNA
Merge
e
Spaca9
α-Tubulin
DNA
Merge
f
Saxo2
α-Tubulin
DNA
Merge

**Supplementary Fig. 1: Molecular characterization of *Xenopus laevis* spermiogenesis**

**a,** Immunofluorescence images of actin at progressing stages of spermiogenesis. Microtubules (magenta), nuclei (blue) and actin (green) are shown. **b,** Immunofluorescence images of δ-tubulin at progressing stages of spermiogenesis. Microtubules (magenta), nuclei (blue) and δ-tubulin (green) are shown. **c,** Immunofluorescence images of EB1 at progressing stages of spermiogenesis. Microtubules (magenta), nuclei (blue) and EB1 (green) are shown. **d,** Immunofluorescence images of γ-tubulin at progressing stages of spermiogenesis. Microtubules (magenta), nuclei (blue) and γ-tubulin (green) are shown. **e,** Immunofluorescence images of Spaca9 at progressing stages of spermiogenesis. Microtubules (magenta), nuclei (blue) and Spaca9 (green) are shown. **f,** Immunofluorescence images of Saxo2 at progressing stages of spermiogenesis. Microtubules (magenta), nuclei (blue) and Saxo2 (green) are shown. Scale bars: 5 µm.

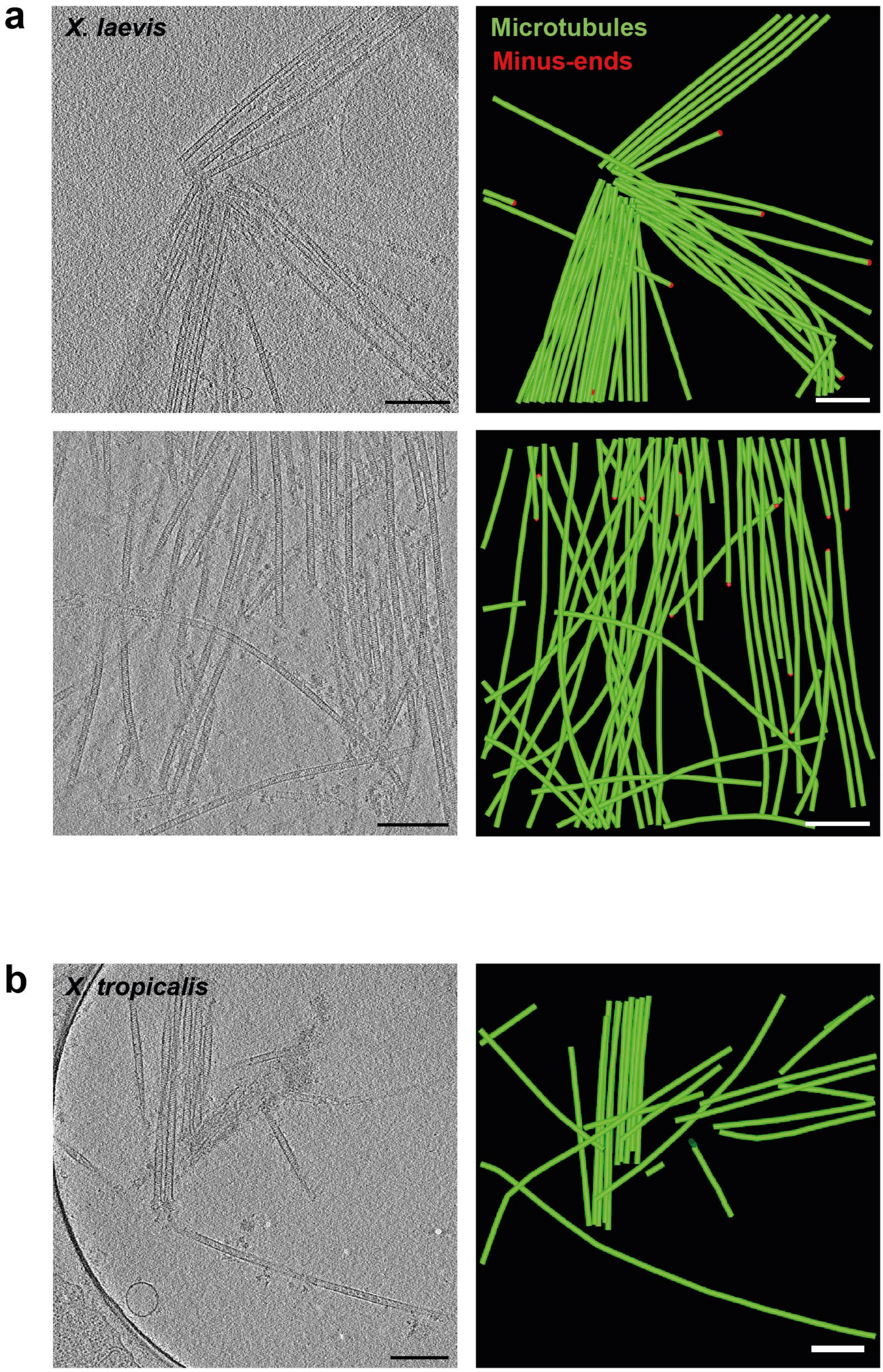
a
X. laevis
Microtubules
Minus-ends
b
X. tropicalis

**Supplementary Fig. 2: Observation of microtubules released from *Xenopus laevis and Xenopus tropicalis* spermatids**

Cryo-electron tomography images of microtubules released from *Xenopus laevis* (**a**,) and *Xenopus tropicalis* (**b**,) spermatids after Triton X-100 treatment (left) and their corresponding models (right). Scale bars: 200 nm.

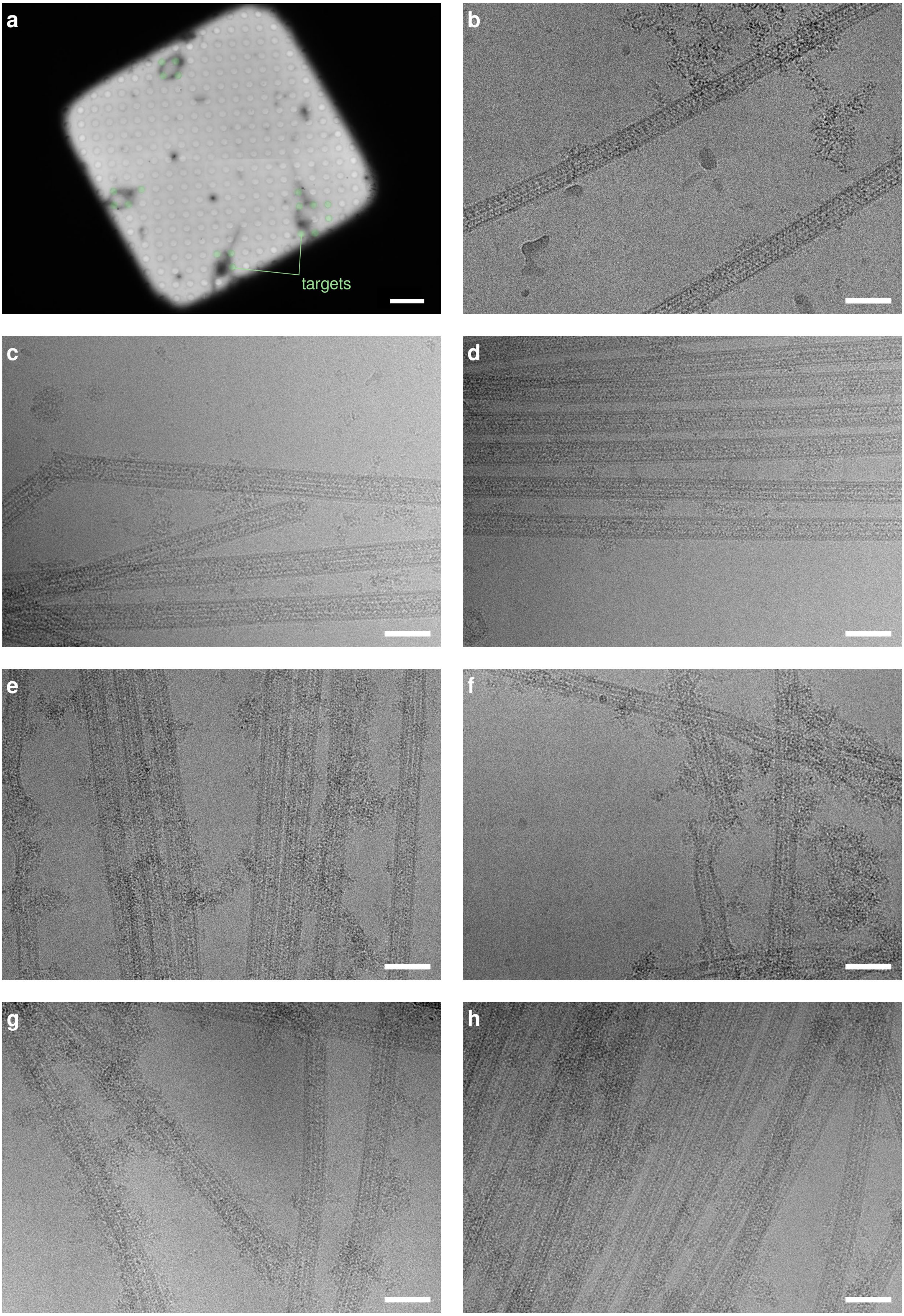
a
b
c
d
e
f
g
h
targets

**Supplementary Fig. 3. High-resolution cryo-EM data collection images of *Xenopus laevis* spermatid microtubules.**

**a**, Example of manual targeting within a grid square. Holes containing microtubules and positioned at the periphery of basket-like structures were manually targeted (green dots). Scale bar: 10 µm. **b-h**, Representative micrographs illustrating the diversity of the imaged microtubules, including variation in diameter, external decoration, conformation and of microtubule density. Scale bars: 50 nm.

1

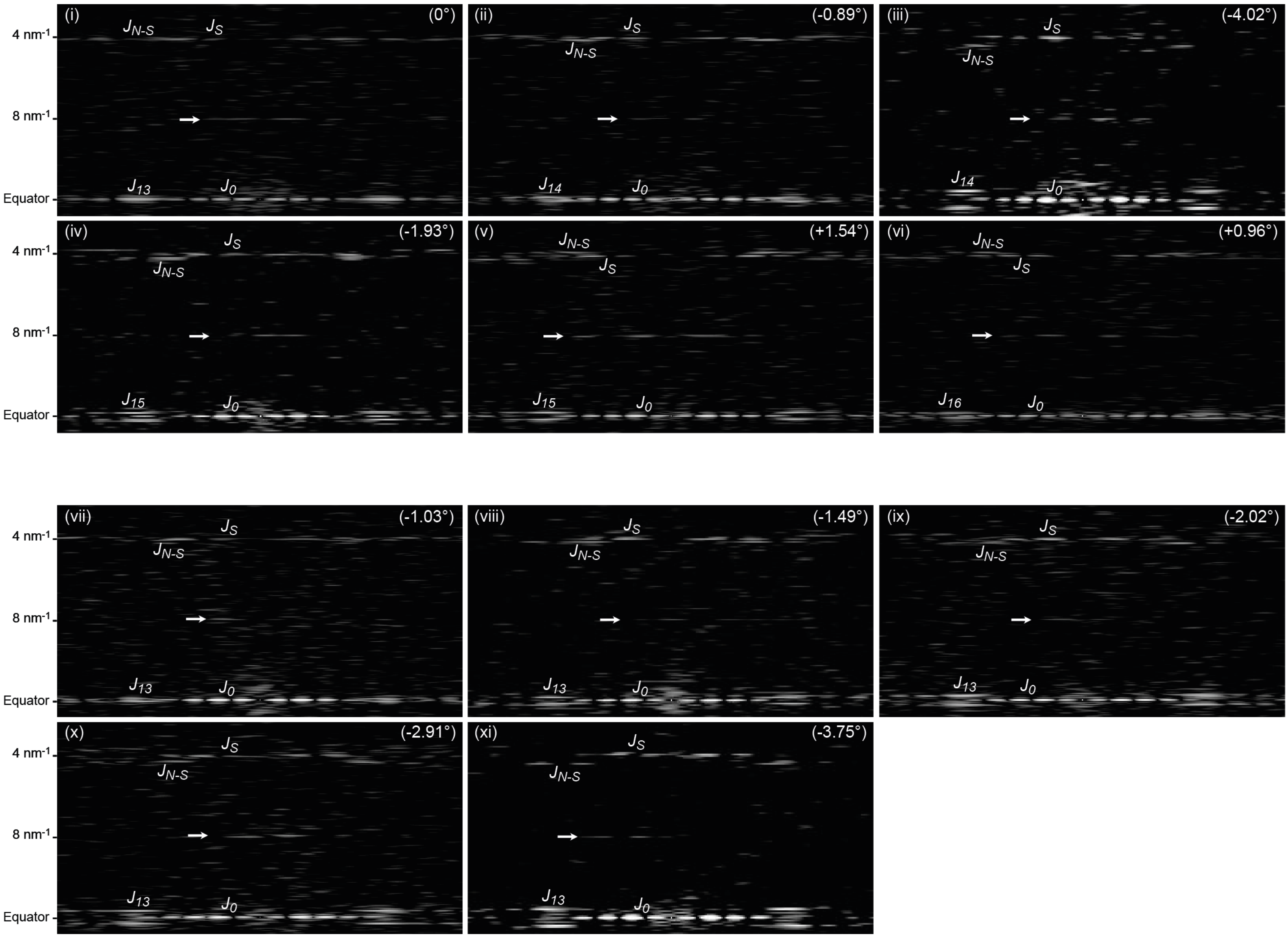

(i)
(0°)
(ii)
(-0.89°)
(iii)
(-4.02°)
(iv)
(-1.93°)
(v)
(+1.54°)
(vi)
(+0.96°)
(vii)
(-1.03°)
(viii)
(-1.49°)
(ix)
(-2.02°)
(x)
(-2.91°)
(xi)
(-3.75°)
4 nm-1
8 nm-1
Equator
$J_{N-S}$
$J_S$
$J_{13}$
$J_{14}$
$J_{15}$
$J_{16}$
$J_0$

**Supplementary Fig. 4: Determination of the sign of the protofilament skew angle from microtubule image Fourier transforms**

Fourier transforms of the microtubule images presented in Fig. 3d,f. In 13 protofilament microtubules with parallel protofilaments ((i), $\theta$ = 0°), the layer lines of order $N$ ($J_{13}$) overlap with that of order 0 ($J_0$) on the equator of the Fourier transforms. Similarly, layer lines located at 4 nm$^{-1}$ reflecting the periodicity of tubulin subunits and their helical arrangement ($J_S$ and $J_{N-S}$) overlap at the same position. For microtubules with skewed protofilaments, $J_N$ is shifted from the equator with a distance related to the amplitude of the protofilament skew angle $\theta$. The sign of $\theta$ can be determined by the relative positions of the $J_S$ and $J_{N-S}$ layer lines. It is negative when $J_S$ is closer to the equator than $J_{N-S}$ (ii-iv, vii-xi), and positive in the alternate case (v, vi). Arrows point the 8 nm$^{-1}$ layer line. The value of the protofilament skew angle is given by the moiré period in filtered images of the microtubules (Fig. 3d,f).

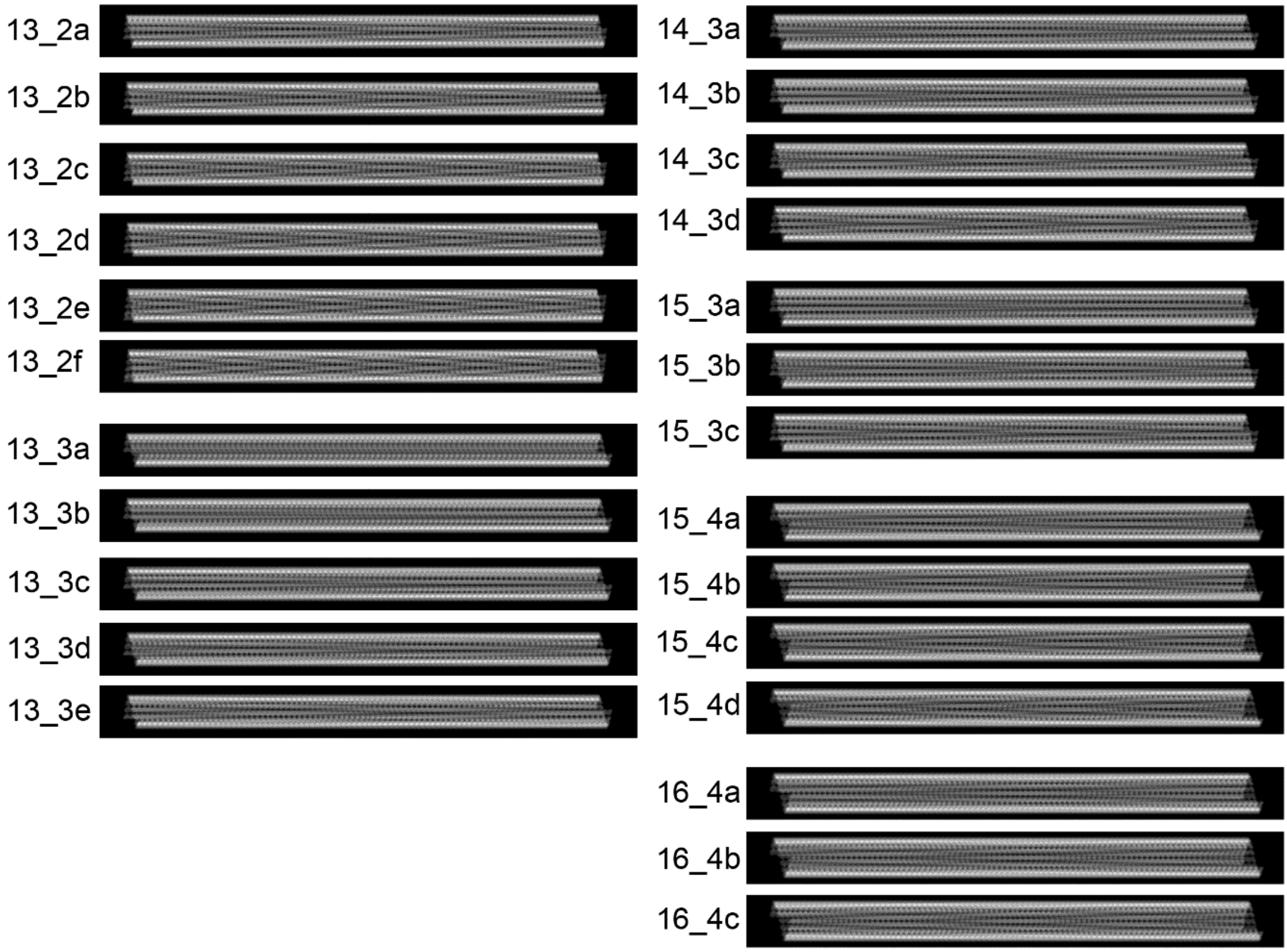
13_2a
13_2b
13_2c
13_2d
13_2e
13_2f
13_3a
13_3b
13_3c
13_3d
13_3e
14_3a
14_3b
14_3c
14_3d
15_3a
15_3b
15_3c
15_4a
15_4b
15_4c
15_4d
16_4a
16_4b
16_4c

**Supplementary Fig. 5:** Projected images of densities generated from the atomic models, after their low-pass filtration to 15 Å resolution with a pixel size of 4 Å, and 1,024 pixels length using parameters listed in Supplementary Table 2.

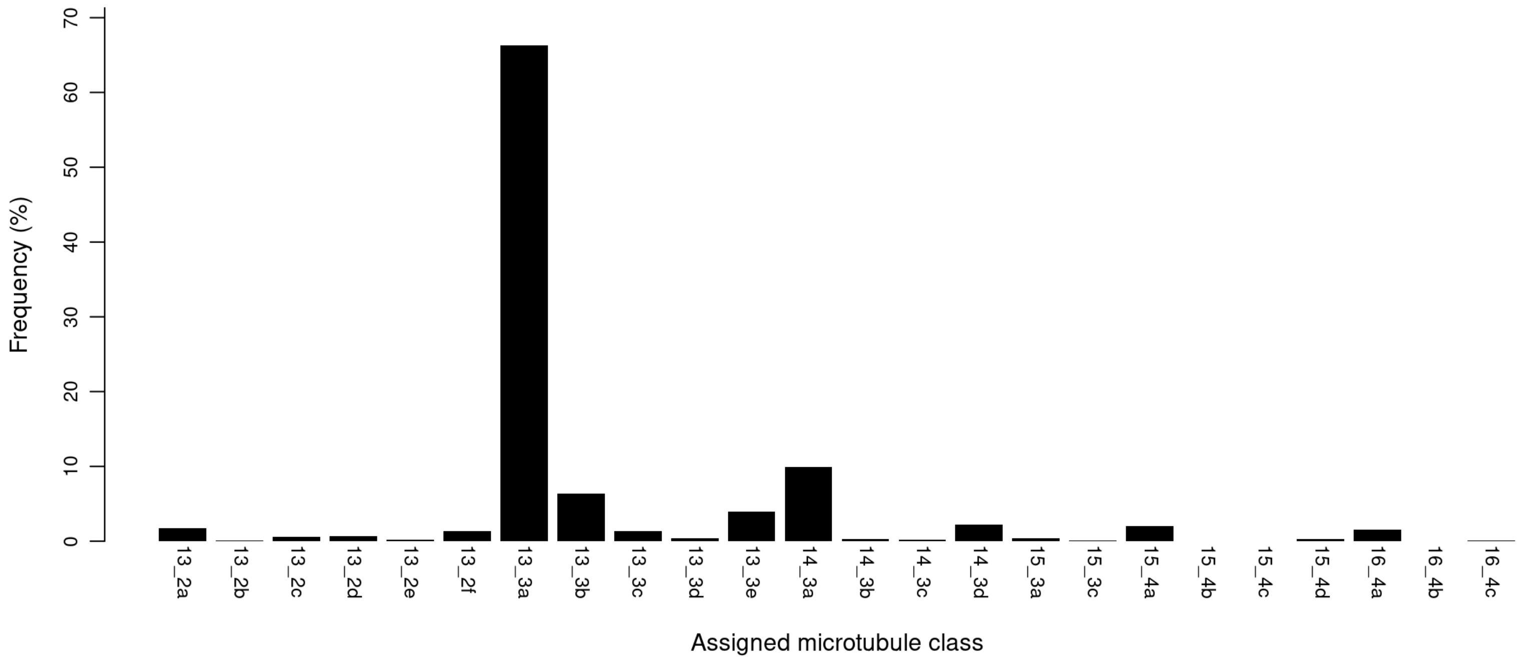


**Supplementary Fig. 6. Reference-based 2D classification of microtubule diversity in the high-resolution cryo-EM dataset of *Xenopus laevis* spermatid microtubules.**
The proportion of microtubule segments assigned to each class is indicated. Classes are defined in Supplementary Table 2.

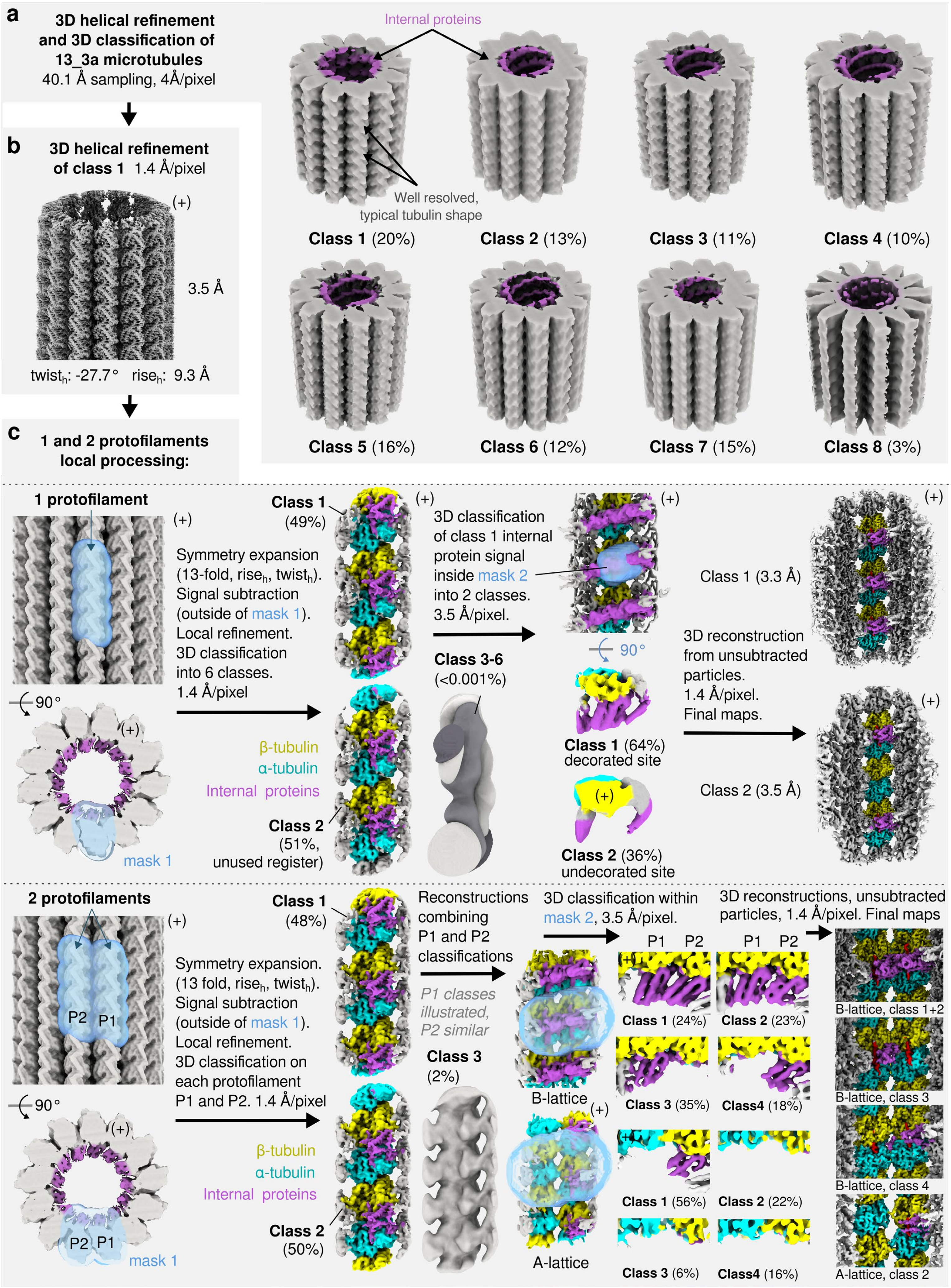
a
3D helical refinement and 3D classification of 13_3a microtubules
40.1 Å sampling, 4Å/pixel
Internal proteins
Well resolved, typical tubulin shape
Class 1 (20%)
Class 2 (13%)
Class 3 (11%)
Class 4 (10%)
Class 5 (16%)
Class 6 (12%)
Class 7 (15%)
Class 8 (3%)
b
3D helical refinement of class 1 1.4 Å/pixel
(+)
3.5 Å
twist$_h$: -27.7° rise$_h$: 9.3 Å
c
1 and 2 protofilaments local processing:
1 protofilament
(+)
90°
mask 1
Symmetry expansion (13-fold, rise$_h$, twist$_h$). Signal subtraction (outside of mask 1). Local refinement. 3D classification into 6 classes. 1.4 Å/pixel
β-tubulin
α-tubulin
Internal proteins
Class 1 (49%)
Class 2 (51%, unused register)
3D classification of class 1 internal protein signal inside mask 2 into 2 classes. 3.5 Å/pixel.
Class 3-6 (<0.001%)
90°
Class 1 (64%) decorated site
Class 2 (36%) undecorated site
3D reconstruction from unsubtracted particles. 1.4 Å/pixel. Final maps.
Class 1 (3.3 Å)
Class 2 (3.5 Å)
2 protofilaments
P2 P1
mask 1
Symmetry expansion. (13 fold, rise$_h$, twist$_h$). Signal subtraction (outside of mask 1). Local refinement. 3D classification on each protofilament P1 and P2. 1.4 Å/pixel
β-tubulin
α-tubulin
Internal proteins
Class 1 (48%)
Class 2 (50%)
Reconstructions combining P1 and P2 classifications
P1 classes illustrated, P2 similar
Class 3 (2%)
B-lattice
A-lattice
3D classification within mask 2, 3.5 Å/pixel.
P1 P2
Class 1 (24%)
Class 2 (23%)
Class 3 (35%)
Class4 (18%)
Class 1 (56%)
Class 2 (22%)
Class 3 (6%)
Class4 (16%)
3D reconstructions, unsubtracted particles, 1.4 Å/pixel. Final maps
B-lattice, class 1+2
B-lattice, class 3
B-lattice, class 4
A-lattice, class 2

**Supplementary Fig. 7. High-resolution cryo-EM processing strategy for reconstruction of the luminal scaffold.**

**a,** Initial helical refinement and 3D classification of the 13_3a microtubule population. **b,** Helical refinement of the selected homogeneous class, including CTF refinement. **c,** Symmetry expansion and focused processing of short one- and two-protofilament segments comprising four tubulin subunits, used to resolve the α/β-tubulin register, luminal protein occupancy, and the B- and A-lattice configurations.

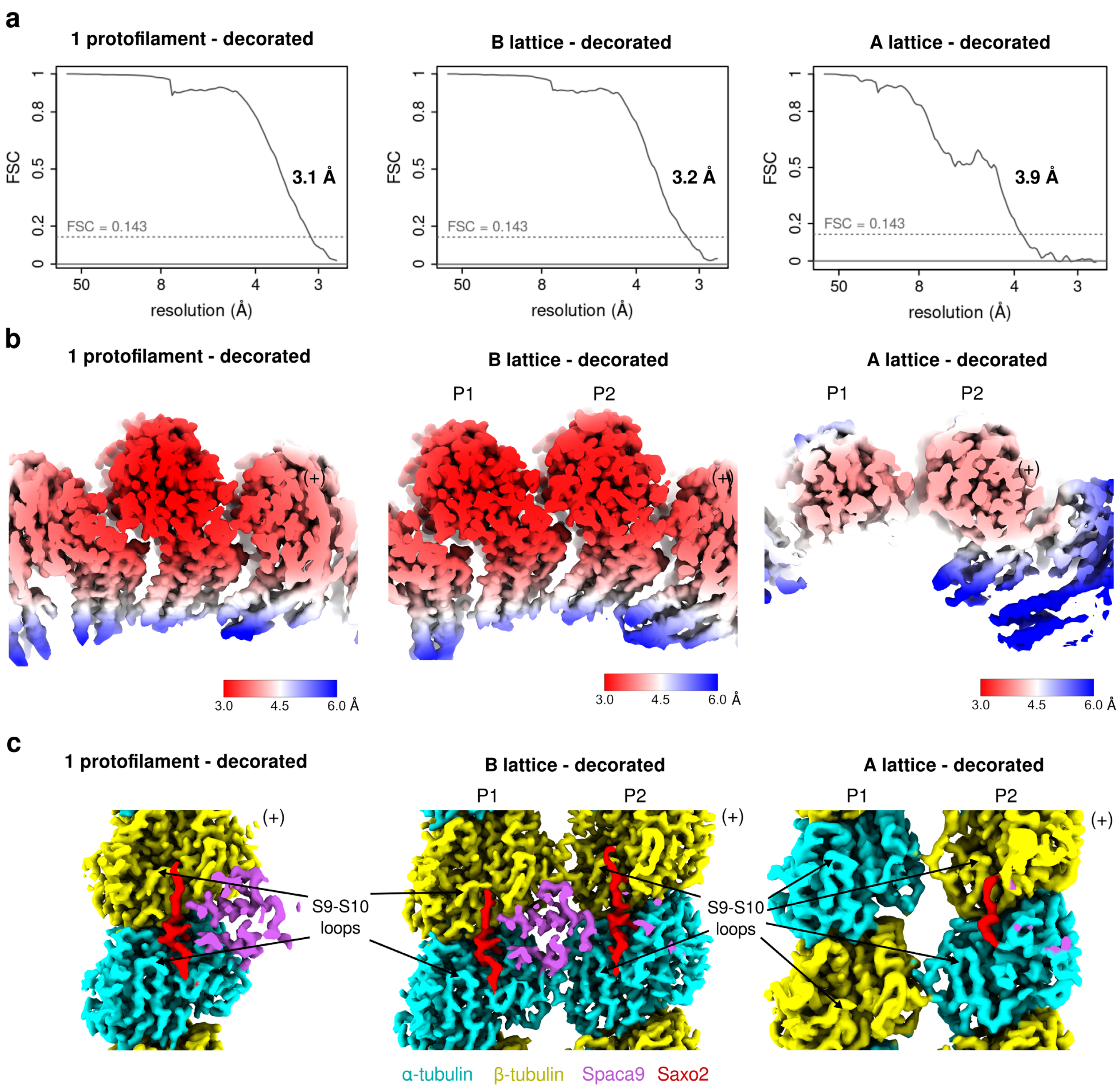

a
1 protofilament - decorated
B lattice - decorated
A lattice - decorated
FSC
resolution (Å)
FSC = 0.143
3.1 Å
3.2 Å
3.9 Å
b
1 protofilament - decorated
B lattice - decorated
A lattice - decorated
P1
P2
(+)
3.0
4.5
6.0 Å
c
1 protofilament - decorated
B lattice - decorated
A lattice - decorated
P1
P2
(+)
S9-S10 loops
α-tubulin
β-tubulin
Spaca9
Saxo2

**Supplementary Fig. 8. Resolution and map quality of the final focused cryo-EM reconstructions of the spermatid microtubule luminal scaffold.**

**a,** Fourier shell correlation (FSC) curves of the final decorated one-protofilament, B-lattice and A-lattice reconstructions, with their corresponding global resolutions indicated. **b,** Local-resolution estimations of the final reconstruction. **c,** Luminal views of the final maps, with the Spaca9 density clipped to better visualize tubulin, illustrating clear separation of the α- and β-tubulin registers.

**a**

| Proteins | iptm_ptm | iQ_score |
|---|---|---|
| saxo2.L | 0.52 | 83.9 |
| hydin.S | 0.72 | 83.1 |
| kif21a.L | 0.75 | 67.4 |
| dlec1.L | 0.63 | 62.4 |
| huwe1.L | Inter-PAE > 10 A | |
| flna.L | Inter-PAE > 10 A | |
| sec16a.L | Inter-PAE > 10 A | |
| prpf8.S | Inter-PAE > 10 A | |
| cfap65.L | Inter-PAE > 10 A | |
| LOC121402910 | Inter-PAE > 10 A | |
| XB5951253.S | Inter-PAE > 10 A | |
| lonp1.L | Inter-PAE > 10 A | |
| copb2.L | Inter-PAE > 10 A | |
| bbs9.L | Inter-PAE > 10 A | |
| plcd4.L | Inter-PAE > 10 A | |
| nop58.L | Inter-PAE > 10 A | |
| XB5765667.L | Inter-PAE > 10 A | |
| saxo4.L | Inter-PAE > 10 A | |
| cnot9.L | Inter-PAE > 10 A | |

**b**

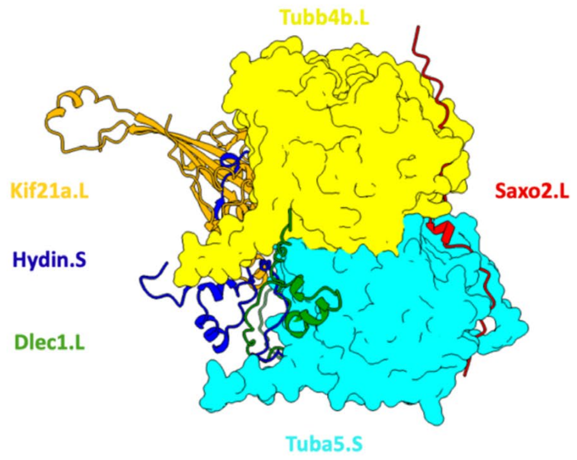


**c**

| Peptides | iptm_ptm | iQ_score |
|---|---|---|
| saxo2.L_242 | 0.87 | 97.5 |
| saxo2.L_377 | 0.89 | 94.7 |
| saxo2.L_141 | 0.81 | 94.3 |
| saxo2.L_343 | 0.88 | 90.3 |
| saxo2.L_107 | 0.85 | 84.6 |
| saxo2.L_74 | 0.85 | 79.8 |
| saxo2.L_309 | 0.86 | 79.6 |
| saxo2.L_209 | 0.88 | 78.6 |
| saxo2.L_175 | 0.86 | 51.2 |
| saxo2.L_17 | Inter-PAE > 10 A | |
| saxo2.L_39 | Inter-PAE > 10 A | |
| saxo2.L_176 | Inter-PAE > 10 A | |
| saxo2.L_276 | Inter-PAE > 10 A | |
| saxo2.L_411 | Inter-PAE > 10 A | |
| saxo2.L_431 | Inter-PAE > 10 A | |
| saxo2.L_453 | Inter-PAE > 10 A | |

**d**

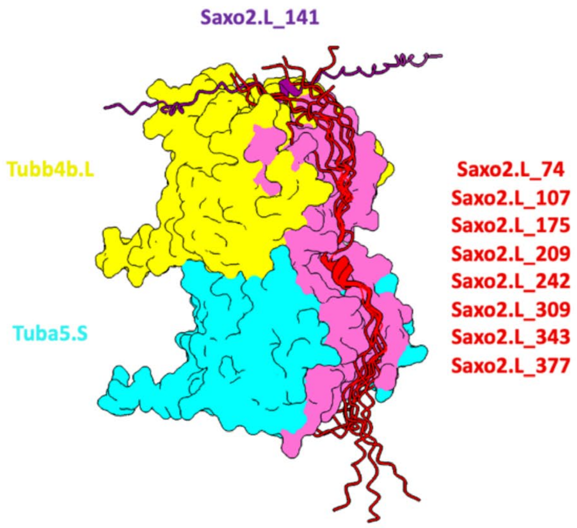

**Supplementary Fig. 9. In silico candidate search for the peptide density in the high-resolution cryo-EM map.**

**a,** Candidate proteins, associated parameters from the AlphaFold3 analysis and iQ-score metric [88] of the 19 proteins selected from the mass spectrometry dataset based on the presence of a matching candidate motif (Methods). The α- and β-tubulin isotypes used for modelling were Tuba5.S (UniProt: Q7ZTP0) and Tubb4b.L (UniProt: P30883), respectively. Inter-chain PAE (predicted aligned error) estimates confidence in the relative positioning of residues from different protein chains, with lower values indicating higher-confidence predicted interfaces. Four proteins passed the interaction-score criteria; however, for three of these, the predicted interaction site was inconsistent with the cryo-EM density because the models did not bind on the luminal face of tubulin. **b**, Predicted structures from the analysis in panel a with an inter-chain PAE ≤ 10 Å, shown at their respective binding sites on tubulin. **c**, AlphaFold3 models of 51-residue Saxo2.L fragments centered on candidate Mn motifs in complex with tubulin. Fragments are named according to the position of their central residue in the Saxo2.L sequence (UniProt: Q6DCB9). Nine fragments were predicted to interact with α- and β-tubulin. **d**, Binding sites of the Saxo2.L fragments from the analysis in panel c with an inter-chain PAE ≤ 10 Å. The interface corresponding to the peptide position observed in the cryo-EM map is highlighted in red. With the exception of Saxo2.L_141, which did not fit the experimental density, the eight retained fragments interacted at the same tubulin interface.

1

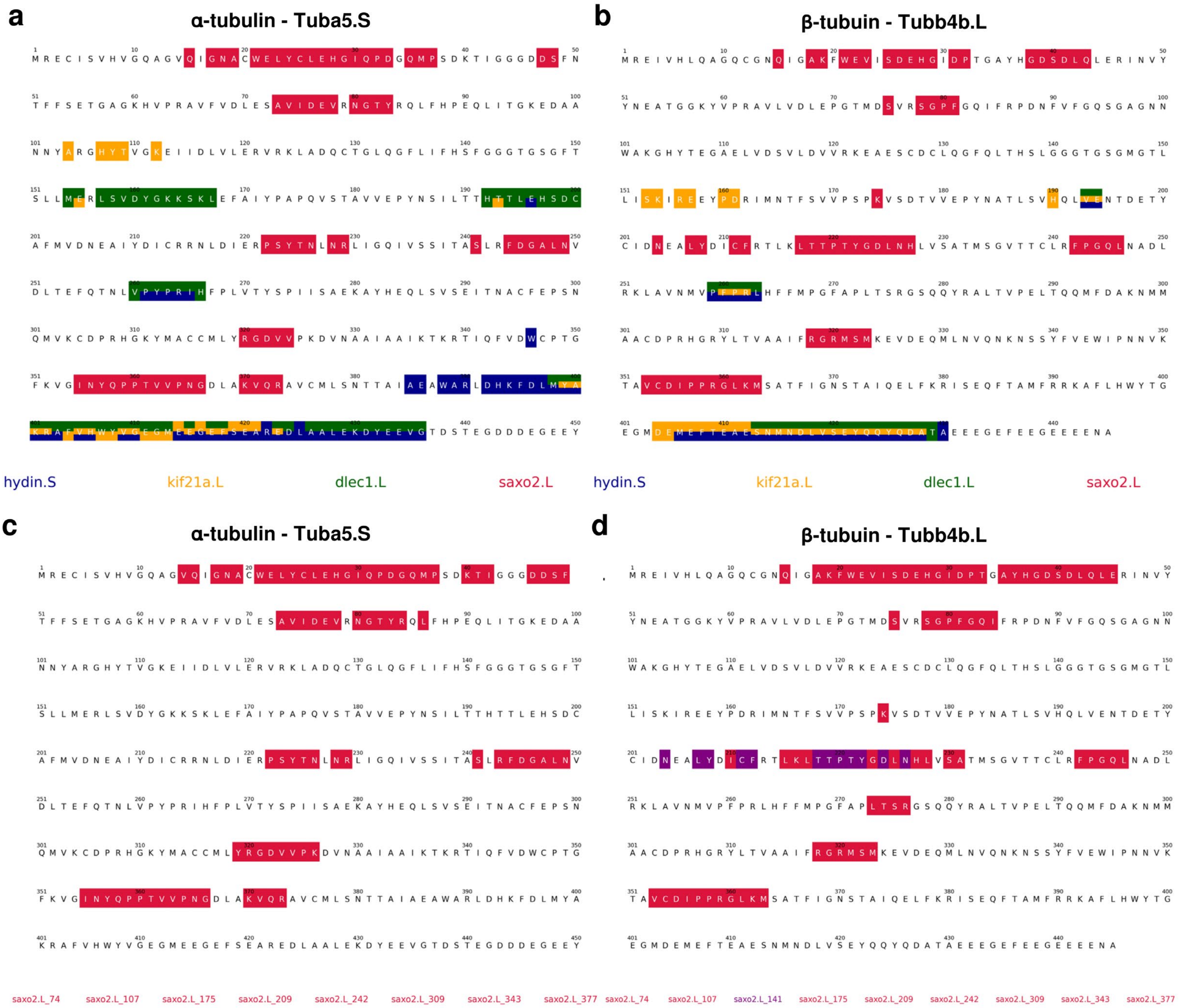
a
α-tubulin - Tuba5.S
b
β-tubuin - Tubb4b.L
hydin.S
kif21a.L
dlec1.L
saxo2.L
hydin.S
kif21a.L
dlec1.L
saxo2.L
c
α-tubulin - Tuba5.S
d
β-tubuin - Tubb4b.L
saxo2.L_74 saxo2.L_107 saxo2.L_175 saxo2.L_209 saxo2.L_242 saxo2.L_309 saxo2.L_343 saxo2.L_377
saxo2.L_74 saxo2.L_107 saxo2.L_141 saxo2.L_175 saxo2.L_209 saxo2.L_242 saxo2.L_309 saxo2.L_343 saxo2.L_377

**Supplementary Fig. 10. Tubulin residues involved in the predicted protein interfaces identified by AlphaFold3 analysis.**

Sequences of α- and β-tubulin are shown with residues at predicted interaction interfaces highlighted according to the interacting protein, using the color code indicated in each panel. The α- and β-tubulin isotypes used for modelling were Tuba5.S (UniProt: Q7ZTP0) and Tubb4b.L (UniProt: P30883), respectively. **a,b,** Interaction residues corresponding to the candidate-protein analysis shown in Supplementary Fig. 9a,b for α- and β-tubulin respectively. **c,d,** Interaction residues corresponding to the Saxo2.L fragment analysis shown in Supplementary Fig. 9c,d, for α- and β-tubulin respectively.

**Supplementary Table 1. Analysis of microtubule configurations**

| *N* | *S* | *θ* | *r* | % |
|---|---|---|---|---|
| 13 | 2 | -1.5 | 7.5152 | 1.74 |
| 13 | 2 | -2.0 | 7.9642 | 0.07 |
| 13 | 2 | -2.5 | 8.4134 | 0.57 |
| 13 | 2 | -3.0 | 8.8630 | 0.72 |
| 13 | 2 | -3.5 | 9.3130 | 0.17 |
| 13 | 2 | -4.0 | 9.7635 | 1.32 |
| 13 | 3 | 0.0 | 9.2538 | 66.31 |
| 13 | 3 | -0.3 | 9.5230 | 6.40 |
| 13 | 3 | -0.7 | 9.8818 | 1.31 |
| 13 | 3 | -1.0 | 10.151 | 0.40 |
| 13 | 3 | -1.3 | 10.420 | 3.91 |

| *N* | *S* | *θ* | *r* | % |
|---|---|---|---|---|
| 14 | 3 | -0.7 | 9.2209 | 9.92 |
| 14 | 3 | -1.0 | 9.4900 | 0.30 |
| 14 | 3 | -1.4 | 9.8490 | 0.23 |
| 14 | 3 | -1.8 | 10.208 | 2.21 |
| 15 | 3 | -0.6 | 8.5583 | 0.38 |
| 15 | 3 | -1.0 | 8.9172 | 0.00 |
| 15 | 3 | -1.4 | 9.2762 | 0.14 |
| 15 | 4 | 1.0 | 9.7961 | 1.98 |
| 15 | 4 | 1.3 | 9.5269 | 0.00 |
| 15 | 4 | 1.6 | 9.2576 | 0.01 |
| 15 | 4 | 1.9 | 8.9882 | 0.29 |
| 16 | 4 | 1.0 | 9.1278 | 1.53 |
| 16 | 4 | 1.3 | 8.8586 | 0.02 |
| 16 | 4 | 1.6 | 8.5893 | 0.06 |

**Supplementary Table 2. Helical parameters of microtubule references used for microtubule classification**

| MT type | *N* | *S* | Theta_deg | r_su_A | Phi_su_deg | Delta_C_A | r_pf_A | Phi_pf_deg |
|---|---|---|---|---|---|---|---|---|
| 13_2a | 13 | 2 | -1.5 | 40.086 | -0.565 | 5.036 | 7.905 | -27.779 |
| 13_2b | 13 | 2 | -2.0 | 40.076 | -0.754 | 5.065 | 7.454 | -27.808 |
| 13_2c | 13 | 2 | -2.5 | 40.062 | -0.941 | 5.101 | 7.003 | -27.837 |
| 13_2d | 13 | 2 | -3.0 | 40.045 | -1.129 | 5.146 | 6.551 | -27.866 |
| 13_2e | 13 | 2 | -3.5 | 40.025 | -1.316 | 5.199 | 6.099 | -27.895 |
| 13_2f | 13 | 2 | -4.0 | 40.002 | -1.503 | 5.260 | 5.646 | -27.924 |
| 13_3a | 13 | 3 | 0.0 | 40.100 | 0.000 | 5.000 | 9.254 | -27.692 |
| 13_3b | 13 | 3 | -0.3 | 40.099 | -0.113 | 5.001 | 8.985 | -27.718 |
| 13_3c | 13 | 3 | -0.7 | 40.097 | -0.264 | 5.008 | 8.625 | -27.753 |
| 13_3d | 13 | 3 | -1.0 | 40.094 | -0.377 | 5.016 | 8.355 | -27.779 |
| 13_3e | 13 | 3 | -1.3 | 40.090 | -0.490 | 5.027 | 8.085 | -27.805 |
| 14_3a | 14 | 3 | -0.7 | 40.097 | -0.245 | 13.189 | 8.625 | -25.767 |
| 14_3b | 14 | 3 | -1.0 | 40.094 | -0.350 | 13.198 | 8.355 | -25.789 |
| 14_3c | 14 | 3 | -1.4 | 40.088 | -0.490 | 13.215 | 7.995 | -25.819 |
| 14_3d | 14 | 3 | -1.8 | 40.080 | -0.630 | 13.237 | 7.635 | -25.849 |
| 15_3a | 15 | 3 | -0.6 | 40.098 | -0.196 | 21.368 | 8.715 | -24.039 |
| 15_3b | 15 | 3 | -1.0 | 40.094 | -0.327 | 21.380 | 8.355 | -24.065 |
| 15_3c | 15 | 3 | -1.4 | 40.088 | -0.457 | 21.398 | 7.995 | -24.091 |
| 15_4a | 15 | 4 | +1.0 | 40.094 | 0.327 | 21.380 | 10.149 | -23.913 |
| 15_4b | 15 | 4 | +1.3 | 40.090 | 0.425 | 21.393 | 10.418 | -23.887 |
| 15_4c | 15 | 4 | +1.6 | 40.084 | 0.523 | 21.409 | 10.685 | -23.861 |
| 15_4d | 15 | 4 | +1.9 | 40.078 | 0.620 | 21.429 | 10.953 | -23.835 |
| 16_4a | 16 | 4 | +1.0 | 40.094 | 0.306 | 29.562 | 10.149 | -22.423 |
| 16_4b | 16 | 4 | +1.3 | 40.090 | 0.398 | 29.575 | 10.418 | -22.400 |
| 16_4c | 16 | 4 | +1.6 | 40.084 | 0.490 | 29.593 | 10.685 | -22.378 |

**Supplementary Table 3. Cryo-EM data collection, refinement and validation statistics.**

| | **1 protofilament-decorated** (EMDB-59871) (PDB 34IK) | **B-lattice-decorated** (EMDB-59872) (PDB 34IL) | **A-lattice-decorated** (EMDB-59873) (PDB 34IM) |
|---|---|---|---|
| **Data collection and processing** | | | |
| Magnification (nominal) | 105,000 | 105,000 | 105,000 |
| Voltage (kV) | 300 | 300 | 300 |
| Electron exposure ($e^{-}$/Å$^2$) | 50 | 50 | 50 |
| Defocus range (µm) [a] | 0.8 ; 2.4 | 0.8 ; 2.4 | 0.8 ; 2.5 |
| Pixel size collection (Å)[b] | 0.832 | 0.832 | 0.832 |
| Final refinement pixel size (Å) | 1.4 | 1.4 | 1.4 |
| Symmetry imposed [c] | Helical | Helical | Helical |
| Rise (Å) | 9.3 | 9.3 | 9.3 |
| Twist (deg) | -27.7 | -27.7 | -27.7 |
| Particle images identified as 13_3a symmetry (no.), 40.1 A spacing | 208,920 | 208,920 | 208,920 |
| Particle images in last helical reconstruction (no.), 40.1 A spacing | 42,581 | 42,581 | 42,581 |
| Single particles – tubulin dimers after expansion (no.) | 276,776 | 276,776 | 276,776 |
| Single particles after classifications (no.)[d] | 175,220 | 108,762 | 16,603 |
| Overall resolution (Å) | 3.1 | 3.2 | 4.0 |
| FSC threshold | 0.143 | 0.143 | 0.143 |
| **Refinement** | | | |
| Model composition | | | |
| Non-hydrogen atoms | 14,931 | 29,856 | 28,394 |
| Protein residues | 1,884 | 3,767 | 3,582 |
| Ligands | 6 | 12 | 12 |
| R.m.s. deviations | | | |
| Bond lengths (Å) | 0.0053 | 0.0071 | 0.0073 |
| Bond angles (°) | 1.23 | 1.39 | 1.66 |
| Validation | | | |
| MolProbity score | 1.59 | 1.67 | 2.10 |
| Clashscore | 4.37 | 5.10 | 9.56 |
| Poor rotamers (%) | 0.41 | 0.65 | 0.98 |
| Ramachandran plot | | | |
| Favored (%) | 94.56 | 94.06 | 88.29 |
| Allowed (%) | 5.09 | 5.60 | 11.00 |
| Disallowed (%) | 0.36 | 0.35 | 0.70 |

[a] Range of the per-particle defocus estimates. The range corresponds to 90% of the particles used, 5% of the particle defocuses are below and 5% are above this range.

[b] Pixel size after magnification anisotropy correction (see methods).

[c] Symmetry imposed during the symmetrical refinement done before the local refinements (rise and twists provided for helical symmetry only).

[d] Number of asymmetric units, corresponding to a tubulin dimer decorated with internal proteins.

**Supplementary Table 4. List of the 40 most abundant proteins detected by mass spectrometry ranked according to their emPAI score.**

The most abundant β-tubulin (Tubb4b.L) is highlighted in yellow, the most abundant α-tubulin (Tuba5.S) in cyan, Spaca9.L in purple and Saxo2.L in red. Two mass spectrometry analyses are shown.

<table>
<tr><th colspan="2">Replicate 1</th></tr>
<tr><th>Description</th><th>emPAI</th></tr>
<tr><td>XBXL10_1g34175|XBmRNA64463| tubb4b.L</td><td>325.67</td></tr>
<tr><td>XBXL10_1g37468|XBmRNA70732| tubb4b.S</td><td>299.39</td></tr>
<tr><td>XBXL10_1g27580|XBmRNA51753| tubb2b.L</td><td>164.11</td></tr>
<tr><td>XBXL10_1g35505|XBmRNA67050| tubb.L</td><td>130.21</td></tr>
<tr><td>XBXL10_1g17216|XBmRNA31918| tubb4a.S</td><td>78.22</td></tr>
<tr><td>XBXL10_1g14287|XBmRNA26952| tubb4a.L</td><td>78.22</td></tr>
<tr><td>XBXL10_1g29289|XBmRNA55038| tubb2b.S</td><td>71.83</td></tr>
<tr><td>XBXL10_1g9867|XBmRNA18437| tuba5.S</td><td>34.74</td></tr>
<tr><td>XBXL10_1g8013|XBmRNA15151| tuba1c.L</td><td>32.39</td></tr>
<tr><td>XBXL10_1g10680|XBmRNA20062| tuba1c.S</td><td>32.39</td></tr>
<tr><td>XBXL10_1g6992|XBmRNA13089| tuba5.L</td><td>29.49</td></tr>
<tr><td>XBXL10_1g28819|XBmRNA54220| vim.S</td><td>28.73</td></tr>
<tr><td>XBXL10_1g8015|XBmRNA15154| prph.L</td><td>26.37</td></tr>
<tr><td>XBXL10_1g40524|XBmRNA76244| tuba1cl.3.L</td><td>22.42</td></tr>
<tr><td>XBXL10_1g8280|XBmRNA15660| krt8.1.L</td><td>22.22</td></tr>
<tr><td>XBXL10_1g18435|XBmRNA33885| banf1.L</td><td>22.06</td></tr>
<tr><td>XBXL10_1g9152|XBmRNA17042| cbr1.1.S</td><td>19.59</td></tr>
<tr><td>XBXL10_1g12927|XBmRNA24474| cimap1a.L</td><td>19.51</td></tr>
<tr><td>XBXL10_1g16596|XBmRNA31183| LOC121402223</td><td>18.64</td></tr>
<tr><td>XBXL10_1g43356|XBmRNA81520| tuba4b.S</td><td>18.58</td></tr>
<tr><td>XBXL10_1g10869|XBmRNA20449| krt8.1.S</td><td>17.87</td></tr>
<tr><td>XBXL10_1g34385|XBmRNA64893| spaca9.L</td><td>17.66</td></tr>
<tr><td>XBXL10_1g40515|XBmRNA76227| tuba4b.L</td><td>17.61</td></tr>
<tr><td>XBXL10_1g41034|XBmRNA77414| hba1.L</td><td>17.05</td></tr>
<tr><td>XBXL10_1g15973|XBmRNA29852| cimap1a.S</td><td>16.43</td></tr>
<tr><td>XBXL10_1g27774|XBmRNA52130| tubb6.L</td><td>16.12</td></tr>
<tr><td>XBXL10_1g29461|XBmRNA55404| tubb6.S</td><td>16.12</td></tr>
<tr><td>XBXL10_1g15315|XBmRNA28601| LOC108712690</td><td>15.75</td></tr>
<tr><td>XBXL10_1g26898|XBmRNA50375| vim.L</td><td>15.37</td></tr>
<tr><td>XBXL10_1g22782|XBmRNA42459| LOC108716546</td><td>13.58</td></tr>
<tr><td>XBXL10_1g38937|XBmRNA73489| krt19.L</td><td>12.69</td></tr>
<tr><td>XBXL10_1g2934|XBmRNA5247| lmnb1.L</td><td>12.49</td></tr>
<tr><td>XBXL10_1g34174|XBmRNA64462| LOC108698120</td><td>12.38</td></tr>
<tr><td>XBXL10_1g10681|XBmRNA20066| prph.S</td><td>12.16</td></tr>
<tr><td>XBXL10_1g13257|XBmRNA25137| saxo2.L</td><td>11.82</td></tr>
<tr><td>XBXL10_1g42192|XBmRNA79320| krt19.S</td><td>11.17</td></tr>
<tr><td>XBXL10_1g10899|XBmRNA20491| cab39l.S</td><td>10.67</td></tr>
<tr><td>XBXL10_1g18772|XBmRNA34612| hsbp1.L</td><td>9.75</td></tr>
<tr><td>XBXL10_1g29638|XBmRNA55792| zc2hc1a.S</td><td>9.71</td></tr>
<tr><td>XBXL10_1g2818|XBmRNA5010| ccdc62.L</td><td>8.78</td></tr>
</table>

<table>
<tr><th colspan="2">Replicate 2</th></tr>
<tr><th>Description</th><th>EmPAI</th></tr>
<tr><td>XBXL10_1g34175|XBmRNA64463| tubb4b.L</td><td>325.67</td></tr>
<tr><td>XBXL10_1g37468|XBmRNA70732| tubb4b.S</td><td>299.39</td></tr>
<tr><td>XBXL10_1g27580|XBmRNA51753| tubb2b.L</td><td>164.11</td></tr>
<tr><td>XBXL10_1g35505|XBmRNA67050| tubb.L</td><td>130.21</td></tr>
<tr><td>XBXL10_1g14287|XBmRNA26952| tubb4a.L</td><td>78.22</td></tr>
<tr><td>XBXL10_1g17216|XBmRNA31918| tubb4a.S</td><td>78.22</td></tr>
<tr><td>XBXL10_1g29289|XBmRNA55038| tubb2b.S</td><td>71.83</td></tr>
<tr><td>XBXL10_1g9867|XBmRNA18437| tuba5.S</td><td>34.74</td></tr>
<tr><td>XBXL10_1g43357|XBmRNA81521| tuba1cl.2.S</td><td>34.46</td></tr>
<tr><td>XBXL10_1g10680|XBmRNA20062| tuba1c.S</td><td>32.39</td></tr>
<tr><td>XBXL10_1g8013|XBmRNA15151| tuba1c.L</td><td>32.39</td></tr>
<tr><td>XBXL10_1g18724|XBmRNA34516| tubb3.L</td><td>31.64</td></tr>
<tr><td>XBXL10_1g6992|XBmRNA13089| tuba5.L</td><td>29.49</td></tr>
<tr><td>XBXL10_1g28819|XBmRNA54220| vim.S</td><td>28.73</td></tr>
<tr><td>XBXL10_1g8015|XBmRNA15154| prph.L</td><td>26.37</td></tr>
<tr><td>XBXL10_1g40524|XBmRNA76244| tuba1cl.3.L</td><td>22.42</td></tr>
<tr><td>XBXL10_1g8280|XBmRNA15660| krt8.1.L</td><td>22.22</td></tr>
<tr><td>XBXL10_1g18435|XBmRNA33885| banf1.L</td><td>22.06</td></tr>
<tr><td>XBXL10_1g9152|XBmRNA17042| cbr1.1.S</td><td>19.59</td></tr>
<tr><td>XBXL10_1g12927|XBmRNA24474| cimap1a.L</td><td>19.51</td></tr>
<tr><td>XBXL10_1g16596|XBmRNA31183| LOC121402223</td><td>18.64</td></tr>
<tr><td>XBXL10_1g43356|XBmRNA81520| tuba4b.S</td><td>18.58</td></tr>
<tr><td>XBXL10_1g10869|XBmRNA20449| krt8.1.S</td><td>17.87</td></tr>
<tr><td>XBXL10_1g34385|XBmRNA64893| spaca9.L</td><td>17.66</td></tr>
<tr><td>XBXL10_1g40515|XBmRNA76227| tuba4b.L</td><td>17.61</td></tr>
<tr><td>XBXL10_1g41034|XBmRNA77414| hba1.L</td><td>17.05</td></tr>
<tr><td>XBXL10_1g15973|XBmRNA29852| cimap1a.S</td><td>16.43</td></tr>
<tr><td>XBXL10_1g27774|XBmRNA52130| tubb6.L</td><td>16.12</td></tr>
<tr><td>XBXL10_1g29461|XBmRNA55404| tubb6.S</td><td>16.12</td></tr>
<tr><td>XBXL10_1g15315|XBmRNA28601| LOC108712690</td><td>15.75</td></tr>
<tr><td>XBXL10_1g26898|XBmRNA50375| vim.L</td><td>15.37</td></tr>
<tr><td>XBXL10_1g22782|XBmRNA42459| LOC108716546</td><td>13.58</td></tr>
<tr><td>XBXL10_1g43765|XBmRNA82374| hba1.S</td><td>12.88</td></tr>
<tr><td>XBXL10_1g38937|XBmRNA73489| krt19.L</td><td>12.69</td></tr>
<tr><td>XBXL10_1g2934|XBmRNA5247| lmnb1.L</td><td>12.49</td></tr>
<tr><td>XBXL10_1g34174|XBmRNA64462| LOC108698120</td><td>12.38</td></tr>
<tr><td>XBXL10_1g10681|XBmRNA20066| prph.S</td><td>12.16</td></tr>
<tr><td>XBXL10_1g13257|XBmRNA25137| saxo2.L</td><td>11.82</td></tr>
<tr><td>XBXL10_1g42192|XBmRNA79320| krt19.S</td><td>11.17</td></tr>
<tr><td>XBXL10_1g10899|XBmRNA20491| cab39l.S</td><td>10.67</td></tr>
</table>

**Supplementary Table 5. Comparison of Saxo2.L and Saxo4.L mass spectrometry results**

Two mass spectrometry analyses are shown.

| | Number of residues | Specific Spectral Count | EmPAI |
|---|---|---|---|
| **Replicate 1** | | | |
| XBXL10_1g13257\|XBmRNA25137\| saxo2.L | 469 | 226 | 11.82 |
| XBXL10_1g17896\|XBmRNA33058\| saxo4.L | 384 | 7 | 3.44 |
| **Replicate 2** | | | |
| XBXL10_1g13257\|XBmRNA25137\| saxo2.L | 469 | 138 | 11.82 |
| XBXL10_1g17896\|XBmRNA33058\| saxo4.L | 384 | 12 | 3.44 |
